# A quantitative gibberellin signalling biosensor reveals a role for gibberellins in internode specification at the shoot apical meristem

**DOI:** 10.1101/2021.06.11.448154

**Authors:** Bihai Shi, Amelia Felipo-Benavent, Guillaume Cerutti, Carlos Galvan-Ampudia, Lucas Jilli, Geraldine Brunoud, Jérome Mutterer, Elody Vallet, Lali Sakvarelidze-Achard, Jean-Michel Davière, Alejandro Navarro-Galiano, Ankit Walia, Shani Lazary, Jonathan Legrand, Roy Weinstain, Alexander M. Jones, Salomé Prat, Patrick Achard, Teva Vernoux

**Affiliations:** College of Agriculture, South China Agricultural University, Guangdong Laboratory for Lingnan Modern Agriculture, 510642, Guangzhou, China; Laboratoire Reproduction et Développement des Plantes, Univ Lyon, ENS de Lyon, CNRS, INRAE, INRIA, F-69342, Lyon, France; Institut de biologie moléculaire des plantes, CNRS, Université de Strasbourg, 67084 Strasbourg, France; Centre for Research in Agricultural Genomics, 08193 Cerdanyola, Barcelona, Spain; Sainsbury Laboratory, Cambridge University, Cambridge CB2 1LR, United Kingdom; Department of Molecular Biology and Ecology of Plants, Tel Aviv University, Tel Aviv 69978, Israel

## Abstract

Growth at the shoot apical meristem (SAM) is essential for shoot architecture construction. The phytohormones gibberellins (GA) play a pivotal role in coordinating plant growth, but their role in the SAM remains mostly unknown. Here, we developed a ratiometric GA signalling biosensor by engineering one of the DELLA proteins, to suppress its master regulatory function in GA transcriptional responses while preserving its degradation upon GA sensing. We demonstrate that this novel degradation-based biosensor accurately reports on cellular changes in GA levels and perception during development. We used this biosensor to map GA signalling activity in the SAM. We show that high GA signalling is found primarily in cells located between organ primordia that are the precursors of internodes. By gain- and loss-of-function approaches, we further demonstrate that GAs regulate cell division plane orientation to establish the typical cellular organisation of internodes, thus contributing to internode specification in the SAM.

## Introduction

The shoot apical meristem (SAM) at the tip of shoot axes comprises a stem cell niche whose activity produces lateral organs and stem segments in a modular iterative fashion during the whole plant life. Each of these repetitive units or phytomere includes an internode and lateral organs at a node and an axillary meristem at the leaf axil^1^. The growth and organization of phytomeres change during development. In *Arabidopsis thaliana*, internode growth is inhibited during the vegetative phase and axillary meristems rest dormant at the axils of rosette leaves. Upon floral transition, the SAM turns into an inflorescence meristem, producing elongated internodes and axillary buds that form branches at the axils of cauline leaves, and later flowers without leaves^2^. While substantial progress has been made on our understanding of the mechanisms controlling the initiation of leaves, flowers and branches, much less is known on how internodes are initiated.

Growth via cell division and expansion is essential for reiterative organogenesis at the SAM. The tetracyclic diterpenoid hormones gibberellins (GA) are key growth regulators^3–5^, with a crucial role in many embryonic and post-embryonic developmental processes^6^. Central to the GA signalling pathway are the five DELLA proteins, GIBBERELLIC ACID INSENSITIVE (GAI), REPRESSOR OF GA1-3 (RGA) and RGA-Like (RGL) 1-3^7^. These nuclear proteins are composed of an N-terminal DELLA/TVHYNP domain and of a GRAS domain. The GRAS domain allows DELLAs to interact with diverse transcription factors and transcriptional regulators and to suppress growth by modulating their activity^7^. Binding of GA to the GIBBERELLIN INSENSITIVE DWARF1 (GID1) GA receptor promotes GID1 interaction with the N-terminal domain of DELLAs, triggering DELLA degradation by the ubiquitin-dependent proteasome pathway^7–10^, and in turn, de-repressing GA responses. Despite the relatively specialised role of the GRAS and DELLA/TVHYNP domains, residues required for DELLA degradation and partner protein interaction are widely distributed within the protein sequence.

The identification of genes encoding GA catabolic enzymes as direct targets of the class I KNOX meristem identity regulators has led to propose that GA levels are low in the SAM cells, while high GA concentrations trigger growth of lateral organs^3–5^. Low GA levels could then contribute to SAM maintenance^3^. However, more recent analysis indicate also that GAs promote the increase in SAM size during floral transition by regulating the division and expansion of inner SAM cells^11, 12^, in a similar manner as in the root^13, 14^. DELLAs were likewise found to limit meristem size by directly regulating the expression of the cell-cycle inhibitor KRP2 in the internal part of the SAM (the rib zone)^12^. Together, these findings support a role for GA in positively regulating cell division in the inner tissues of the SAM, and thus SAM size. At the same time, several genes encoding GA biosynthetic and catabolic enzymes are expressed specifically in lateral organs^4, 11^, illustrating a complex spatio-temporal GA distribution in the SAM to likely fulfil different functions.

Accessing spatio-temporal GA distribution has been instrumental to better understand functions of these hormones in different tissues and at various developmental stages. Accumulation of GA in the root endodermis and regulation of their cellular level via GA transport were discovered by using bioactive fluorescein (Fl)-tagged GA^15, 16^. More recently, the nlsGPS1 GA FRET sensor revealed that GA levels are correlated with cell elongation in roots, stamen filaments and dark-grown hypocotyls^17^. Here, building on the knowledge on the GA signalling pathway, we report on the engineering and characterization of a quantitative degradation-based GA signalling biosensor. We used this novel biosensor to map the spatio- temporal distribution of GA signalling activity and to quantitatively analyse how GAs regulate cell behaviour in the SAM epidermis. We demonstrate that GAs regulate the orientation of division planes of SAM cells located between organ primordia, therefore specifying the typical cellular organization of internodes.

## Results

### **Modifying the RGA protein for sensor construction** (Fig. 1; Extended Data Fig. 1; Supplementary Table 1)

To generate a degradation-based GA signalling biosensor, we engineered a DELLA protein to meet two criteria, (i) a specific degradation upon GA perception; and (ii) a minimal interference with GA signalling. To do so, we modified DELLAs to preserve the interaction with GID1 while abolishing interactions with partner transcription factors, by introducing mutations in the GRAS domain. Among the 5 *Arabidopsis* DELLAs, RGA displays one of the highest GA-dependent degradation rate^18^, and RGA-GFP fusions are widely used as GA signalling reporter^8^. Leveraging on the results of GRAS domain mutant analyses in rice^19^ and *Arabidopsis*^20, 21^, we generated four modified RGA versions (RGA^m^^1^ to RGA^m^^4^) and tested their ability to meet the above defined criteria (Fig. 1a, Extended Data Fig. 1a). We first used a yeast-two hybrid (Y2H) assay to test the binding capacity of these modified candidates with three well known DELLA interacting partners, JAZ1, TCP14 and IDD2^22^. While RGA^m^^1^ had a minor effect on interactions, RGA^m^^2^, RGA^m^^3^ and RGA^m4^ lost their capacity to interact with these partners (Fig. 1b, Extended Data Fig. 1b). In addition, RGA^m2^, RGA^m3^ and RGA^m4^ were able to bind GID1 in the presence of GA, thus suggesting that these DELLA candidates are still degraded in response to GA (Fig. 1c, Extended Data Fig. 1c). To assess this possibility, we explored GA-dependent degradation of RGA^m2^, RGA^m3^ and RGA^m4^ fused to GFP in transient expression assays. These three candidates were degraded after GA treatment (Fig. 1d, Extended Data Fig. 1d), RGA^m2^-GFP having the fastest degradation kinetics although it was slightly more stable than RGA-GFP. Noteworthy, RGA^m2^ harbours three amino acid substitutions in the GRAS PFYRE motif (Fig. 1a), that is highly conserved in all plant DELLAs. Therefore, this variant is likely to have similar properties in plant species other than *Arabidopsis*.

**Figure 1.**
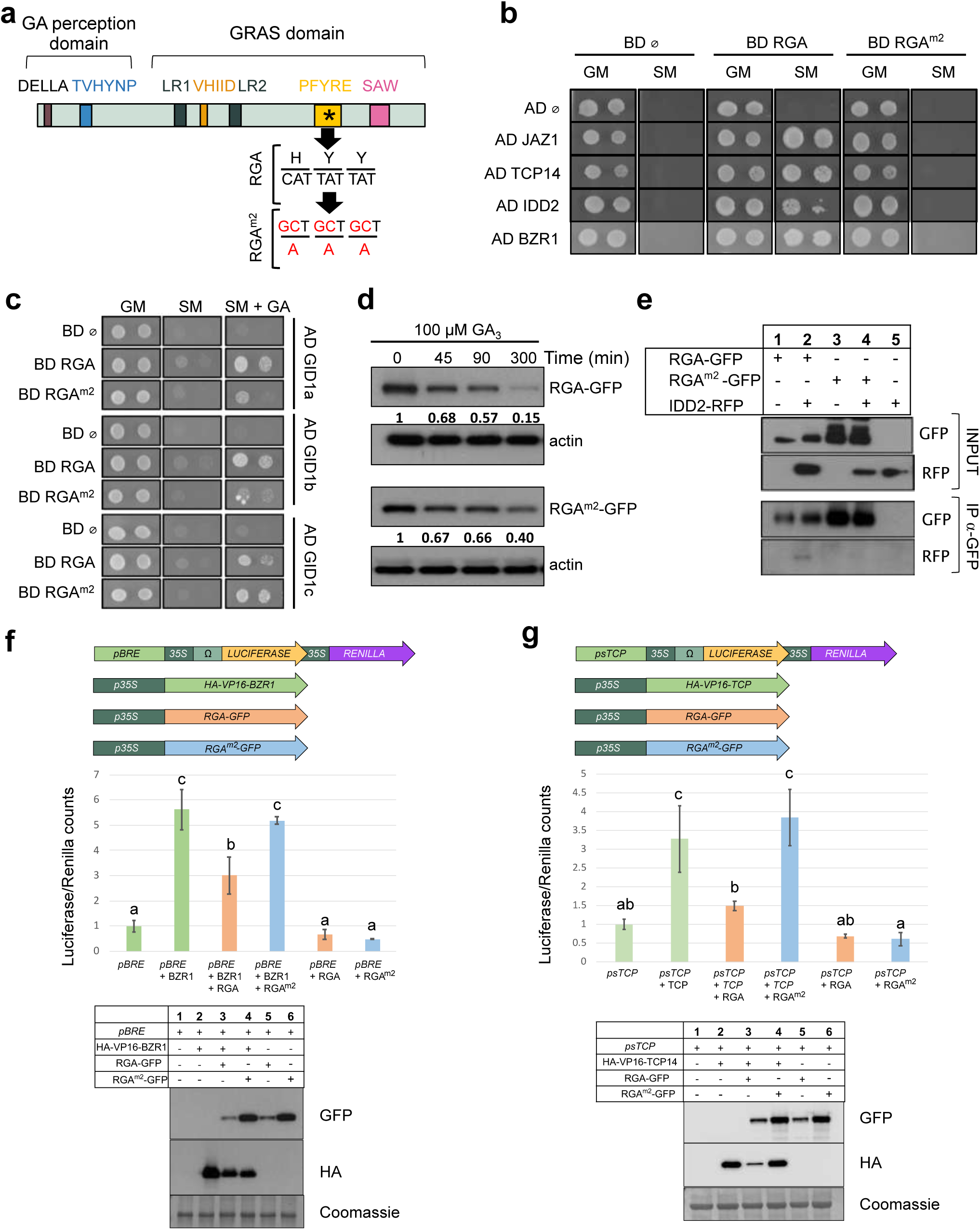
The modified RGA^m2^ protein is an inactive DELLA that remains sensitive to GA. **a**, Schematic representation of the domain structure of a typical DELLA protein. The nucleic acids and amino acids mutated in RGA^m2^ are indicated in red. **b**, Yeast two hybrid (Y2H) assays in which RGA and RGA^m2^ were tested pairwise with four known DELLA-interacting partners: JAZ1, TCP14, IDD2 and BZR1. Empty pGBKT7 and pGADT7 vectors were included as negative controls. Photos show the growth of the yeast on control media (GM) and on selective media (SM). **c**, Pairwise Y2H interaction assays between RGA or RGA^m2^ and the three GA receptors GID1a, GID1b and GID1c. Photos show the growth of the yeast on control media (GM), selective media (SM) and SM media supplemented with 100 μM GA_3_. **d**, Time-course analysis of GA-induced degradation of RGA (upper panel) and RGA^m2^ protein (lower panel). Immunodetection of RGA-GFP and RGA^m2^-GFP protein in *35S::RGA-GFP* and *35S::RGA^m2^-GFP N. benthamiana* agro-infiltrated leaves treated with 100 mM cycloheximide (CHX) and 100 µM GA_3_ for the indicated times. Numbers indicate RGA-GFP and RGA^m2^-GFP levels relative to actin levels, used as loading control. **e**, Coimmunoprecipitation assays between RGA or RGA^m2^ and IDD2. Protein extracts from different combinations of *N. benthamiana* agro-infiltrated leaves with *35S::RGA-GFP*, *35S::RGA^m2^-GFP* and *35S::IDD2-RFP* were immunoprecipitated with anti-GFP antibodies. The co-immunoprecipitated protein (IDD2-RFP) was detected by anti-RFP antibodies. **f-g,** Effect of RGA and RGA^m2^ on BZR1 (**f**) and TCP14 (**g**) transcriptional activities in *N. benthamiana* agro-infiltrated leaves with a combination of BZR1, TCP14, RGA and RGA^m2^ effector constructs and corresponding Luciferase/Renilla reporter constructs, as indicated (top panels). Transcriptional activities are represented as the ratio of Luciferase and Renilla (used as internal control) activities, relative to the value obtained for the reporter construct alone that was set to 1. Different letters denote significant differences (*p* < 0.05) using ANOVA Tukey’s tests for multiple comparisons. Bottom panels: immunodetection of RGA-GFP, RGA^m2^-GFP, HA-VP16-BZR1 and HA-VP16-TCP14 from *N. benthamiana* agro-infiltrated leaves used for transcriptional activity assays.

Based on the above results, we selected RGA^m2^ for further analysis. We next showed that RGA^m2^ is unable to bind with the BZR1 DELLA interacting partner in yeast and a larger screen confirmed that these mutations abolish interactions with practically all known DELLA partners (Fig. 1b, Supplementary Table 1). Moreover, co-immunoprecipitation studies demonstrated that binding of RGA^m2^ to IDD2 is also strongly reduced *in planta* compared to RGA (Fig. 1e). Finally, transient expression assays confirmed that, while RGA represses BZR1 and TCP14 transcriptional activities, RGA^m2^ did not (Fig. 1f-g). Taken together, these data indicate that the expression of RGA^m2^ in plants, hereafter named RGA^mPFYR^, might have a limited impact on GA signalling. Thus, RGA^mPFYR^ constitutes a suitable DELLA variant candidate for engineering a degradation-based biosensor that monitors GA signalling activity and more specifically the combinatorial effect of GA and of its complex perception machinery.

### Engineering a GA signalling sensor (Fig. 2; Extended Data Fig. 2-6)

To create a ratiometric GA signalling sensor, we fused the RGA^mPFYR^ protein to the fast maturing yellow fluorescent protein VENUS^23^ and co-expressed this fusion protein together with a nuclear-localized non-degradable reference protein, TagBFP-NLS, under the *pUBQ10*^24^ or *pRPS5a*^25, 26^ constitutive promoter. We used the 2A self-cleaving peptide to allow for a stoichiometric production of both fluorescent proteins, enabling quantification of GA signalling activity using fluorescence intensity ratio between RGA^mPFYR^-VENUS and TagBFP^27, 28^ (Fig. 2a). We named the sensor lines qRGA^mPFYR^ (<u>q</u>uantitative RGA^mPFYR^) and first analysed their TagBFP fluorescence pattern in vegetative and reproductive tissues (Fig. 2b and Extended Data Fig. 2a-b). TagBFP fluorescence of *pUBQ10::qRGA^mPFYR^* lines was homogeneously distributed in hypocotyl, the vegetative shoot meristem, cotyledons and roots (except in the root tip) but the TagBFP signal was unevenly distributed in the inflorescence SAM, with a stronger signal in organ boundaries (Extended Data Fig. 2a). Conversely, the TagBFP signal was strong in the root tip and the vegetative SAM in *pRPS5a::RGA^mPFYR^* lines (Fig. 2b), and showed a homogenous distribution in the inflorescence SAM (Extended Data Fig. 2b). Hence, the choice of one of these constructs depends on the analysed tissue: we used *pUBQ10::RGA^mPFYR^*in subsequent experiments in seedlings, and *pRPS5a::RGA^mPFYR^* for experiments performed in inflorescence SAM.

**Figure 2.**
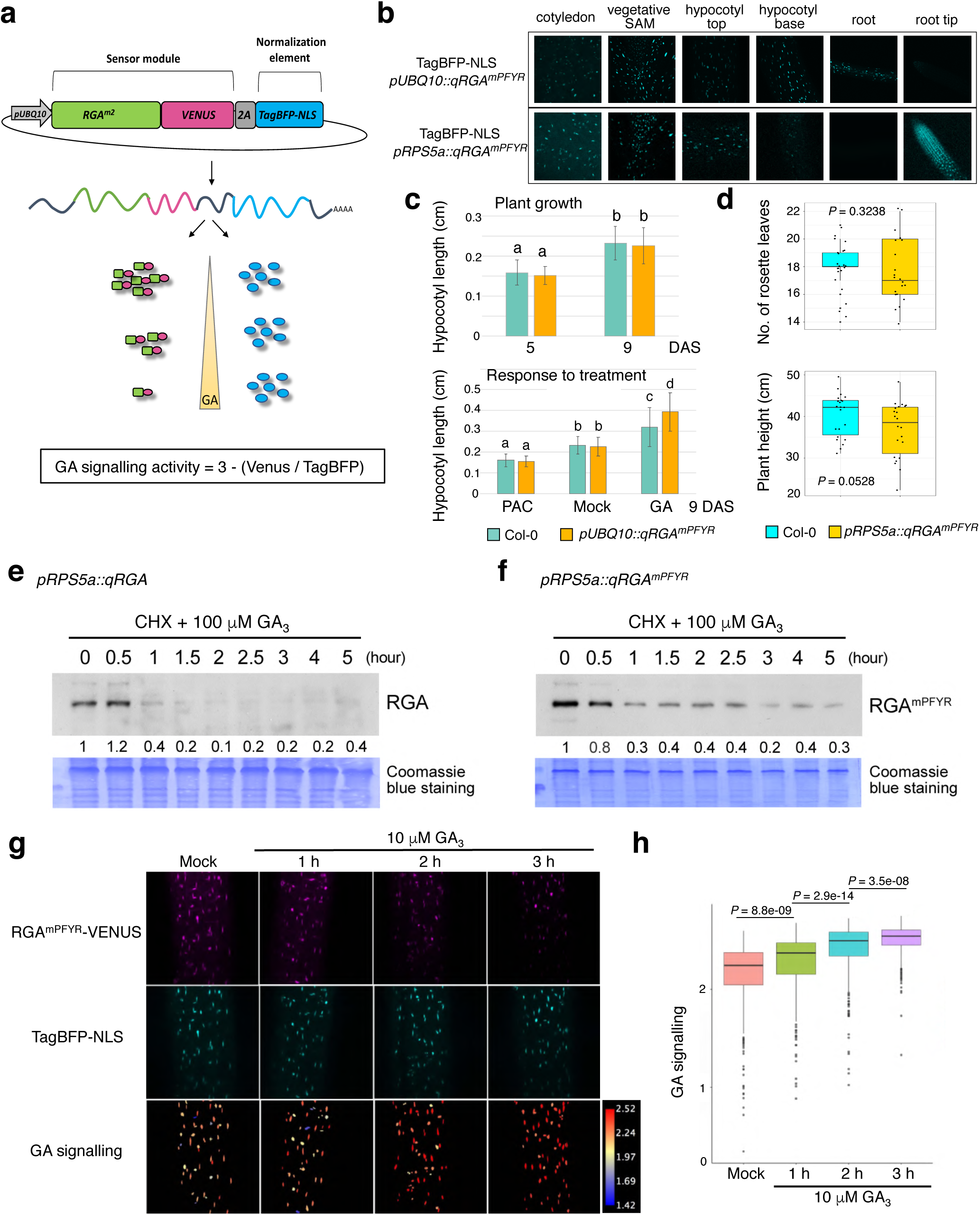
The qRGA^mPFYR^ sensors monitor changes in GA levels. **a**, Schematic representation of the qRGA^mPFYR^ construct composed of two elements: the sensor module (RGA^mPFYR^-VENUS), and the normalization element (TagBFP fused with a nuclear localization signal (NLS)). The two elements are linked by a 2A self-cleaving peptide, and driven by the same promoter allowing stoichiometric expression. GA signalling activity is measured as 3 minus the ratio between VENUS and TagBFP signal intensities. **b**, TagBFP expression pattern of *pUBQ10::qRGA^mPFYR^* and *pRPS5a::qRGA^mPFYR^* sensors monitored in cotyledon, vegetative SAM, hypocotyl and root of 7-day-old seedlings. **c**, Hypocotyl length of wild-type (Col-0) and *pUBQ10::qRGA^mPFYR^* seedlings. Upper panel: Hypocotyl length at 5 and 9 days after sowing (DAS). Lower panel: Hypocotyl length of 9-d-old seedlings grown for 4 days on MS media and then transferred to MS media supplemented with 5 μM PAC or 10 μM GA_3_, and controls (Mock). Different letters denote significant differences (*p* < 0.05) using ANOVA Tukey’s tests for multiple comparisons. **d**, Boxplot representations of the number of rosette leaves (upper panel) and final plant height (lower panel) of wild-type (Col- 0) and *pRPS5a::qRGA^mPFYR^* adult plants. *P* values are from Wilcoxon rank-sum tests. **e**,**f**, Time-course analysis of GA-induced RGA and RGA^mPFYR^ degradation. 7-d-old *pRPS5a::qRGA* (**e**) and *pRPS5a::qRGA^mPFYR^* seedlings (**f**) were treated with 100 µM cycloheximide (CHX) and 100 µM GA_3_. At indicated time points, total proteins were extracted and analysed by immunoblot using RGA antibodies. Numbers indicate the fold increase in RGA and RGA^mPFYR^ protein levels relative to blue-stained protein signal. The experiment was repeated twice with similar results. **g**, Representative confocal images of RGA^mPFYR^-VENUS and TagBFP signals, and corresponding heatmap representation of GA signalling activity in hypocotyls of 5-d-old *pUBQ10::qRGA^mPFYR^* seedlings treated with 10 μM GA_3_ for the time indicated (and Mock control). **h**, Boxplot representation of GA signalling activity (means of at least 8 seedlings) measured in seedling hypocotyls grown in the same conditions as in **g**. *P* values are from Kruskal-Wallis tests. The experiment was repeated twice with similar results.

To test that qRGA^mPFYR^ activity is indeed not interfering with signalling activity and thus with plant growth, we first investigated the effects of exogenous GA and paclobutrazol (PAC; an inhibitor of GA biosynthesis) treatments on the growth of qRGA^mPFYR^ plants. We found that hypocotyl length of qRGA^mPFYR^ seedlings was similar to that of wild-type under mock conditions and upon GA or PAC treatment, although hypocotyls were slightly longer after GA treatment (Fig. 2c). Similarly, shoot development and plant fertility were not significantly affected in qRGA^mPFYR^ plants (Fig. 2d and Extended Data Fig. 2c). Last, we showed that while qd17RGA plants (d17RGA is a mutant version of RGA that is fully insensitive to GA^29^) exhibited a severe dwarf phenotype reminiscent of GA-insensitive mutants, qd17RGA^mPFYR^ plants had similar rosette size and height than wild-type plants (Extended Data Fig. 2d-i). Altogether, these results demonstrate that qRGA^mPFYR^ negligibly interferes with plant growth and GA responses.

Next, we assessed if qRGA^mPFYR^ can detect changes in GA levels. Consistent with the above transient expression assays, GA treatment induced the degradation of RGA^mPFYR^ in *pRPS5a::qRGA^mPFYR^*seedlings, although the protein tends to be slightly more stable than RGA (Fig. 2e,f). Accordingly, while the TagBFP signal was unaffected in hypocotyls of *pUBQ10::qRGA^mPFYR^* seedlings upon GA application, the sensor element RGA^mPFYR^-VENUS fluorescence was substantially reduced after this treatment (Fig. 2g). Similar results were observed in qRGA^mPFYR^ roots for which GA and PAC application respectively reduced and increased RGA^mPFYR^-VENUS signal (Extended Data Fig. 3). In contrast, VENUS fluorescence did not change in any of the treatments in RGA^m3^-VENUS and RGA^m4^-VENUS lines, consistently with the lower GA-dependent degradation rate of these variants (Extended Data Fig. 3).

**Figure 3.**
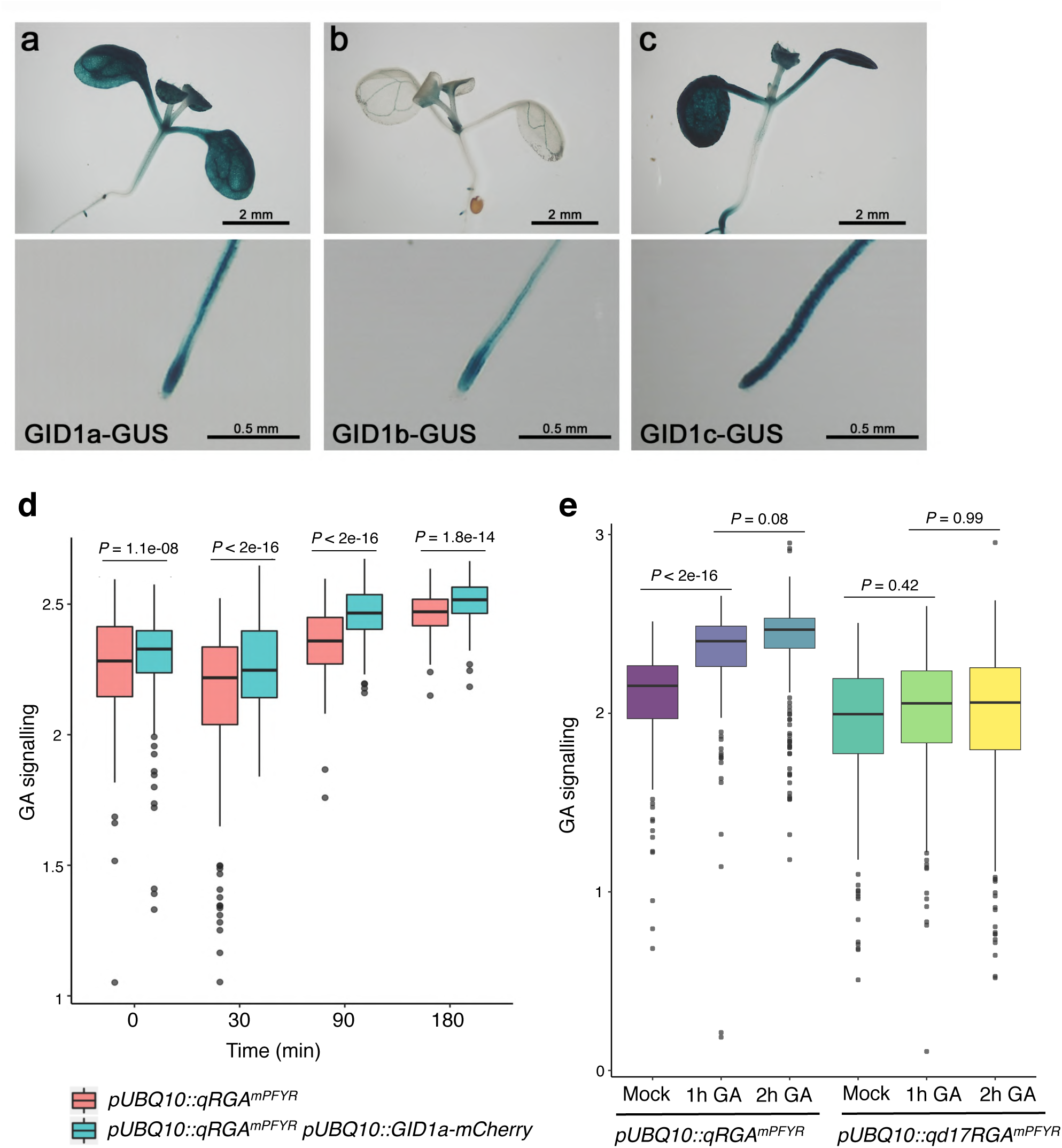
The qRGA^mPFYR^ sensor signal depends on GA receptor activity. **a-c**, Expression pattern of *GID1a*, *GID1b* and *GID1c* in shoots (upper panel) and root tips (lower panel) of 7-d-old *pGID1a::GID1a-GUS* (a), *pGID1b::GID1b-GUS* (b) and *pGID1c:: GID1c-GUS* (c) seedlings. **d**,**e**, Boxplot representations of GA signalling activity in hypocotyls of 5-d-old *pUBQ10::RGA^mPFYR^* and *pUBQ10::RGA^mPFYR^ pUBQ10::GID1a- mCherry* seedlings (**d**) and in *pUBQ10::RGA^mPFYR^* and *pUBQ10::d17RGA^mPFYR^* seedlings (**e**) treated with 100 µM GA_3_ for the time indicated, and untreated controls. *P*-values are from Kruskal-Wallis tests.

In further image analyses, to fully cover a highly variable range of values in VENUS/TagBFP fluorescence ratio, we used 3 – (RGA^mPFYR^-Venus/TagBFP) as a positive proxy for GA signalling activity (hereafter named “GA signalling”; Fig. 2a; see also Supplementary Method 1). This quantitative approach confirmed statistically significant changes in GA signalling in hypocotyls, with an increase and decrease respectively upon GA and PAC treatments compared to untreated seedlings (Fig. 2g,h; Extended Data Fig. 4a-d). Exogenous GA and PAC treatments induced similar responses in the SAM, although the effect of PAC was less pronounced than in hypocotyl (Extended Data Fig. 5). Furthermore, GA signalling activity increased in the SAM with both exogenous GA concentration and treatment duration (Extended Data Fig. 4e-i), showing that qRGA^mPFYR^ is suitable to be used as a GA signalling sensor in the SAM as in all the other tissues tested.

**Figure 4.**
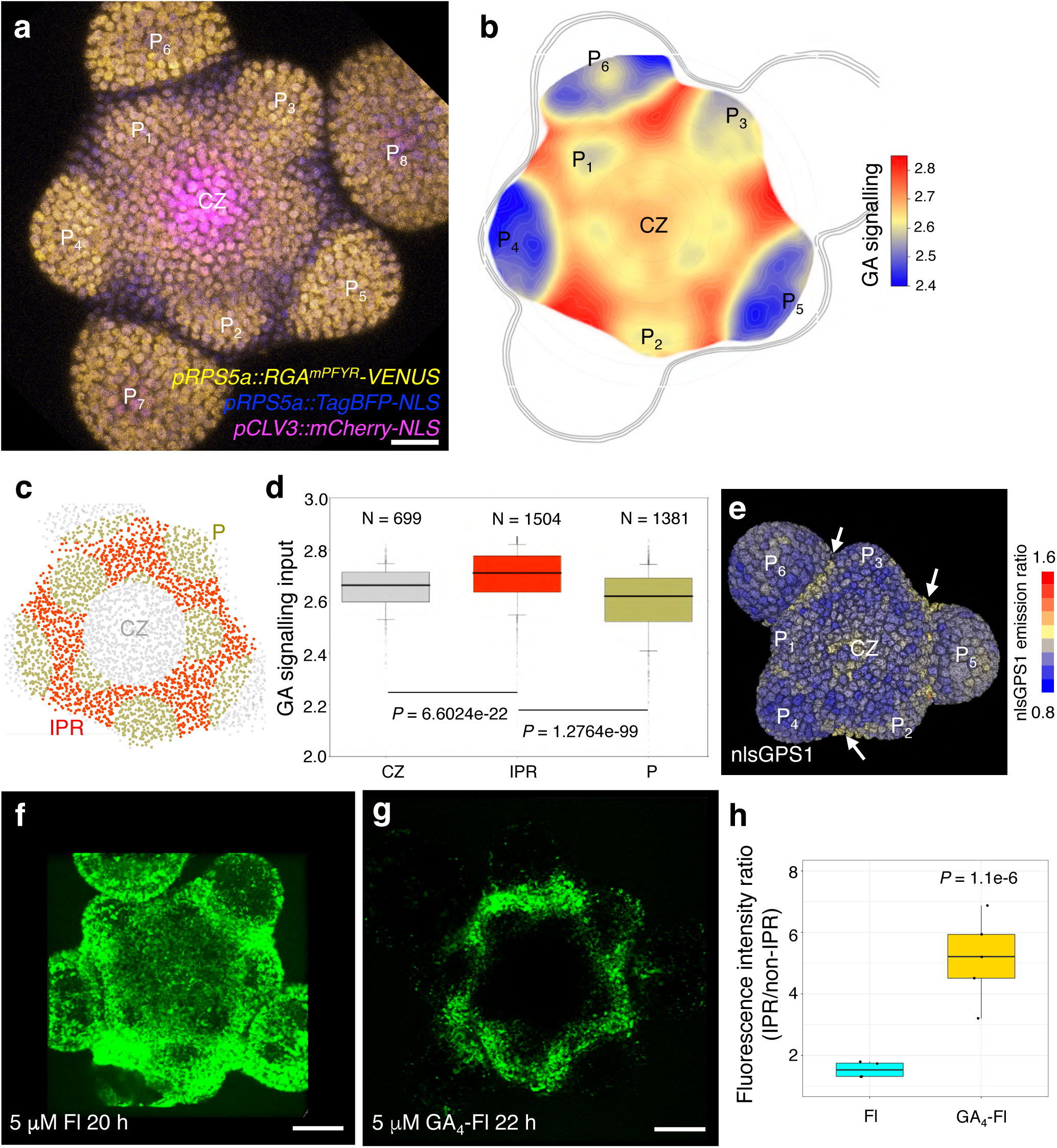
A GA signalling activity map reveals high levels of GA signalling activity in inter-primordia cells in the SAM. **a**, Maximum intensity projection showing the expression of *pRPS5a::qRGA^mPFYR^*and *pCLV3::mCherry-NLS* in the SAM (overlay). CZ, central zone; P, primordium. **b**, Heatmap representation of L1 GA signalling activity averaged from seven SAMs aligned using the CLV3 domain as a reference. **c-d**, Quantification of GA signalling activity in central zone (CZ), inter-primordia region (IPR) and primordia (P) indicated by different colours as shown in (**c**). *P* values are from one-way ANOVA. **e,** 3D visualization of nlsGPS1 emission ratio in the SAM. Primordia stages are estimated according to morphology. Arrows highlight higher nlsGPS emission ratio in the boundaries and IPR. **f-g,** Fluorescence distribution in wild-type (L*er*) SAMs treated with fluorescein (Fl, **f**) and GA_4_-Fl (**g**). **h**, Comparison of the ratio of average fluorescence intensity in the IPR to that in the non-IPR (excluding primordia) between Fl and GA_4_-Fl treatment in the SAM. *P* value is from one-way ANOVA with Turkey’s test for multiple comparisons. The experiment was repeated twice with similar results. Scale bars = 20 μm.

**Figure 5.**
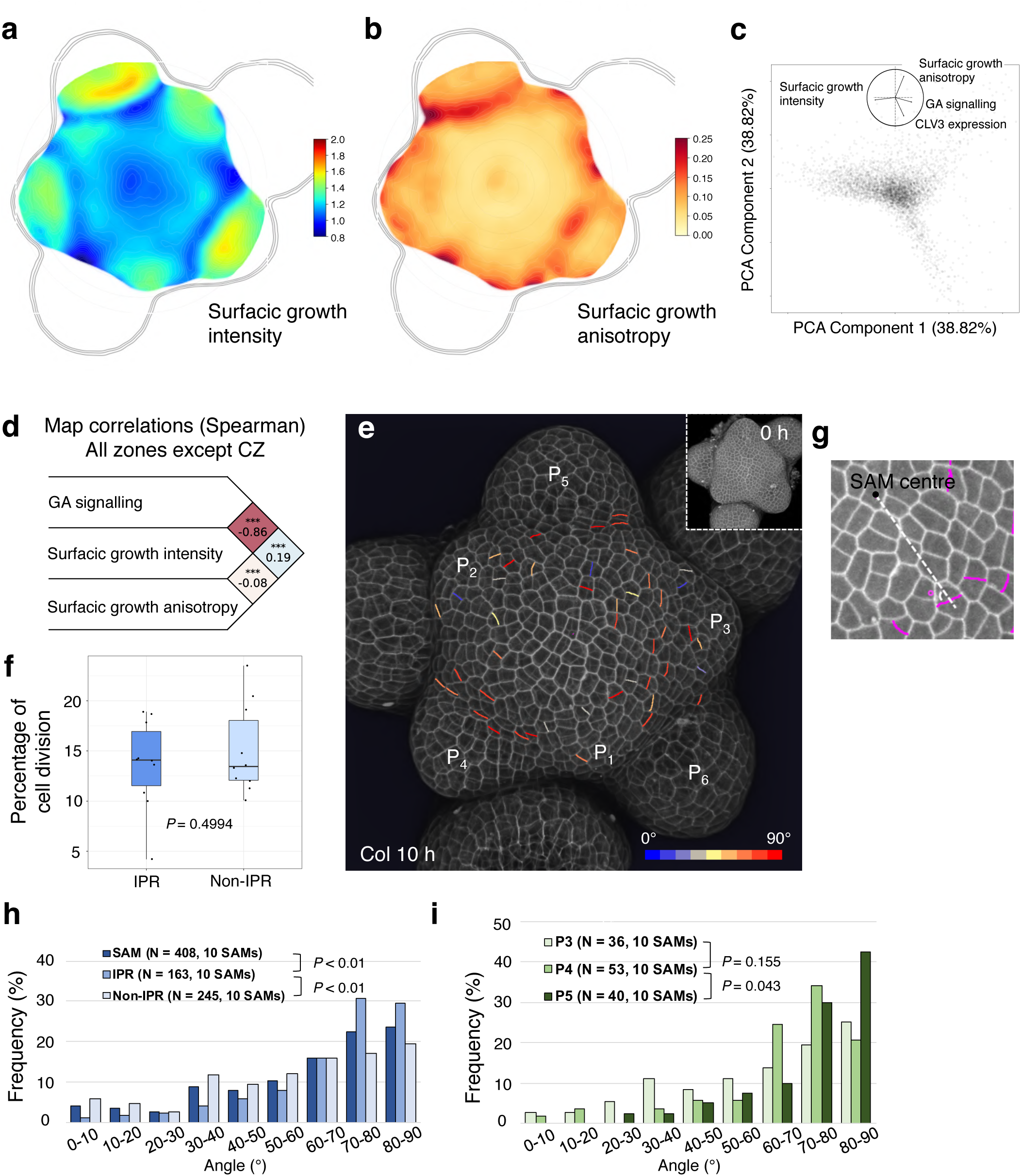
High GA signalling activity correlates positively with growth anisotropy and transverse cell division orientation. a-b, Averaged surfacic growth (**a**) and growth anisotropy (**b**) heat maps (used as proxies for cell expansion intensity and direction, respectively) in the SAM. **c**, PCA analysis including the following variables: GA signalling, surfacic growth intensity, surfacic growth anisotropy and CLV3 expression. **d**, Spearman correlation study between GA signalling, surfacic growth intensity and surfacic growth anisotropy in tissue scale but excluding CZ. **e**, 3D visualization of Col-0 SAM L1 cells using confocal microscopy. New cell walls formed in the SAM (but not primordia) in 10 h are coloured according to their angle values. The insert shows the corresponding 3D image at 0 h. **f**, Boxplot representation of cell division frequency in the IPR and non-IPR of Col-0 SAM. *P* value is from Welch two samples t-test. **g**, A schematic diagram showing how the angle of a new cell wall (magenta) relative to the radial direction from the centre of the SAM (white dotted line) was measured. Only acute angle values (i.e: 0- 90°) are considered. **h**, Frequency histograms of division plane orientation of cell from the entire SAM (dark blue), the IPR (medium blue) and non-IPR (light blue), respectively. **i**, Frequency histograms of division plane orientation of cells located in front and sides of P_3_ (light green), P_4_ (mediate green) and P_5_ (dark green), respectively. *P* values are from Kolmogorov-Smirnov tests. The experiment was repeated twice with similar results.

Finally, we asked whether qRGA^mPFYR^ is able to report for changes in endogenous GA levels using growing hypocotyls. We previously showed that nitrate promotes growth by increasing GA synthesis and in turn DELLA degradation^30^. Accordingly, we observed that hypocotyl length of *pUBQ10::qRGA^mPFYR^*seedlings grown on adequate nitrate supply (10 mM NO_3_^-^) was significantly longer compared to those grown on nitrate-deficient conditions (Extended Data Fig. 6a). Consistent with the growth response, GA signalling was higher in hypocotyls of seedlings grown with 10 mM NO_3_^-^ compared with those grown in absence of nitrate (Extended Data Fig. 6b,c). Thus qRGA^mPFYR^ also allows monitoring changes in GA signalling resulting from endogenous changes in GA concentration.

### qRGA^mPFYR^ fluorescence depends on GA receptor activity (Fig. 3 and Extended Data Fig. 7)

As GA signalling activity depends on both GA concentration and GA perception, we analysed the expression of the three *GID1* receptors in vegetative and reproductive tissues. In seedlings, GID1-GUS reporter lines showed that *GID1a* and *c* are highly expressed in cotyledons (Fig. 3a-c). Moreover, all three receptors are expressed in leaves, lateral root primordia, root tip (excluding root cap for *GID1b*) and vasculature (Fig. 3a-c). In inflorescence SAM, we only detected a GUS signal for GID1b and 1c (Extended Data Fig. 7a-c). *In situ* hybridization confirmed these expression patterns and further demonstrated that *GID1c* is homogenously expressed at a low level in the SAM while *GID1b* shows higher expression at the SAM periphery (Extended Data Fig. 7d-l). A *pGID1b::2xmTQ2-GID1b* translational fusion further revealed a graded *GID1b* expression ranging from low or no expression in the SAM centre to high expression in organ boundaries (Extended Data Fig. 7m). Thus, GID1 receptors are unevenly distributed across and within tissues. In a subsequent experiment, we also observed that overexpressing *GID1* (*pUBQ10::GID1a-mCherry*) enhanced qRGA^mPFYR^ sensitivity to external GA application in hypocotyls (Fig. 3d). By contrast, the fluorescence measured from qd17RGA^mPFYR^ in hypocotyls was insensitive to GA (Fig. 3e). Taken together, these results confirm that the qRGA^mPFYR^ biosensor reports for the combinatorial action of GA and GA receptors and suggest that differential expression of GID1 receptors can modulate the emission ratio of the sensor.

### A GA signalling map in the shoot apical meristem (Fig. 4; Extended Data Fig. 8-10)

Distribution of GA signalling within the SAM has remained elusive so far. Thus, we used plants expressing qRGA^mPFYR^ together with the *pCLV3::mCherry-NLS* stem cell reporter to compute a high-resolution quantitative map of GA signalling activity, focusing on the L1 layer (epidermis; Fig. 4a-b, see Methods and Supplementary Method 1) given the key role of L1 in controlling growth at the SAM^31^. Here, expression of *pCLV3::mCherry-NLS* provided a fixed geometric reference for analysing the spatio-temporal distribution of GA signalling activity^32^. Although GAs were proposed to be required for lateral organ development^4^, we observed that GA signalling was lower in flower primordia (P) from the P_3_ stage onward (Fig. 4a-b), while young P_1_ and P_2_ primordia had intermediate activity similar to the one found in the central zone (Fig. 4a-b). Higher GA signalling activity was found in the boundaries of organ primordia, starting from P_1_/P_2_ (on the lateral sides of the boundary) and culminating from P_4_, and in all peripheral zone cells located between primordia (Fig. 4a-b; Extended Data Fig. 8a-b). This higher GA signalling activity was not only monitored in the epidermis, but also in the L2 and upper L3 layers (Extended Data Fig. 8b). This reveals a GA signalling pattern mostly mirroring primordia distribution. This inter-primordia-region (IPR) distribution results from the progressive establishment of a high GA signalling activity between developing primordia and the central zone, while in parallel GA signalling activity decreases in primordia (Fig. 4c-d).

The distribution of GID1b and GID1c receptors (Extended Data Fig. 7b-c, g-m) suggests that differential expression of GA receptors contributes to shape the GA signalling activity pattern in the SAM. We wondered if differential accumulation of GA could also be involved. To investigate this possibility, we used the nlsGPS1 GA FRET sensor^17^. An increased emission ratio was detected in nlsGPS1 SAMs treated with 10 µM GA_4+7_ for 100 min (Extended Data Fig. 9), indicating that nlsGPS1 responds to changes in GA concentration in the SAM as it does in roots^17^. The spatial distribution of the nlsGPS1 emission ratio indicates that GA levels are relatively low in the SAM external layers, but it shows that they are elevated in the centre of the SAM and in the boundaries (Fig. 4e, Extended Data Fig. 9a,c). We also treated SAMs with fluorescent GAs (GA_3_-, GA_4_-, GA_7_-Fl) or with Fl alone as a negative control. The Fl signal was distributed in the whole SAM, including the central zone and primordia, although with lower intensity (Fig. 4f, Extended Data Fig. 10d). In contrast, all the three GA-Fls specifically accumulated in primordia boundaries and, to different degrees, in part of the rest of the IPR, with GA_7_-Fl accumulating in the largest domain in the IPR (Fig. 4g, Extended Data Fig. 10). Fluorescence intensity quantification demonstrated higher IPR to non-IPR intensity ratio in the GA-Fl treated SAMs, compared to Fl-treated SAMs (Fig. 4h, Extended Data Fig. 10c). Taken together, these results suggest that GAs are present at higher levels in the IPR cells closest to organ boundaries. This indicates that the SAM GA signalling activity pattern results from both differential expression of the GA receptors and from the differential accumulation of GA in IPR cells closest to organ boundaries. Our analysis thus identifies an unexpected spatio-temporal GA signalling pattern, with a lower activity in the centre of the SAM and in primordia, while activity is elevated in the IPR of the peripheral zone.

### Correlation analyses suggest a role for GA signalling in cell division plane orientation in the shoot apical meristem (**Fig. 5**)

To understand the role of differential GA signalling activity at the SAM, we analysed the correlation between GA signalling activity, cell expansion and cell division using time-lapse live imaging of qRGA^mPFYR^ *pCLV3::mCherry-NLS* SAMs. Given the role of GA in growth regulation, a positive correlation was expected with cell expansion parameters. We thus first compared maps of GA signalling activity to those of cell surfacic growth rate (as a proxy for cell expansion intensity for a given cell and daughter cells if it divides) and growth anisotropy, which measures the directionality of cell expansion (here also for a given cell and daughter cells if it divides; Fig. 5a-b, see Methods and Supplementary Method 1). Our cell surfacic growth intensity map of the SAM was consistent with previous observations^33, 34^, with a minimal growth rate in boundaries and a maximal rate in developing flowers (Fig. 5a). A principal component analysis (PCA) showed an anti-correlation between GA signalling activity and cell surfacic growth intensity (Fig. 5c). It further showed that the main axis of variability, encompassing GA signalling input and growth intensity, was orthogonal to the direction defined by high expression of *CLV3*, which argues in favour of excluding cells from the centre of the SAM in the rest of the analysis. Spearman correlation analyses confirmed the PCA results (Fig. 5d), suggesting that higher GA signalling in the IPR does not lead to higher cell expansion. However, the correlation analysis demonstrated a mild positive association between GA signalling activity and growth anisotropy (Fig. 5c,d), suggesting that higher GA signalling in the IPR acts on cell growth orientation and possibly cell division plane positioning.

Thus, we next studied the correlation between GA signalling and cell division activity, by identifying newly formed cell walls during a time course analysis (Fig. 5e). This methodology allows us to measure both the frequency and orientation of cell divisions. Strikingly, we found that cell division frequency was similar in the IPR and the rest of the SAM (non-IPR, Fig. 5f), showing that differences in GA signalling between IPR and non-IPR cells do not have a major effect on cell division. This, together with the positive correlation between GA signalling and growth anisotropy, led us to ask whether GA signalling activity could act on cell division plane orientation. We measured the orientation of new cell walls as the acute angle relative to the radial axis connecting the centre of the meristem to the centre of new cell walls (Fig. 5g), and observed that cells had a clear tendency to divide at angles closer to 90° relative to the radial axis, with the highest frequency observed at 70-80° (23.28%) and 80-90° (22.62%) (Fig. 5h) i.e. corresponding to cell divisions oriented in the circumferential/transverse direction. To explore the contribution of GA signalling to this cell division behaviour, we analysed separately the cell division parameters in the IPR and non-IPR (Fig. 5h). We observed that the angle of cell division was different in the IPR compared to non-IPR or the entire SAM, with IPR cells showing a higher rate of transverse cell divisions, i.e. 70-80° and 80-90° (33.86% and 30.71% respectively) (Fig. 5h). Thus, our observations reveal a link between high GA signalling and cell division plane orientation that parallels the correlation between GA signalling activity and growth anisotropy (Fig. 5c,d). To further establish spatial conservation of this link, we measured the orientation of division planes in IPR cells above primordia starting at stage P_3_, given that the highest GA signalling activity is detected in this region from stage P_4_ (Fig. 4). The division angles of the IPR above P_3_ and P_4_ did not show statistically significant differences, although an increase in the frequency of transverse cell divisions was observed in the IPR above P_4_ (Fig. 5i). Differences in cell division plane orientation were however statistically significant in IPR cells above P_5,_ in which the frequency of transverse cell divisions was drastically increased (Fig. 5i). Taken together, these results suggest that GA signalling could control orientation of cell division in the SAM coherently with previous reports^35, 36^, with high GA signalling likely inducing a transverse orientation of cell divisions in the IPR.

### GA signalling activity positively regulates transverse cell divisions in the shoot apical meristem (Fig. 6; Extended Data Fig. 11-13)

Cells in the IPR are expected not to be incorporated into primordia but rather in the internodes^2, 37, 38^. A transverse orientation of cell divisions in the IPR could generate the typical organization in parallel longitudinal cell files of the internode epidermis. Our observations above indicate that GA signalling is likely to act in this process by regulating cell division orientation.

Loss-of-function of multiple *DELLA* genes leads to constitutive GA response, and thus *della* mutants could be used to test this hypothesis^39^. We first analysed the expression patterns of the five *DELLA* genes in the SAM. Transcriptional fusion GUS lines^40^ showed that *GAI, RGA, RGL1*, and, to a much lesser extent, *RGL2* are expressed in the SAM (Extended Data Fig. 11a-d). *In situ* hybridization further showed that *GAI* mRNA specifically accumulates in primordia and developing flowers (Extended Data Fig. 11e). *RGL1* and *RGL3* mRNA were detected throughout the SAM dome and in older flowers, while *RGL2* mRNA was more abundant in the boundary regions (Extended Data Fig. 11f-h). Confocal imaging of *pRGL3::RGL3-GFP* SAM confirmed the expression observed with *in situ* hybridization and showed that the RGL3 protein accumulates in the central part of the SAM (Extended Data Fig. 11i). Using a *pRGA::GFP-RGA* line, we also found that the RGA protein accumulates in the SAM, but its abundance is reduced in boundaries starting from P_4_ (Extended Data Fig. 11j). Notably, the expression patterns of *RGL3* and *RGA* are compatible with a higher GA signalling activity in the IPR, as detected with qRGA^mPFYR^ (Fig. 4). In addition, these data indicate that all *DELLAs* are expressed in the SAM and that collectively, their expressions cover the entire SAM.

We next analysed the cell division parameters in wild-type (L*er*, control) and quintuple (global) *gai-t6 rga-t2 rgl1-1 rgl2-1 rgl3-4 della* mutants (Fig. 6a-b). Interestingly, we observed a statistically significant change in frequency distribution of cell division angles in the SAM of global *della* mutant compared to wild-type (Fig. 6c). This change in the global *della* mutant resulted from an increase in frequency of 80-90° angles (34.71 % vs 24.55 %) and to a lesser extent of 70-80° angles (23.78 % vs 20.18 %), i.e. corresponding to transverse cell divisions (Fig. 6c). The frequency of non-transverse divisions (0-60°) was also lower in global *della* mutant (Fig. 6c). The increased occurrence of transverse cell divisions was easily visible in the SAM of global *della* mutant (Fig. 6b). The frequency of transverse cell divisions in the IPR was also higher in global *della* mutant compared to wild-type (Fig. 6d). Outside of the IPR region, distribution of the angle of cell division was more homogeneous in wild-type, while in the global *della* mutant it was skewed toward tangential divisions, as in the IPR (Fig. 6e). We thus show that constitutive activation of GA signalling in the SAM induces transverse cell division both in the IPR and the rest of the SAM.

**Figure 6.**
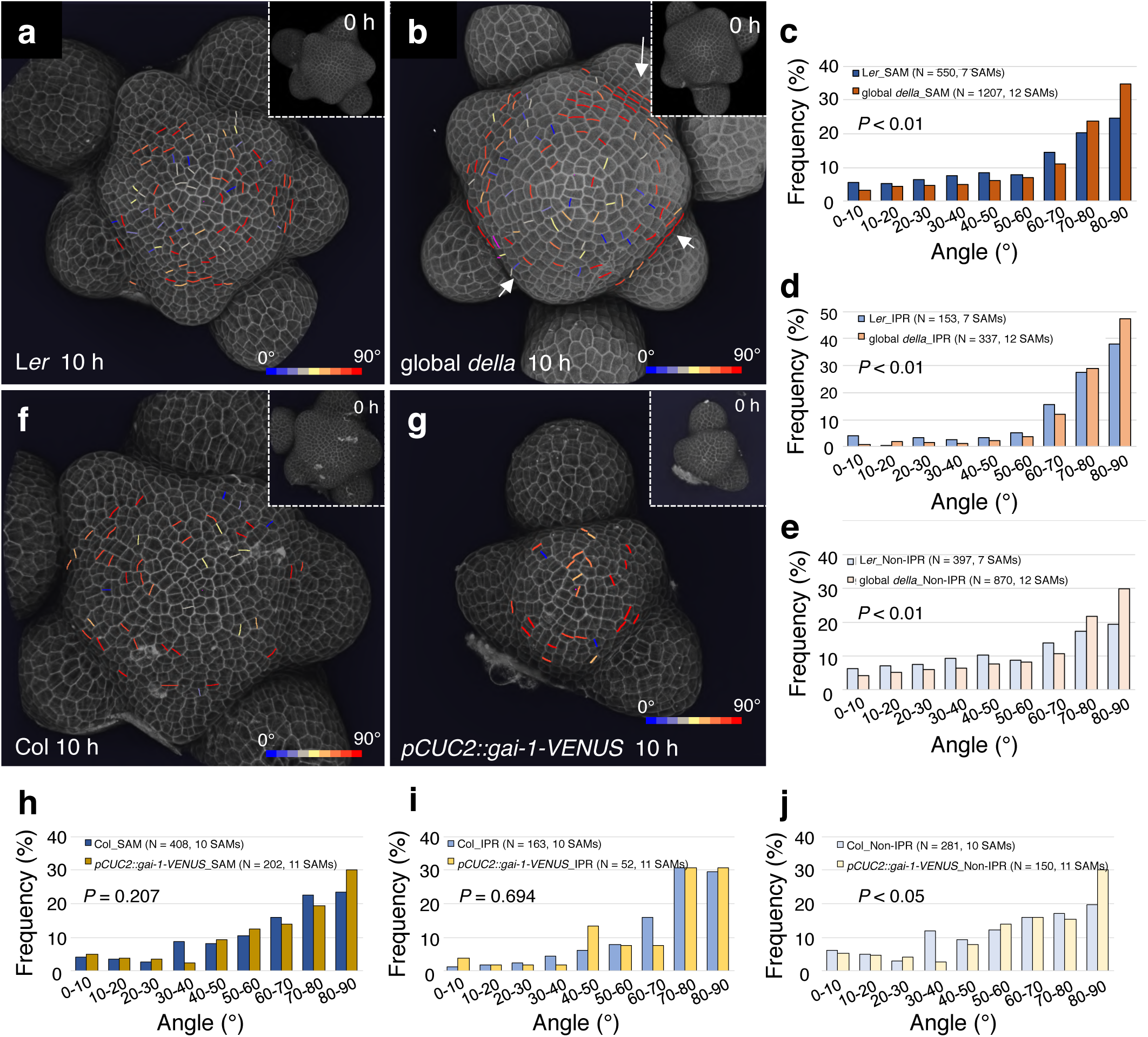
Cell division orientation distribution in the SAM are modified in global *della* mutant and *pCUC2::gai-1-VENUS* transgenic plant. a-b, 3D visualisation of the L1 layer of PI-stained L*er* (**a**) and global *della* mutant (**b**) SAM using confocal microscopy. New cell walls formed in the SAM (but not primordia) in 10 h are shown and coloured according to their angle values. Inserts show the SAM at 0 h. Arrows in (**b**) highlight examples of aligned cell files in global *della* mutant. **c**-**e**, Comparison of frequency distribution of division plane orientation of cells in the entire SAM (**d**), IPR (**e**) and non-IPR (**f**) between L*er* and global *della*. *P* values are from Kolmogorov-Smirnov tests. The experiment was repeated three times with similar results. **f-g**, 3D visualisation of confocal image stacks of PI-stained SAMs of Col-0 (**i**) and *pCUC2::gai-1-VENUS* (**j**) transgenic plants. New cell walls formed in the SAM (but not primordia) in 10 h are shown as in (**a**,**b**). **h-j**, Comparison of frequency distribution of division plane orientation of cells located in the entire SAM (h), IPR (i) and non-IPR (j) between Col-0 and *pCUC2::gai-1-VENUS* plants. *P* values are from Kolmogorov-Smirnov tests. The experiment was repeated three times with similar results.

We then tested the effect of inhibiting GA signalling specifically in the IPR. To do so, we used the *CUP-SHAPED COTYLEDON 2* (*CUC2*) promoter to drive expression of the dominant negative gai-1 protein fused to VENUS (in a *pCUC2::gai-1-VENUS* line). *CUC2* promoter drives expression in a large part of the IPR (including the boundary cells) in the SAM starting from P_4_ (Extended Data Fig. 12k). The distribution of cell division angles in the entire SAM or in the IPR of *pCUC2::gai-1-VENUS* plants did not show statistically significant differences with respect to the wild-type, although unexpectedly, we found in these plants a higher frequency of 80-90° divisions in non-IPR cells (Fig. 6f-j).

The orientation of cell division has been proposed to be influenced by the geometry of the SAM and notably by tensile stresses prescribed by the curvature of the tissue^41^. We thus asked whether the SAM shape of the global *della* mutant and *pCUC2::gai-1-VENUS* plants was changed. As previously shown^12^, the size of the global *della* mutant SAM was bigger than wild-type (Extended Data Fig. 12a,b,d). However, the SAM curvature was identical in the two genotypes (Extended Data Fig. 12b,e,f,g,h,j). We observed a comparable increase in size in the quadruple *gai-t6 rga-t2 rgl1-1 rgl2-1 della* mutant, again without modification of the curvature compared to wild-type (Extended Data Fig. 12c,d,f,i,j). Frequency of cell division orientation was also affected in the quadruple *della* mutant, but to a lesser extent than in the global *della* mutant (Extended Data Fig. 13). This dosage effect, along with the absence of effects on curvature, suggests that the remaining RGL3 activity in quadruple *della* mutants limits the changes in cell division orientation caused by the loss of DELLA activity, and that changes in the occurrence of transverse cell division depend on changes in GA signalling activity rather than in SAM geometry. By contrast, the size of *pCUC2::gai-1-VENUS* SAMs was reduced, whereas it had a significantly higher curvature (Extended Data Fig. 12l-q). This change in the *pCUC2::gai-1-VENUS* SAM curvature could generate a mechanical stress distribution whereby high circumferential stress starts at a short distance from the SAM centre^42^. This could counteract in part the effect of GA signalling changes by increasing the probability of cell division with circumferential/transverse orientation, thus explaining our observations.

Taken together, our data support a positive role for higher GA signalling in transverse orientation of the cell division plane in the IPR. They further suggest that the curvature of the meristem can also influence the orientation of the cell division plane in the IPR.

### High GA signalling initiates internode specification in the shoot apical meristem (Fig. 7; Extended Data Fig. 14)

Transverse orientation of division planes in the IPR as a result of higher GA signalling activity opens the possibility that GAs pre-organize radial cell files in the epidermis within the SAM to specify the cellular organization later found in the internode epidermis. Indeed, such cell files are often visible in the SAM images of the global *della* mutant (Fig. 6b). Therefore, to further understand the developmental function of the GA signalling spatial pattern in the SAM, we used time-lapse imaging to analyse the cell spatial organization in the IPR in wild-type (L*er* and Col-0), global *della* mutant and *pCUC2::gai-1-VENUS* transgenic plants.

We marked L*er* cells above and on the side of P_4_ according to their developmental fates (analysed 34 hrs after the first observation, i.e. more than two plastochrones) using three different colours: yellow for those incorporated into the primordia in the vicinity of P_4_, green for those located in the IPR and magenta for those contributing to both (Fig. 7a-c). At t0 (0 h), 1-2 layers of IPR cells were visible in front of P_4_ (Fig. 7a). As expected, when these cells divide, they mostly do it with a transverse division plane (Fig. 7a-c). Similar results were obtained with Col-0 SAMs (focusing on P_3_ that had a comparable folding at the boundary than P_4_ in L*er*), although in this genotype the formation of the crease at the flower boundary hides the IPR cells more rapidly (Fig. 7g-i). Thus, their division pattern and localization in the IPR support the idea that these cells correspond to internode precursors.

**Figure 7.**
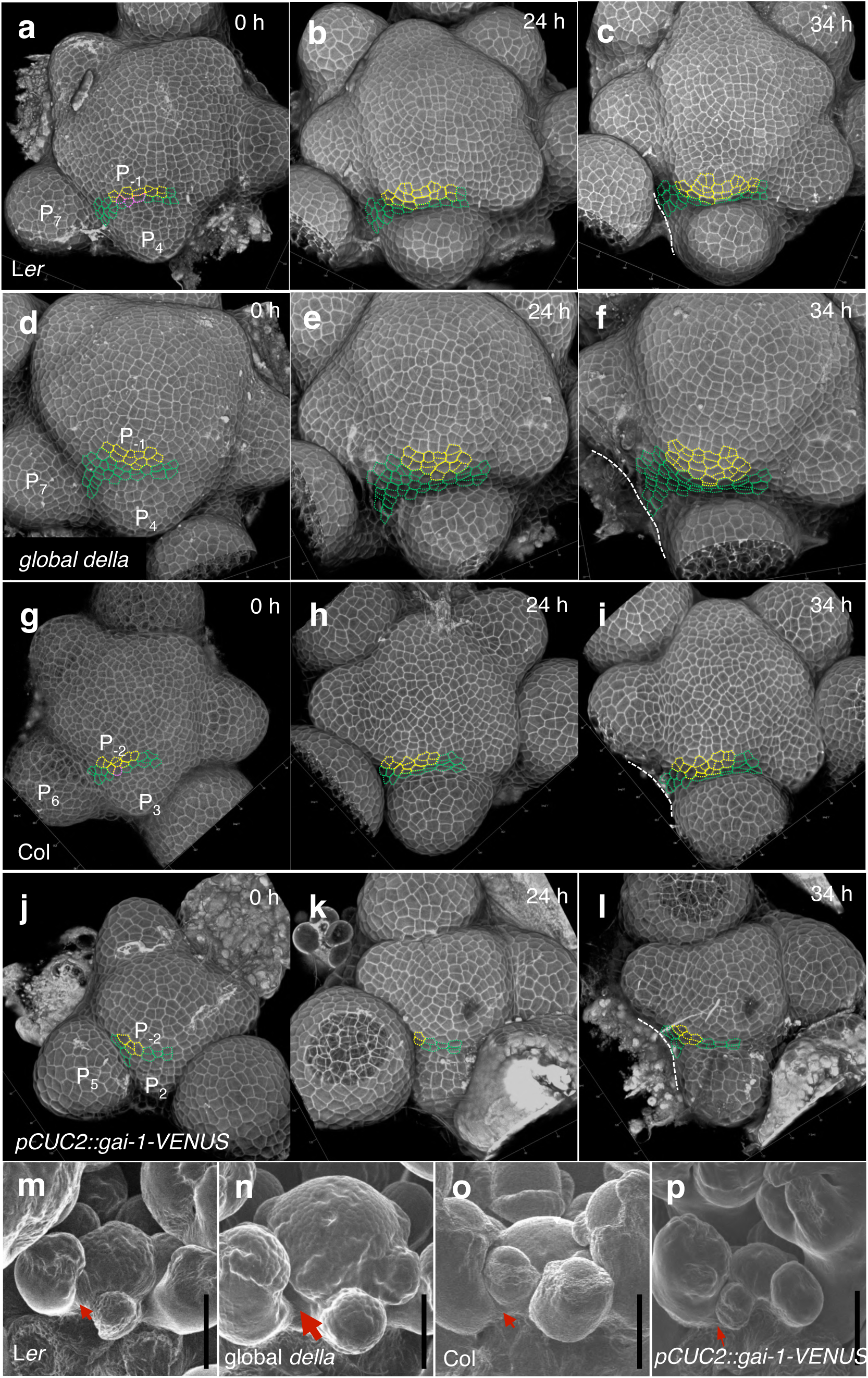
Emergence of the internode cellular organisation in the inter-primordia region in wild-type and in plants with modified GA signalling activity. **a-l**, Time-lapse (0 h, 24 h, 34 h) visualization of the L1 of the SAM from confocal mciroscopy of L*er* (**a-c**), global *della* (**d-f**), Col-0 (**g-i**) and *pCUC2::gai-1-VENUS* transgenic (**j-l**). The imaging time is indicated at the right up corner of each panel. Cells outlined with yellow dotted lines are those incorporated into a primordium at 34 h, and cells in green are found in the IPR at 34 h. Those in magenta produce both primordium and IPR cells. White dotted lines mark the edge of the elder primordium (that have been removed at 34h except in (**c**)). **m-p,** Scanning electron microscopy images of the shoot apex of L*er* (**m**), global *della* (**n**), Col-0 (**o**), and *pCUC2::gai-1-VENUS* (**p**) allowing to analyse the early establishment of internode just below the SAM. The different size of arrows indicates the range of early internode length in the different genetic background.

Compared to L*er*, 1-2 extra layers of IPR cells were observed in front of P_4_ at t0 (0 h) in the SAM of global *della* mutants. These cells divided several times in 34 hrs (Fig. 7d-f, compared to 7a-c), and in consequence, the mostly transverse divisions of IPR cells led to a higher population of cells organized in radial cell files (Fig. 7d-f, compared to 7a-c). This indicates that the higher GA signalling activity in global *della* mutant SAMs promotes internode specification. We conducted a similar analysis in *pCUC2::gai-1-VENUS* plants. Since expression of this transgene causes changes in SAM geometry (Extended Data Fig. 12m,n,p,q), we analysed IPR cells above the first primordium showing a comparable folding at the boundary as Col-0 P_3_. In these plants, opposite to global *della* mutants, much less cell divisions occurred in the IPR and there was no clear sign of an organization in radial cell files (Fig. 7j-l), thus showing that inhibition of GA signalling in the IPR perturbs the specification of the cellular organisation of internodes in the SAM. In lines with these results, we were able to detect the appearance of internodes in the global *della* mutant just below the SAM using electron microscopy, while flowers remained compacted in L*er* (Fig. 7m-n). By contrast, organs were much more compacted in the SAM of *pCUC2::gai-1-VENUS* than in Col-0 (Fig. 7o-p), consistent with the taller and shorter inflorescence stem of the global *della* mutant^39, 43^ and *pCUC2::gai-1-VENUS* plants, respectively (Extended Data Fig. 14a-b). Our results thus support the hypothesis that higher GA signalling activity in the IPR specifies the cellular organization of internodes in the SAM, through a regulation of the orientation of cell division planes (Extended Data Fig. 15).

## Discussion

Here, we developed a new ratiometric GA signalling biosensor, qRGA^mPFYR^, that provides information on GA function at the cellular level by allowing quantitative mapping of GA signalling activity that results from the combinatorial action of GA and GA receptors concentrations, with minimal interference with the endogenous signalling pathway. To this end, we have engineered a modified DELLA protein, RGA^mPFYR^, that has lost its capacity to bind DELLA interacting partners but that remains sensitive to GA-induced proteolysis. qRGA^mPFYR^ responds to both exogenous and endogenous changes in GA levels and its dynamic sensing properties allow the assessment of spatio-temporal changes in GA signalling activity during developmental processes. qRGA^mPFYR^ is also a highly flexible tool as it can be adapted to a variety of tissues simply by changing, if needed, the promoter used for its expression and is very likely transferable to other species given the conserved nature of the GA signalling pathway and of the PFYRE motif in angiosperms.

Internode development is a key trait for plant architecture and crop improvement. qRGA^mPFYR^ revealed a higher GA signalling activity specifically in cells of the IPR, that are the precursors of internodes. By combining quantitative image analysis and genetics, we show that the GA signalling pattern imposes circumferential/transverse cell division planes in the SAM epidermis, shaping the cell division organization required for internode development. Few developmental regulators of the orientation of the cell division plane during development have been identified^44, 45^. Our work provides a striking example where GA signalling activity regulates this cellular parameter. DELLA can interact with the prefoldin complex^36^ and GA signalling could thus regulate the orientation of the cell division plane through a direct effect on cortical microtubule orientation^35, 36, 46, 47^. The fact that we show that unexpectedly not cell elongation nor cell division but only growth anisotropy correlates in the SAM with higher GA signalling activity is coherent with a direct effect of GAs on cell division orientation in the IPR. However, we cannot eliminate the possibility that this effect could also be indirect, e.g. mediated by GA-induced softening of the cell wall^48^. Changes in cell wall properties induces mechanical stress^49, 50^ that could also affect cell division plane orientation by acting on cortical microtubule orientation^34, 41, 51^. A combined effect of GA-induced mechanical stress and direct regulation by GA of microtubule orientation could then participate to create the specific pattern of cell division orientation in the IPR to specify the internode and further work is needed to test this idea. Likewise, previous works have highlighted the importance of the DELLA interacting proteins TCP14 and 15 in the control of internode patterning^52, 53^ and these factors could convey GA action, together with *BREVIPEDICELLUS* (*BP*) and *PENNYWISE* (*PNY*), which regulate internode development and have been shown to affect GA signalling^2, 54^. Early cytological studies showed that both the inner tissues and the peripheral zone of the SAM are required for internode development in *Arabidopsis*^2, 37^. The fact that GAs positively regulate cell division in the inner tissues^12^ supports a dual function of GAs in regulating meristem size and internode at the SAM. Patterns of oriented cell division are also highly regulated in the inner SAM tissues, and this regulation is essential to stem growth^44^. It will be interesting to explore whether GAs also play a role in orienting cell division planes in the inner tissues of the SAM and thus synchronize internode specification and development within the SAM.

## Methods

### Growth conditions and Plant material

Plants were grown on soil or *in vitro* on 1x Murashige-Skoog (MS) medium (Duchefa) supplemented with 1% sucrose and 1% agar (Sigma) under standard conditions (16h photoperiod at 22°C), except for hypocotyl and root growth experiments, for which the seedlings were grown on vertical plates under continuous light and 22°C. For experiments with nitrate, plants were grown on MS modified medium without nitrogen (bioWORLD plant media) supplemented with an adequate nitrate concentration (0 or 10 mM KNO_3_), 0.5 mM NH_4_-succinate, 1% sucrose and 1% type-A agar (Sigma) under long-day photoperiod.

We used the following *Arabidopsis* mutants and transgenic lines in the L*er* background: *gai- t6 rga-t2 rgl1-1 rgl2-1* quadruple *della*^39^, *gai-t6 rga-t2 rgl1-1 rgl2-1 rgl3-4* global *della*^55^ or Col-0: *pGID1a::GID1a-GUS*, *pGID1b::GID1b-GUS*, *pGID1c::GID1c-GUS*, *pGAI::GUS, pRGA::GUS*, *pRGL1::GUS*, *rgl2-5* (a promoter trap GUS line)^56^, *pRGA::GFP-RGA*^8^, *pRGL3::RGL3-GFP*, *pCLV3::mCherry-NLS*^57^, *nlsGPS1*^17^.

All other transgenic lines were generated in the Col-0 background. To generate these lines and to perform transient assays in *Nicotiana benthamiana* leaves, plasmids were constructed in the following way. RGA cDNA (AGI code AT2G01570) and RGA^m1^, RGA^m2,^ RGA^m3^ and RGA^m4^ mutant variants were obtained by PCR using specific primers numbered 1 to 8 in Supplementary Table 2, and inserted into pDONR221 (Thermo Fisher Scientific) by Gateway cloning and recombined with pB7FWG2^58^ to generate *p35S::RGA-GFP* and *p35S::RGA^m1/m2/m3/m4^-GFP*. To generate *pRPS5a::qRGA, pUBQ10:qRGA*, *pRPS5a::qRGA^mPFYR^*, *pUBQ10:qRGA^mPFYR^*, *pRPS5a::RGA^m1/m3/m4^-VENUS-2A-BFP*, *pUBQ10::RGA^m1/m3/m4^-VENUS-2A-BFP*, and *pCUC2::gai-1-VENUS*, the promoter region of *pRPS5a* (1.7 kb fragment), *pUBQ10* (2.5 kb fragment) or *pCUC2* (3.2 kb fragment) inserted into pDONR P4-P1R (Thermo Fisher Scientific), *RGA-VENUS*, *RGA^mPFYR^ -VENUS*, *RGA^m1/m3/m4^ -VENUS* or *gai-1* cDNA inserted into pDONR221, and *2A-TagBFP* or *VENUS* (for *pCUC2::gai-1-VENUS)* inserted into pDONR P2R-P3 (Thermo Fisher Scientific), were recombined into pB7m34GW^58^.

d17RGA (RGA deleted of the 17 amino acids DELLAVLGYKVRSSEMA composing the DELLA domain^29^) and d17RGA^mPFYR^ mutant variant obtained by PCR using primers 1, 2, 5, and 6 (Supplementary Table 2) were inserted into pDONR221 and recombined into pB7m34GW with *p35S* (inserted into pDONR P4-P1R) and *VENUS* (inserted into pDONR P2R-P3) to generate *p35S::d17RGA-VENUS* and *p35S::d17RGA^mPFYR^-VENUS*. *d17RGA-VENUS* and *d17RGA^mPFYR^-VENUS* were then amplified by PCR using primers 1 and 12, inserted into pDONR221 and recombined as previously with *pRPS5a* or *pUBQ10* and *2A- TagBFP* to generate *pRPS5a::qd17RGA*, *pUBQ10:qd17RGA*, *pRPS5a::qd17RGA^mPFYR^*, *pUBQ10:qd17RGA^mPFYR^*.

To obtain *pUBQ10::GID1a-mCherry*, *GID1a* cDNA inserted into pDONR221 was recombined with pDONR P4-P1R-*pUBQ10* and pDONR P2R-P3-*mCherry* into pB7m34GW. *p35S:IDD2-RFP* was obtained by recombining *IDD2* cDNA inserted in pDONR221 into pB7RWG2^58^. To get *pGID1b::2xmTQ2-GID1b*, a 3.9-kb fragment upstream of the coding region of *GID1b* and a 4.7-kb fragment including *GID1b* cDNA (1.3 kb) and terminator (3.4 kb) were first amplified using primers in Supplementary Table 2, then inserted into pDONR P4-P1R (Thermo Fisher Scientific) and pDONR P2R-P3 (Thermo Fisher Scientific), respectively, and finally recombined into pGreen 0125^59^ destination vector with pDONR221 2xmTQ2^60^ by Gateway cloning. To make *pCUC2::LSSmOrange*, the promoter sequence of *CUC2* (3229 bp upstream the ATG), followed by the coding sequence for the large stokes shift mOrange (LSSmOrange)^61^ with a N7 nuclear localization signal, and the NOS transcription terminator, was assembled into the pGreen Kanamycin destination vector using 3 fragment Gateway recombination system (Invitrogen). For transactivation assays, *pTCP- LUC* and *pBRE-LUC* reporter constructs (containing synthetic promoters) and *p35S::3xHA-VP16-TCP14* and *p35S::3xHA-VP16-BZR1* encoding effector proteins were used as previously described^62, 63^. Plant binary vectors were inserted into *Agrobacterium tumefaciens* GV3101 strain and respectively introduced into *Nicotiana benthamiana* leaves by agro- infiltration and *Arabidopsis* Col-0 by floral dip. *pUBQ10::qRGA^mPFYR^ pUBQ10::GID1a- mCherry* and *pCLV3::mCherry-NLS qRGA^mPFYR^* were isolated from F3 and F1 progeny of the appropriate crosses, respectively.

### Pharmacological treatments

Chemical treatments with GA (GA_3_, Sigma or Duchefa) and paclobutrazol (PAC, Duchefa) were performed at the concentrations and time indicated in figures. For Y2H assays, 100 µM GA_3_ were added in selective media to promote interaction between RGA and GID1. For RGA and RGA^m1/m2/m3/m4^ degradation kinetics, *N. benthamiana* agro-infiltrated leaf discs were incubated with 100 µM GA_3_ and 100 mM cycloheximide (Sigma) over 300 min. For hypocotyl length measurements and fluorescence analyses of qRGA^mPFYR^ hypocotyls and roots, seedlings were grown for 5 days on MS agar medium and then transferred for 4 days on MS agar plates supplemented with 5 µM PAC or 10 µM GA_3_. For GA_3_ treatment on qRGA^mPFYR^, GA_4+7_ treatment on nlsGPS1, and GA-Fl treatment on L*er*, dissected shoot apices were inserted into Apex Culture Medium (ACM, 1/2x MS medium (Duchefa), 1% sucrose, 1% agarose, 2 mM MES (Sigma), 1x vitamin solution (myo-Inositol 100 mg/L, nicotinic acid 1 mg/L, pyridoxine hydrochloride 1 mg/L, thiamine hydrochloride 10 mg/L, glycine 2 mg/L), 200 nM N6-Benzyladenine) with indicated concentrations of GA/GA-Fl, and also immersed under 200 µl of GA/GA-Fl solution of indicated concentrations for a indicated period of time. For PAC treatment on qRGA^mPFYR^, 50 µM PAC were sprayed on the whole inflorescence every two days for a period of 5 days before observation.

### Yeast two-hybrid assays

For Y2H assays, both the full-length and the C-terminal part of RGA and RGA^m1/m2/m3/m4^ (named M5 version, amino acids 199 to 587; the N-terminal part is subject to self-activation in yeast^64^) inserted into pDONR221 were recombined into pGBKT7 (Clontech) to obtain BD- RGA, BD-RGA^m1/m2/m3/m4^, BD-M5RGA and BD-M5RGA^m1/m2/m3/m4^. On the other hand, JAZ1, TCP14, IDD2, BZR1, GID1a, GID1b and GID1c cDNAs inserted into pDONR221 were fused to the activation domain GAL4 (AD) after recombination into pGADT7 (Clontech).

Direct interaction assays were carried out following the Clontech procedures. Briefly, BD- M5DELLA and AD-JAZ1, AD-TCP14, AD-IDD2, AD-BZR1 and, on the other hand, BD- DELLA (full length) and AD-GID1 constructs were co-transformed in the yeast strain AH109 and interactions tests were surveyed on selective medium lacking tryptophan, leucine and histidine. In some cases, the medium was supplemented with 3-amino-1, 2, 4 triazole (3AT, Sigma) or with GA to promote interaction between DELLA and GID1.

To confirm that the RGA^m2^ mutation impaired interaction with all RGA protein partners, we used the C-terminal part of RGA and RGA^m2^ (amino acids 199 to 587), to probe the Arabidopsis transcription factors REGIA + REGULATORS (RR) arrayed library, following the protocol described in Castrillo et al.^65^. TFs in the RR library were fused to GAL4 activation domain of the pDEST22 vector and independently transformed into the yeast strain YM4271 in 96-well plates. The RGA^m2^ protein fused to the GAL4 BD in the pDEST32 vector was transformed into the pJ694 yeast strain and used as bait. Replicates of the library were grown overnight on SD-Trp solid media and inoculated together with 100 μL of an overnight RGA^m2^ culture grown on SD-Leu on microtiter plates containing 100 μL of YPDA per well. Plates were incubated for 2 days at 30°C for mating, and diploid colonies were selected in new 96-well plates containing 200 μL of SD-Leu/Trp. 5 μL of the diploid cell cultures were lastly tested for protein interaction by placing them on solid SD medium lacking both Leu and Trp (positive growth control), and on SD medium lacking Leu, Trp and His, in the presence of 1 mM 3-aminotriazol (3-AT) (Sigma-Aldrich). Results were expressed in the form of a heat map for the strength of interaction according to the colony growth after five days of incubation at 30°C.

### Co-immunoprecipitation assays

CoIP assays were performed on *N. benthamiana* agro-infiltrated leaves with *p35S::IDD2- RFP*, *p35S::RGA-GFP* or *p35S::RGA^m2^-GFP*. Three days after infiltration, total proteins were extracted with the native extraction buffer [Tris-HCl (pH 7.5) 50 mM, glycerol 10%, Nonidet P-40 0.1% supplemented with Complete Protease Inhibitors 1X (Roche)], and then incubated for 2h at 4°C with 50 µl of anti-GFP antibody conjugated with paramagnetic beads (Miltenyi Biotec). After incubation, samples were loaded onto a magnetic column system (µ columns ; Miltenyi Biotec) to recover the immunoprotein complexes according to manufacturer’s protocol. The immunoprecipitated (RGA-GFP and RGA^m2^-GFP) and co-immunoprecipitated (IDD2-RFP) proteins were detected by western-blot with anti-GFP (JL8; Clontech) and anti-RFP (6G6; Chromotek), respectively.

### Immunodetection analyses

Plant materials were ground in 2x SDS-PAGE buffer followed by heating at 95°C for 5 min. After centrifugation at 13000 *g* for 5 min, total proteins were separated on 8.5% SDS-PAGE gel and transferred to an immobilon-P (PVDF) membrane (Millipore). Membranes are then saturated with blocking buffer (TBS 1x, Tween-20 0.1%; milk 5%) and incubated with a 2000-fold dilution of anti-GFP (JL8; Clontech), anti-RGA (Agrisera), anti-HA (Sigma) or anti-actin (Agrisera) antibodies, and a 5000-fold dilution of peroxidase-conjugated goat anti- rabbit or mouse IgG (Invitrogen). Signals were detected using the Luminata Forte Western HRP Substrate (Millipore).

### Transactivation assays

Transactivation assays were performed using the Dual-Glo Luciferase Assay System (Promega) according to manufacturer’s instructions. Briefly, *N. benthamiana* leaves were agro-infiltrated with *pTCP-LUC* and *pBRE-LUC* reporter constructs, *p35S::3xHA-VP16- TCP14* or *p35S::3xHA-VP16-BZR1* encoding effector proteins, and *p35S::RGA-GFP* or *p35S::RGA^m2^-GFP*. Three days after infiltration, total proteins were extracted in lysis buffer (Promega), then Firefly and control Renilla LUC activities were quantified with FLUOstar Omega luminometer (BMG Labtech) using OMEGA2 software version 5.50 R4. For loading control, protein levels were analysed by immunodetection.

### GUS staining

For GUS expression detection in inflorescence apex, 28-d-old plants grown under LD conditions were used. After fixation in 90% cold acetone at room temperature for 20 min, shoot apexes were transferred into GUS staining solution containing 1 mM potassium ferrocyanide, 1 mM potassium ferricyanide and 1 mg/ml 5-bromo-4-chloro-3-indolyl-*ß*-D- glucuronide (X-Gluc), vacuum-infiltrated on ice for 15 min, and incubated overnight at 37 °C in dark before washing with ethanol series and microscope detection. For the experiment with seedlings, 7-d-old plantlets grown in vitro under LD conditions were vacuum-infiltrated for 15 min in GUS staining solution containing 2 mM potassium ferrocyanide, 2 mM potassium ferricyanide and 0.25 mg/ml X-Gluc, and incubated at 37°C for 24 h. Then GUS solution was replaced with ethanol 70% and seedlings were observed using optical microscope.

### *In situ* hybridizations

RNA in situ hybridizations were performed according to Vernoux et al.^66^. with some minor modifications on fixation. Briefly, shoot apices with ∼1cm stem were collected and immediately fixed in FAA solution (3.7% Formaldehyde, 5% Acetic Acid, 50% Ethanol) precooled at 4 °C. After vacuum treatments for 2x 15 min, the fixative was changed and the samples were incubated overnight. Antisense probes of the cDNA and 3’-UTR of *GID1a*, *GID1b*, *GID1c*, *GAI*, *RGL1*, *RGL2* and *RGL3* were synthesized as described in Rozier et al.^67^. using primers indicated in Supplementary Table 2. Immunodetection of the digoxigenin-labelled probes was performed using an anti-digoxigenin antibody, and sections were stained with 5-Bromo-4-chloro-3-indoryl Phosphate (BCIP)/Nitroblue Tetrazolium (NBT) solutions.

### Microscopy

For confocal microscopy observations of hypocotyl, complete seedlings were put on slides and images were obtained with a Zeiss LSM 780 confocal microscope. SAM live imaging was performed as described^32^. Briefly, dissected SAMs were let to recover overnight after dissection. Confocal images were taken with a Zeiss LSM 710 or 700 confocal laser scanning microscope equipped with a water-dipping lens (W Plan-Apochromat 40x/1.0 DIC). The confocal settings were as previously described^32^ and detailed below. To observe GFP or GFP with propidium iodide (PI), a laser of 488 nm was used to excite, and the emission for GFP was 500-520 nm (in some cases it was 500-540 nm), for PI was 610-650 nm. For imaging of qRGA^mPFYR^ or qRGA^mPFYR^ with mCherry and PI, lasers of 514 nm, 405 nm, 561 nm and 488 nm were used for the excitation of VENUS, TagBFP, mCherry and PI, respectively, and the corresponding emission wavelength was 520-560 nm, 430-460 nm, 580-615 nm and 620-660 nm. mTURQUOISE2 (mTQ2) was excited by 445 nm laser and the emission range was 470-510 nm. For FRET detection of nlsGPS1, an acquisition mode of spectral imaging (λ-scan) with emission wavelength from 461 nm to 597 nm was used with excitation at 458 nm.

For optical microscopy, photographs of plants were taken with a LEICA MZ12 stereoscopic microscope equipped with a ZEISS AxioCam ICc5 camera head or Canon camera.

For scanning electronic microscopy of fresh shoot apex, a Hirox SH-3000 table-top microscope equipped with Coolstage (-20 °C to -30 °C) was used.

### Image processing

Confocal stacks were processed in Fiji (fiji.sc) to get max projection or orthogonal views. Older primordia were also removed in Fiji. For 3D visualization of confocal stacks, we used the Zeiss ZEN2 software (Fig. 7) and a rendering using the VTK library^68^ (Fig. 5e, Fig. 6). In both cases, parameters were adjusted accordingly to show mainly the L1 cells.

### Quantification and statistical analysis GA sensor quantification

For qRGA^mPFYR^ quantification and visualization, images of hypocotyls were analyzed using a Python script that performs nuclei detection and signal quantification in 3D. The details of the algorithms used in the analysis are provided in Supplementary Method 1 (Section 2 – Nuclei detection and signal quantification).

For nlsGPS1 quantification and visualization, a series of Fiji macros were used. Briefly, a “seg-auto.ijm” macro was run for segmentation of nuclei based on the sum of signals from all the 14 channels of the spectral imaging. After removing little objects corresponding to signal noise using a “object-screening.ijm” macro, the ratio of DxAm (channel 8) to DxDm (channel 3) of each segmented nucleus was calculated by running a “3D-ratio.ijm”, and from here, the 3D construction of the SAM, with every nucleus showing its ratio value, was also achieved and printed. Statistical analysis was done in R.

### SAM image sequence analysis

To analyze time-lapse confocal acquisitions of SAMs and obtain quantitative measures of GA signalling as well as cellular growth parameters, we developed a computational pipeline. It consists of 3D watershed segmentation for cell identification from the PI staining, nuclei detection and qRGA quantification, temporal registration using the PI image channel and surfacic growth estimation based on expert cell lineages. Individual SAM sequences were then aligned in order to perform population-scale statistics. Extensive details on the algorithms used in this pipeline are provided in Supplementary Method 1.

### Hypocotyl length measurement

To measure hypocotyl length, seedlings were grown in vertical MS medium (Duchefa) with 0.8% (w/v) phytoagar and 1% sucrose for 5 days and transferred to MS, MS supplemented with 10 μM GA_3_ or 5 μM Paclobutrazol for 4 days in continuous light at 22°C. Plates were scanned and hypocotyl length was measured from the images using Fiji.

### Cell division orientation quantification

New cell walls were identified by comparing images obtained at 0 h and 10 h, and masked with a manually drawn line in Fiji. Then, a macro was used to skeletonize and then to measure the angles of the drawn cell walls with an expert-defined centre of the SAM. Further angle frequency distribution was done in Excel using Pivot table. Statistical analyses of Kolmogorov-Smirnov tests were performed using an online tool: http://www.physics.csbsju.edu/stats/KS-test.n.plot_form.html.

### Curvature quantification

The MorphoGraphX software was used to quantify cellular curvature of L1 cells of SAMs of different genetic backgrounds. Statistical analyses were done in R.

### Meristem size measurement

To measure meristem size, a method described before was used^69^. Briefly, the SAM radius was determined by drawing a circle that covers I_1_ and I_2_ and that the center of which is roughly overlapping with the geometrical center of the SAM surface using Fiji software. Statistical analysis was done in R.

### Synthesis and characterization of GA-Fluorescein (GA-Fl)

GA-Fluorescein (GA-Fl) were synthesized using a previously described protocol^15^. Extensive details on synthesis and characterization are provided in Supplementary Method 2.

### Data and software availability

All experimental data and quantified data that support the findings of this study are available from the corresponding authors upon request. Quantitative image and geometry analysis algorithms are provided in Python libraries timagetk, cellcomplex, tissue_nukem_3d and sam_atlas (https://gitlab.inria.fr/mosaic/) made publicly available under the CECILL-C license. The script used to process hypocotyl images is publicly available in the qrga_nuclei_quantification project (https://gitlab.inria.fr/gcerutti/). The pipelines used to analyze SAM image sequences and to produce map visualizations are provided in a separate project (https://gitlab.inria.fr/mosaic/publications/sam_spaghetti) as Python scripts.

---

**Extended Data Figure 1.**
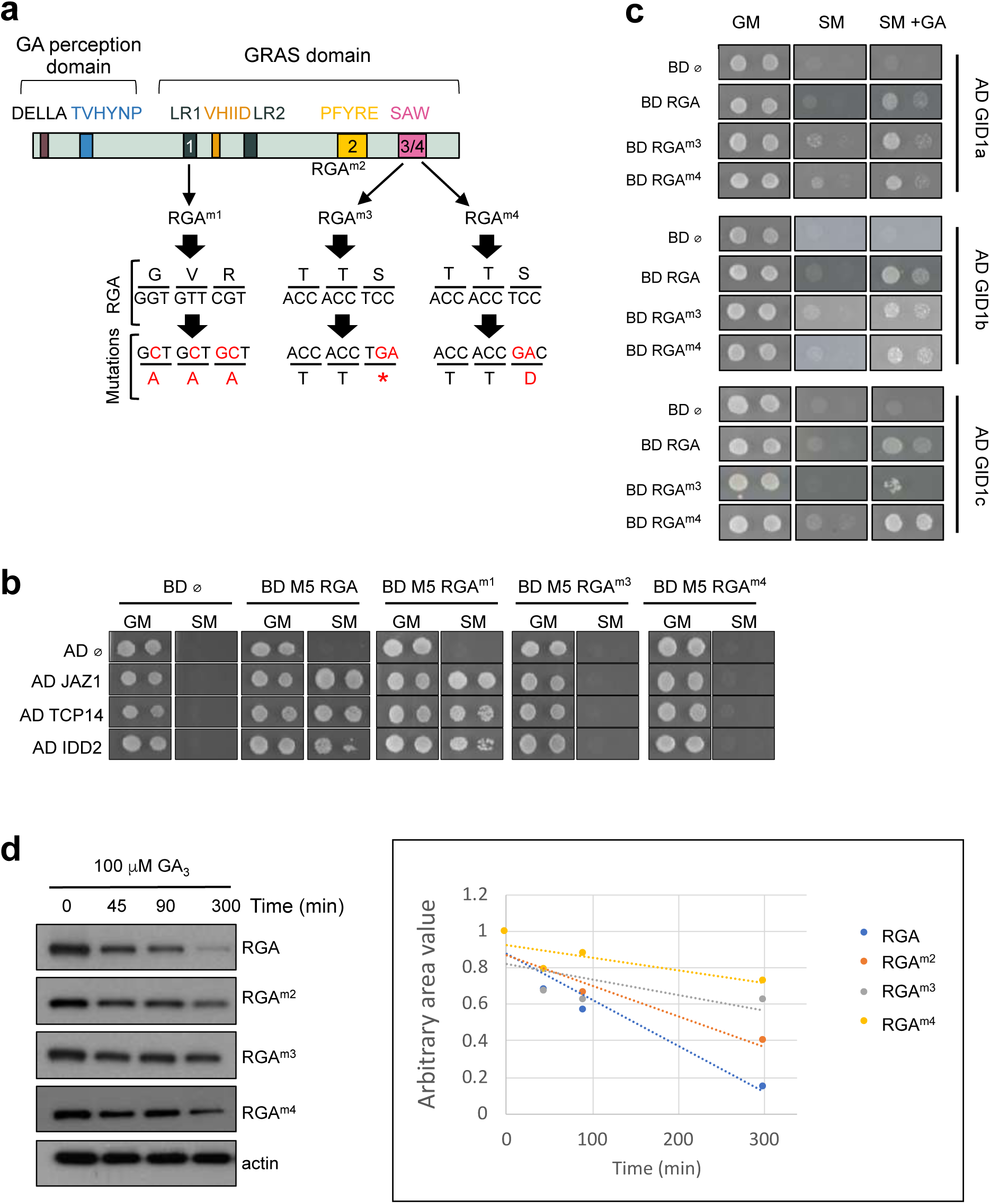
Characterization of mutant DELLA proteins. **a**, Position of the mutated nucleic acids and amino acids of the four candidate modified DELLA proteins used in the study: RGA^m1^, RGA^m2^, RGA^m3^, RGA^m4^. *: stop codon. **b**, Pairwise Y2H interaction assays between RGA, RGA^m1^, RGA^m3^, RGA^m4^ and three DELLA-interacting proteins, JAZ1, TCP14 and IDD2. Empty pGBKT7 and pGADT7 vectors were included as negative controls. Photos show the growth of the yeast on control media (GM) and on selective media (SM). **c**, Pairwise Y2H interaction assays between RGA, RGA^m3^, RGA^m4^ and GID1a, GID1b and GID1c. Photos show the growth of the yeast on control media (GM), selective media (SM) and SM media supplemented with 100 μM GA_3_. **d**, Time-course analysis of GA-induced degradation of RGA, RGA^m2^, RGA^m3^ and RGA^m4^. Left panel: Immunodetection of RGA-GFP, RGA^m2^-GFP, RGA^m3^-GFP and RGA^m4^-GFP proteins respectively in *35S::RGA-GFP*, *35S::RGA^m2^-GFP*, *35S::RGA^m3^-GFP* and *35S::RGA^m4^-GFP N. benthamiana* agro-infiltrated leaves treated with 100 mM cycloheximide (CHX) and 100 µM GA_3_ for the indicated times. Actin is used as sample loading control. Right panel: Quantification of the immunoblot signals depicted as a graph. Values indicate RGA-GFP, RGA^m2^-GFP, RGA^m3^-GFP and RGA^m4^-GFP signals relative to actin signals.

**Extended Data Figure 2.**
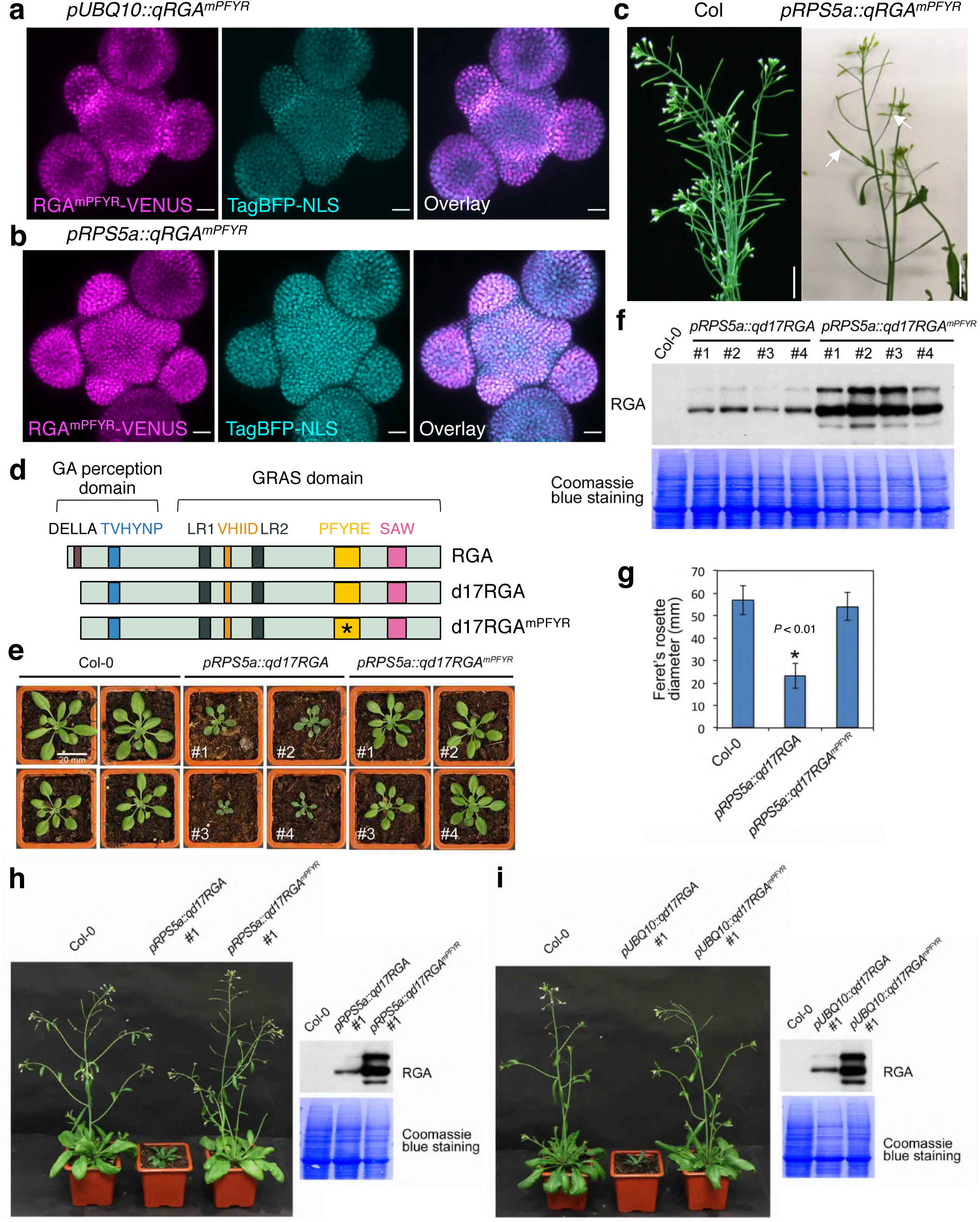
qRGA^mPFYR^ activity is not interfering with plant growth. **a-b**, Fluorescence signals of RGA^mPFYR^-VENUS, TagBFP-NLS and merge channels in *pUBQ10::qRGA^mPFYR^* (a) and *pRPS5a::qRGA^mPFYR^* (b) inflorescence SAM. **c**, Representative growing wild-type (Col-0) and *pRPS5a::qRGA^mPFYR^*plants with flowers and elongated siliques indicative of normal fertility (arrows). **d**, Schematic representation of d17RGA, a mutant version of RGA, deleted of its N-terminal DELLA domain involved in the contact with GID1, rendering the protein insensitive to GA. An asterisk indicates the mPFYR mutation. **e,f**, Representative 3 weeks old T1 independent transgenic *pRPS5a::qd17RGA* and *pRPS5a::qd17RGA^mPFYR^*plants and Col-0 controls (**e**) and immunodetection of d17RGA- VENUS and d17RGA^mPFYR^-VENUS protein levels using RGA antibodies (**f**). **g**, Feret’s rosette diameter (mean ± s.d.) of 3 weeks old Col-0, *pRPS5a::qd17RGA* and *pRPS5a::qd17RGA^mPFYR^* plants. Asterisk indicates a significant difference (*p <* 0.01) using one-way ANOVA (n = 20). **h**,**i**, Representative *pRPS5a::qd17RGA* and *pRPS5a::qd17RGA^mPFYR^* (**h**) and *pUBQ10::qd17RGA* and *pUBQ10::qd17RGA^mPFYR^* (**i**) adult plants (and Col-0 controls). On the right is shown the corresponding immunodetection of d17RGA-VENUS and d17RGA^mPFYR^-VENUS protein levels using RGA antibodies.

**Extended Data Figure 3.**
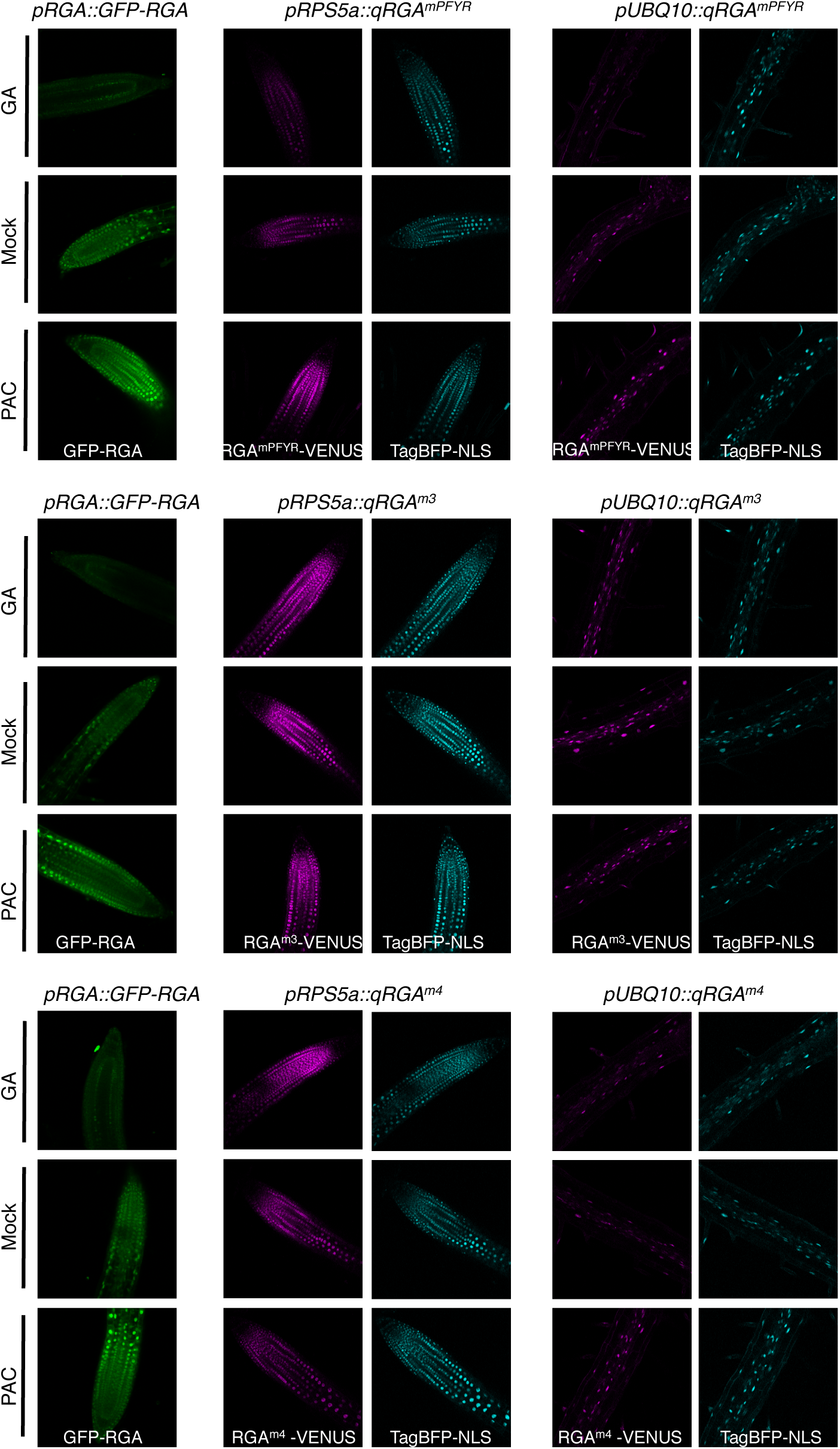
Behaviour of the different GA signalling sensor lines in response to changes in GA levels in seedling root. Representative VENUS and TagBFP signals obtained in seedling roots of qRGA^mPFYR^, qRGA^m3^ and qRGA^m4^ lines driven with *pUBQ10* and *pRPS5a* promoters, treated with 5 μM PAC or 10 μM GA_3_ (and Mock controls). *pRGA::GFP-RGA* is used as a positive control of GA and PAC treatments.

**Extended Data Figure 4.**
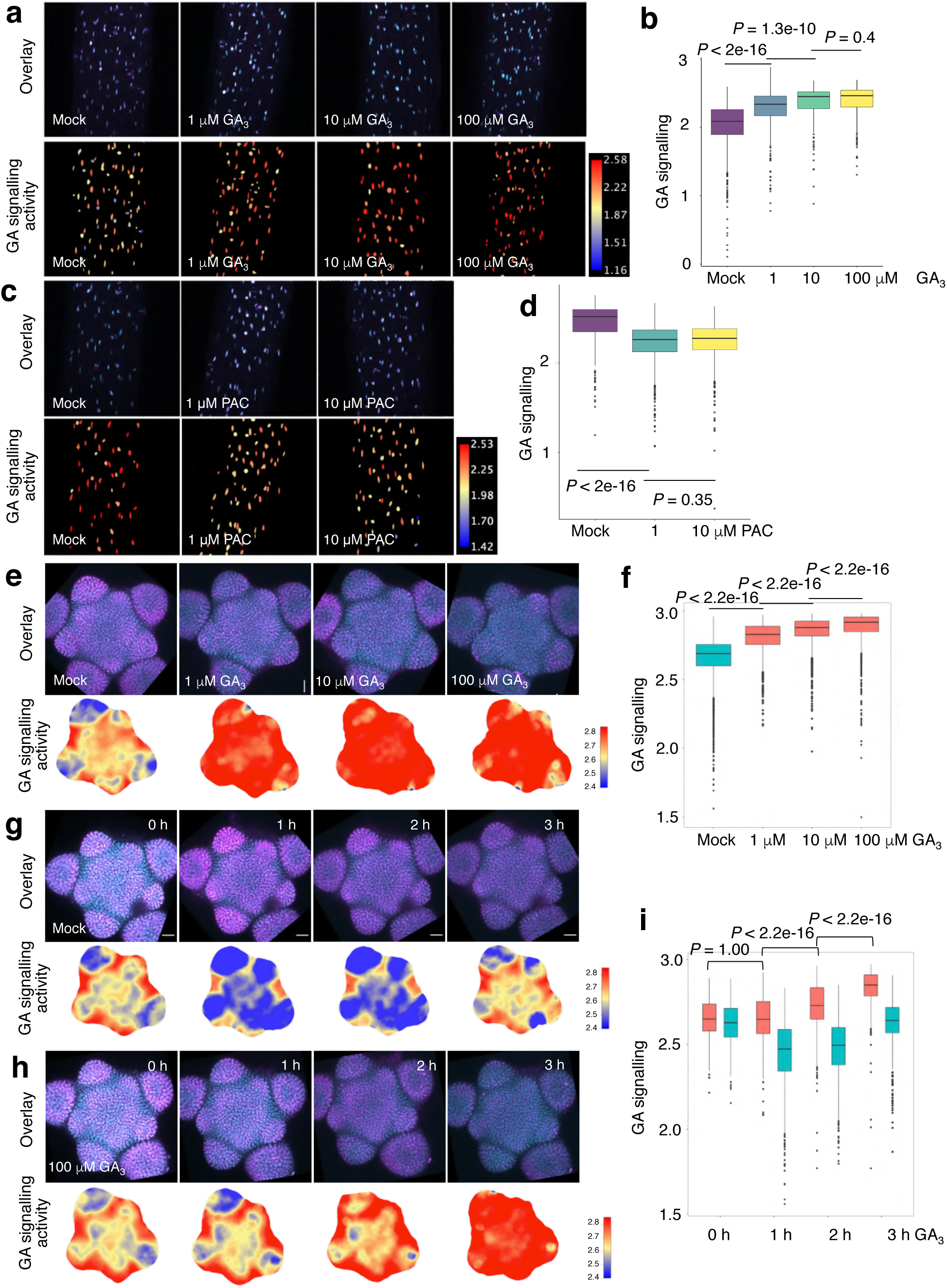
Dose and temporal responses of *qRGA^mPFYR^*to exogenous GA or PAC treatment in seedling hypocotyls and the SAM. **a-d**, Overlay of VENUS and TagBFP maximum intensity projection (upper row), and corresponding heatmap representation of GA signalling activity (lower row) in hypocotyls of 5-d-old *pUBQ10::RGA^mPFYR^* seedlings treated with 1, 10 and 100 μM GA_3_ for 90 min (**a**), or with 1 and 10 μM PAC for 24 h (**c**) and Mock controls. Boxplot representations of GA signalling activity (means of at least 8 seedlings) are shown in (**b**) and (**d**). *P*-values are from Kruskal-Wallis tests. **e-f**, Overlay of VENUS and TagBFP maximum intensity projection (upper row) and corresponding heatmap representation of GA signalling activity (lower row) of qRGA^mPFYR^ SAMs treated for 3 h with 1 μM, 10 μM and 100 μM GA_3_, and mock controls (e). Boxplot representation of GA signalling activity is shown in (f). **g-i**, Overlay of VENUS and TagBFP maximum intensity projection (g, h, upper rows) and corresponding heatmap representation of GA signalling activity (g, h, lower rows) of qRGA^mPFYR^ SAM treated with 100 μM GA_3_ for 1 h, 2 h and 3 h, and mock controls. Boxplot representation of GA signalling activity is shown in (i). *P* values are from Krustal-Wallis tests with Dunn’s all pair rank comparison test. Experiments were repeated twice with similar results. Scale bars = 20 μm.

**Extended Data Figure 5.**
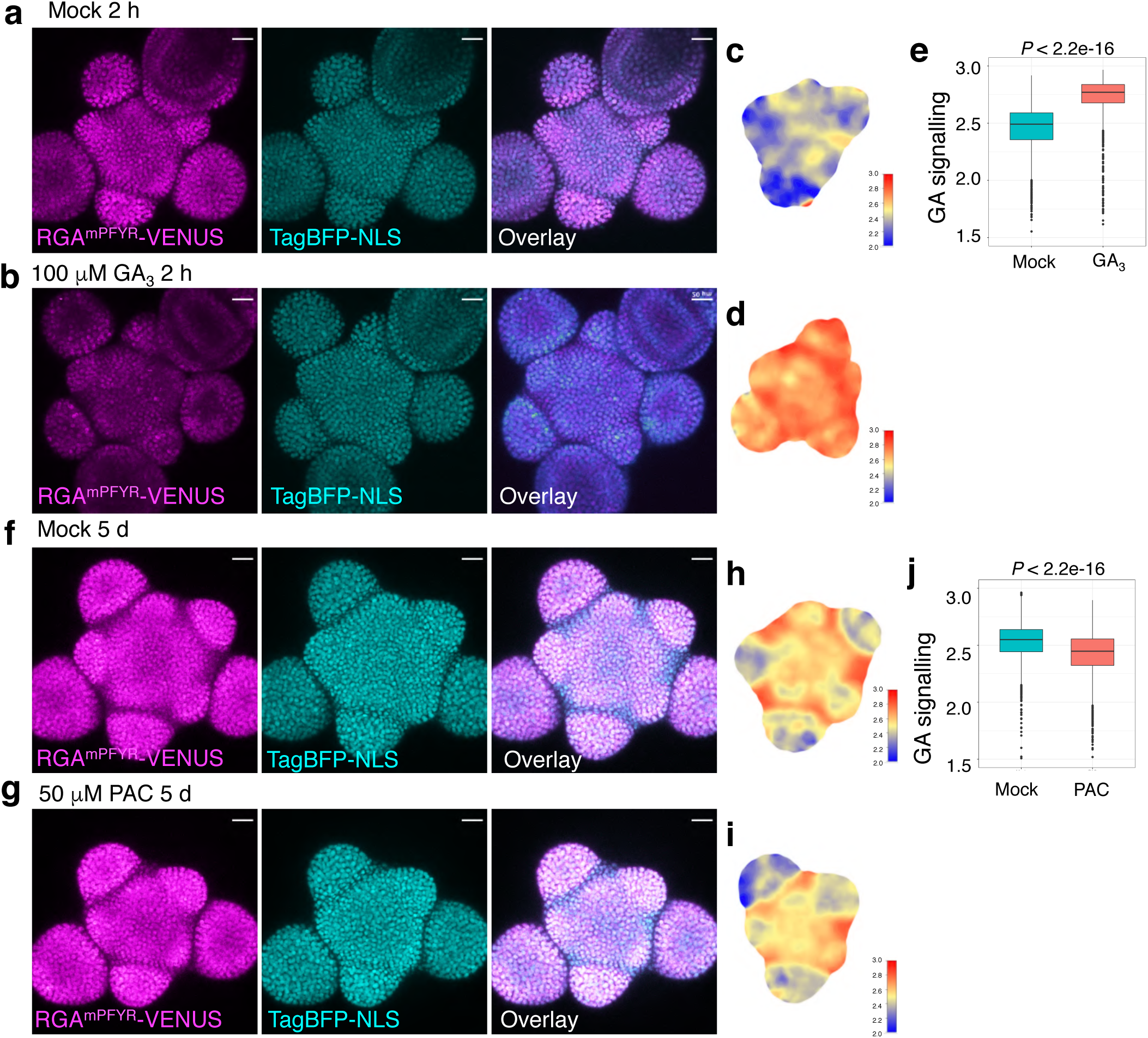
Responses of *pRPS5a::qRGA^mPFYR^*to exogenous GA or PAC treatment in the SAM. **a-e**, Fluorescence signals of RGA^mPFYR^-VENUS, TagBFP-NLS and overlay in *pRPS5a::qRGA^mPFYR^* SAM, 2h after a treatment with 100 μM GA_3_ (b) and mock controls (a). (c, d) Heatmap representations of GA signalling activity. The older floral primordia were removed digitally. (e) Boxplot representations of GA signalling activity in SAM. **f-i**, Fluorescence signals of RGA^mFPYR^-VENUS, TagBFP-NLS and overlay in *pRPS5a::qRGA^mPFYR^*SAM, 5 days after a treatment with 50 μM PAC (g) and mock (f). (h, i) Heatmap representations of GA signalling activities from (s, x). Older floral primordia were removed digitally. (u) Boxplot representation of GA signalling activity in SAM. *P* values are from Wilcoxon rank-sum tests. Experiments were repeated twice with similar results. Scale bars = 20 μm (a, b, f, g).

**Extended Data Figure 6.**
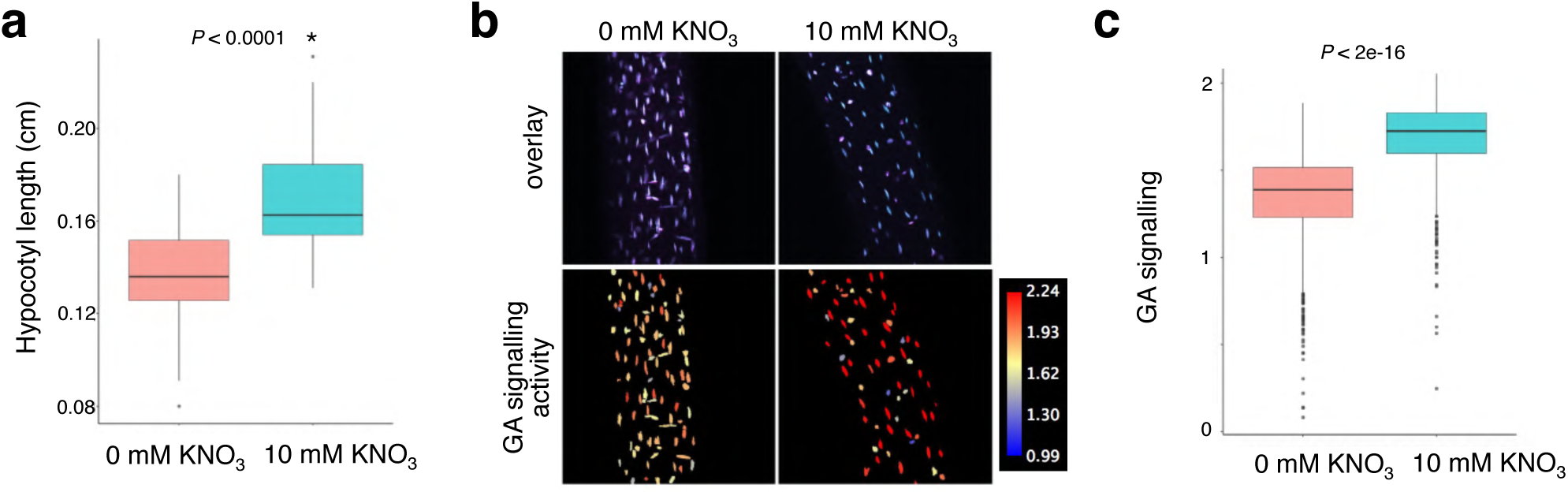
Response of qRGA^mPFYR^ to changes in endogenous GA levels in seedling hypocotyls. **a**, Hypocotyl length of 6-d-old *pUBQ10::RGA^mPFYR^* seedlings grown on nitrate-deficient conditions and on 10 mM KNO_3_. Asterisk indicates a significant difference (*p <* 0.0001) using Student’s t-test (n > 24). **b**, **c**, Overlay of VENUS and TagBFP maximum intensity projection, and corresponding heatmap representation of GA signalling activity in hypocotyls of 5-d-old *pUBQ10::RGA^mPFYR^* seedlings grown on nitrate-deficient conditions and on 10 mM KNO_3_ (**b**). Boxplot representation of GA signalling activity (means of at least 6 seedlings) is shown in (**c**). *P* value is from Wilcoxon Rank Sum Test.

**Extended Data Figure 7.**
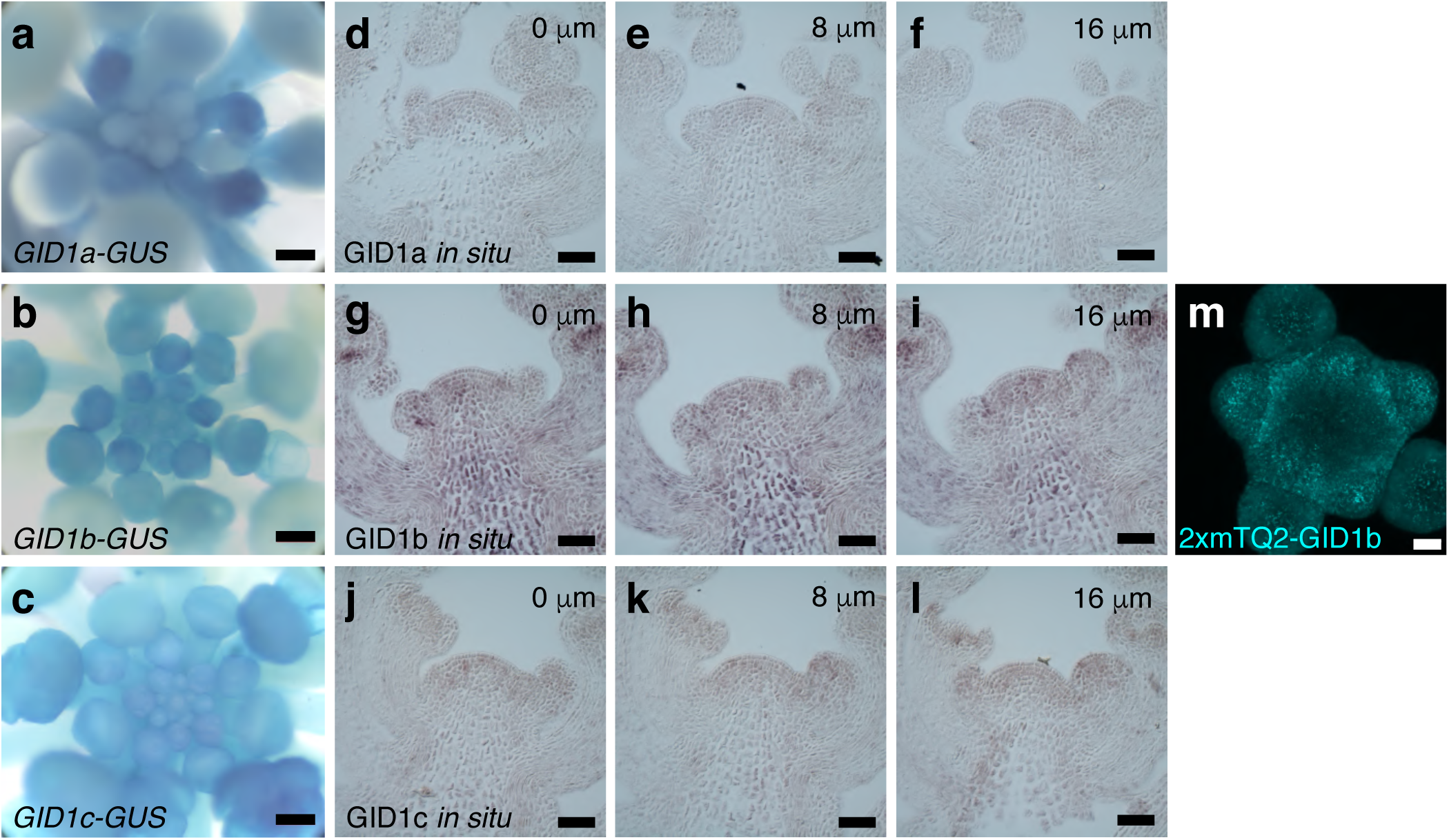
Expression patterns of GA receptors in the SAM. **a-c**, Expression patterns of *GID1a*, *GID1b* and *GID1c* in the SAM of *pGID1a::GID1a-GUS* (a), *pGID1b::GID1b-GUS* (b) and *pGID1c::GID1c-GUS* (c) adult plants. **d-l**, *In situ* localization of *GID1a* (d-f), *GID1b* (g-i) and *GID1c* (j-l) mRNA in inflorescence apex. 3 consecutive sections are shown for each. **m**, Maximum intensity projection showing the expression of *pGID1b::2xmTQ2-GID1b*. Scale bars = 200 μm (a-c) and 20 μm (d-m).

**Extended Data Figure 8.**
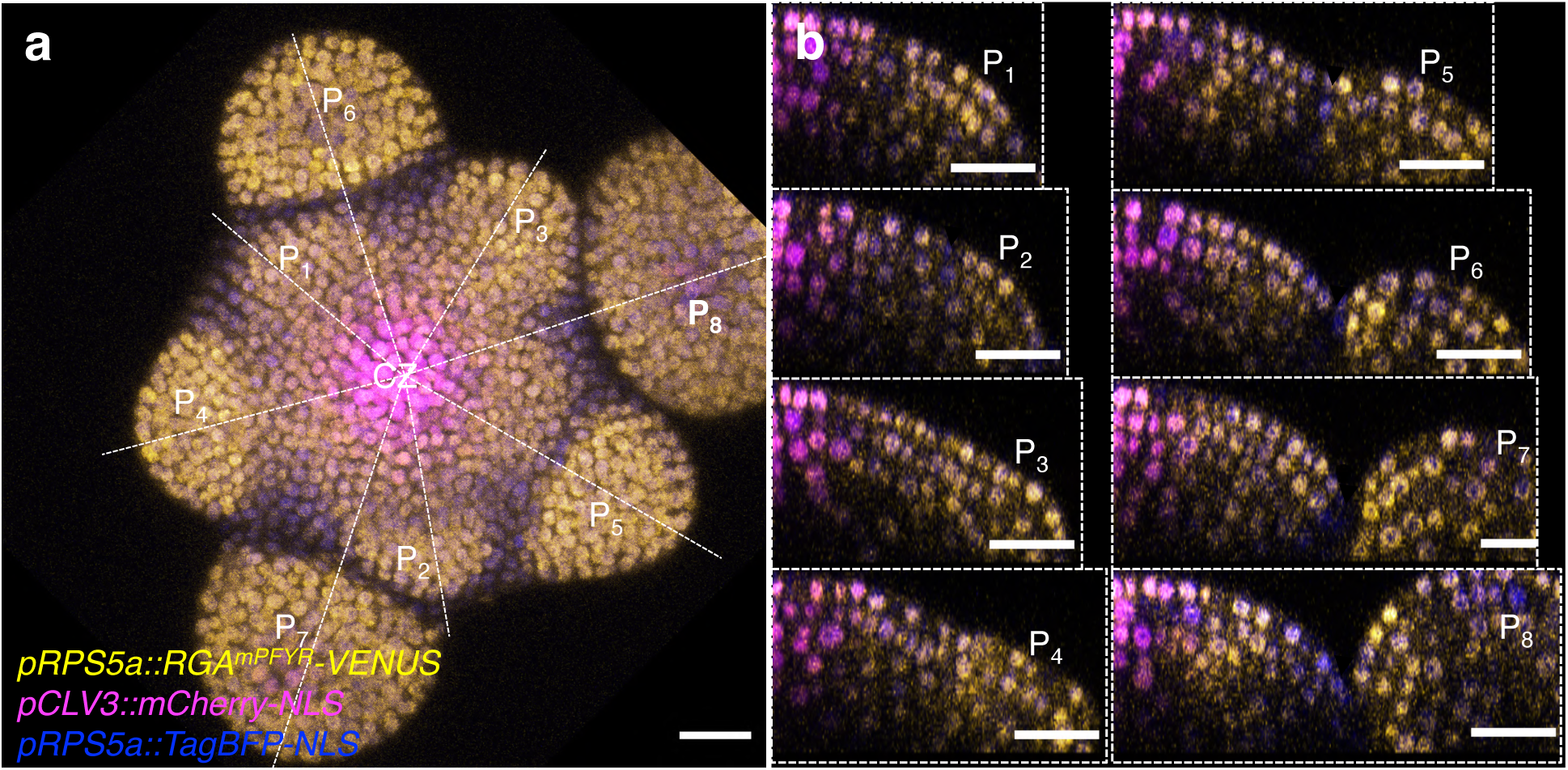
Digital longitudinal sections through the centre of the SAM and primordia of qRGA^mPFYR^ plants. **a**, Maximum intensity projection showing the expression of *pRPS5a::qRGA^mPFYR^ pCLV3::mCherry-NLS* plants in the SAM as in Fig 4a. CZ, central zone. **b**, Signals for qRGA^mPFYR^-VENUS, TagBFP-NLS and mCherry-NLS in digital longitudinal sections through the centre of CZ and primordia P_1_ to P_8_ along the white dotted lines indicated in (**a**). Scale bars = 20 μm.

**Extended Data Figure 9.**
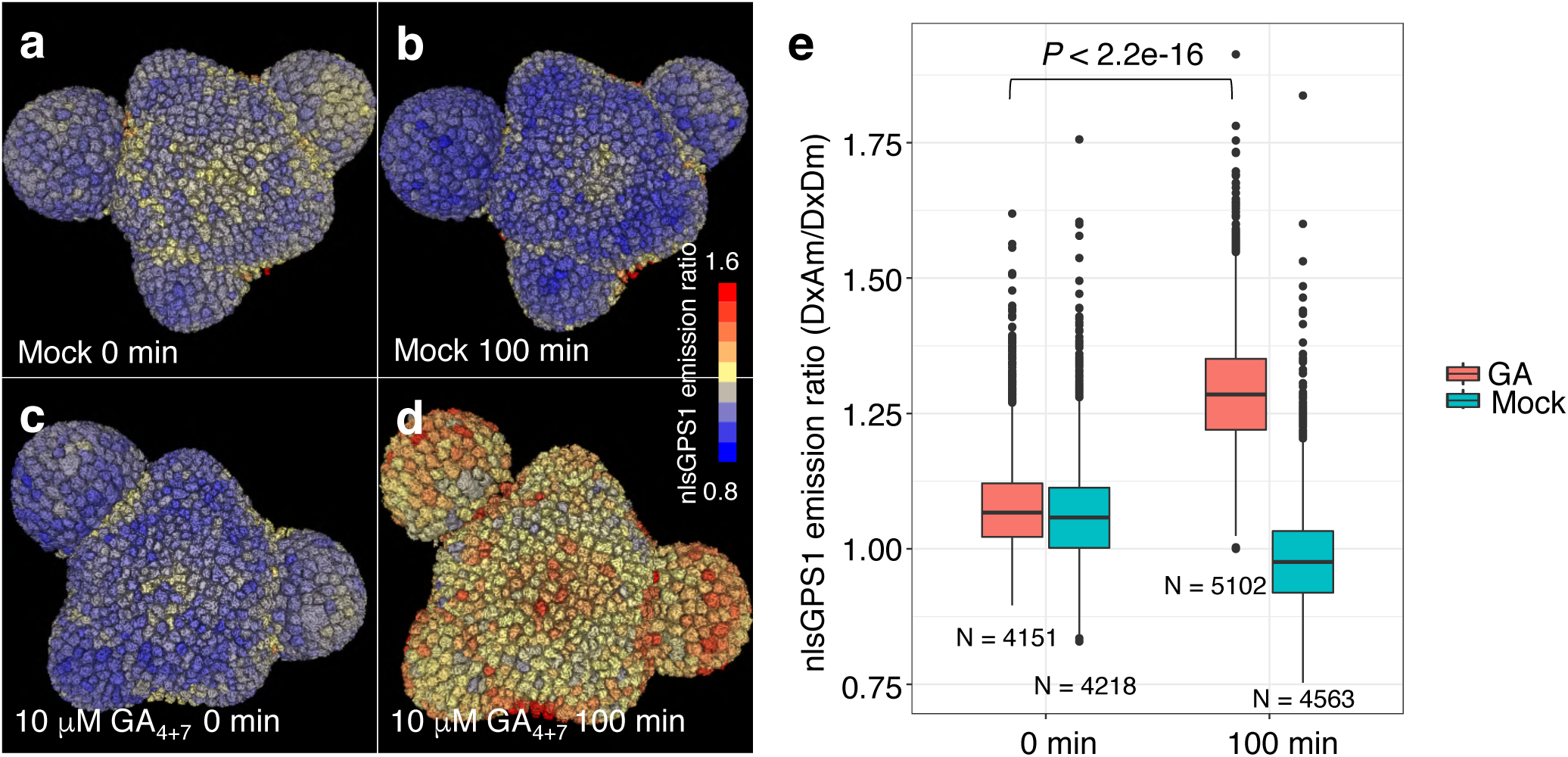
Response of nlsGPS1 to GA treatment in the SAM. **a-d**, 3D visualization of nlsGPS1 expression in the SAM before (**a**, **c**) and after (**b**, **d**) mock (**a**, **b**) and 10 μM GA_4+7_ treatment (**c**, **d**). **e**, Boxplot representation of nlsGPS1 emission ratios from nuclei analysed in three SAMs. The number of nuclei (N) is indicated. *P* value is from one-way ANOVA with Turkey’s test for multiple comparisons. The experiment was repeated twice with similar results.

**Extended Data Figure 10.**
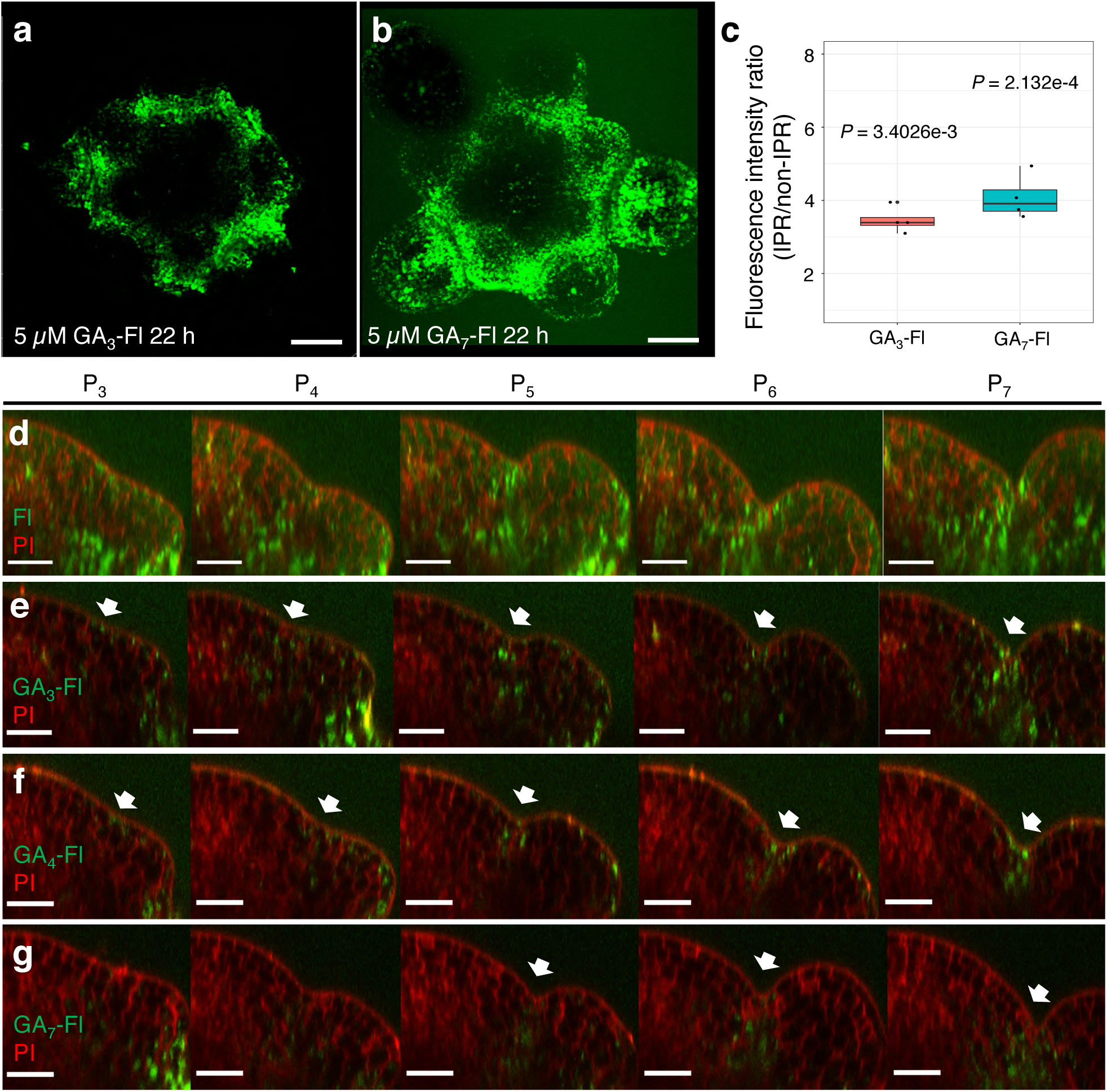
Fluorescence distribution in SAMs treated with Fl and GA_3_-, GA_4_- and GA_7_-Fl. **a-b**, Fluorescence distribution in wild-type (L*er*) SAMs treated with GA_3_-Fl (**a**) and GA_7_-Fl (**b**). **c**, Ratio of average fluorescence intensity in the IPR to that in the non-IPR (excluding primordia) after GA_3_-Fl and GA_7_-Fl treatment in the SAM, compared to Fl treatment (Fig 4f, h). *P* values are from Wilcoxon rank-sum tests. **d-g,** Digital longitudinal sections through the centre of CZ and primordia P_3_ to P_7_ of L*er* SAMs treated with 5 μM fluorescein (**d**), GA_3_-Fl (**e**), GA_4_-Fl (**f**), or GA_7_-Fl (**g**) (green) and stained with PI (red). Scale bars = 20 μm.

**Extended Data Figure 11.**
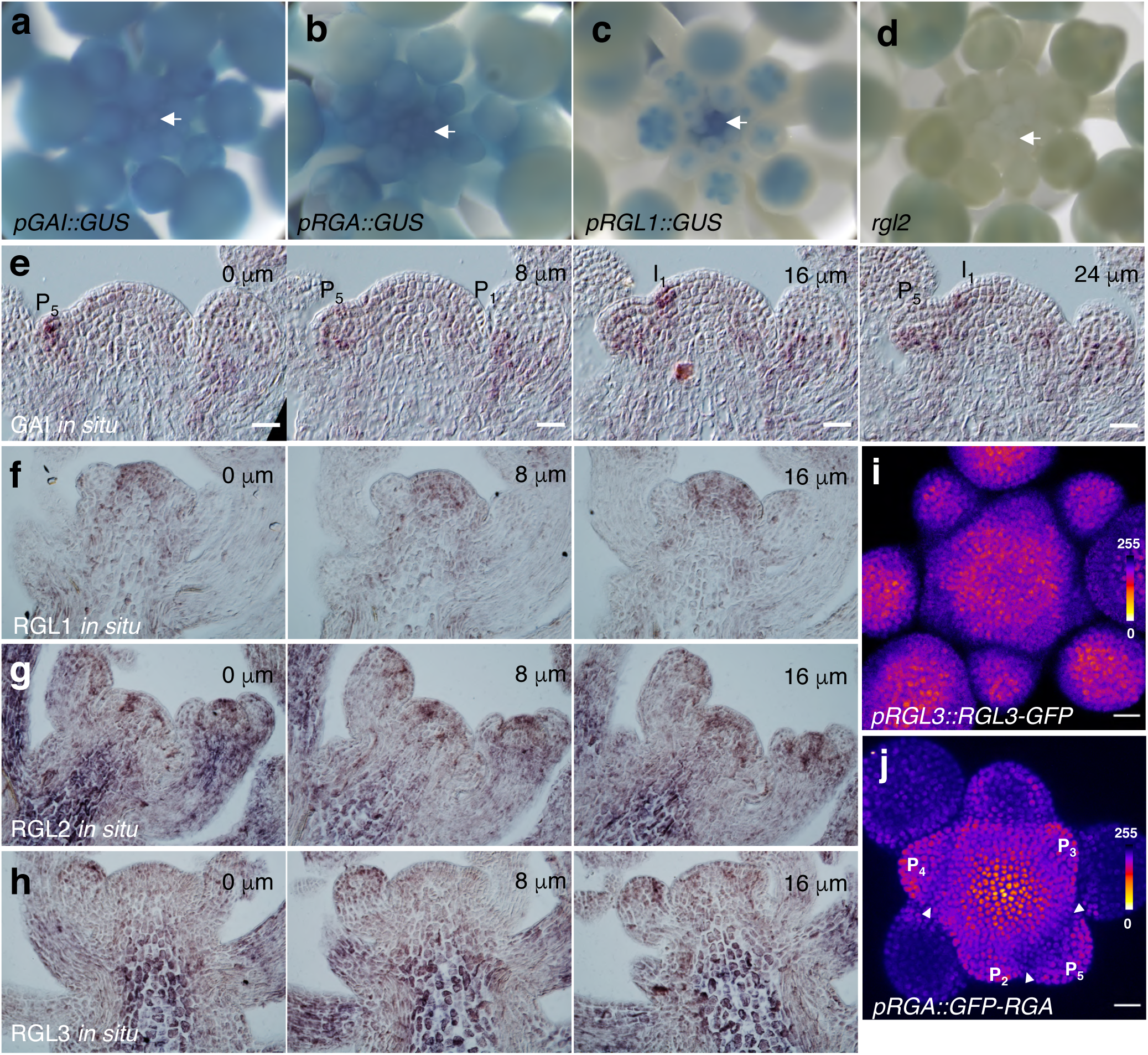
Expression patterns of DELLA-encoding genes and accumulation DELLA proteins in the SAM. **a-d**, GUS signal in the shoot apex of *pGAI::GUS* (**a**), *pRGA::GUS* (**b**), *pRGL1::GUS* (**c**) and *rgl2-5* (a promoter trap GUS line) mutant (**d**). **e**-**h**, *In situ* hybridization serial sections through a shoot apex using probes of GAI (**e**), RGL1 (**f**), RGL2 (**g**) and RGL3 (**h**). **i-j,** Intensity heatmaps of maximum intensity-projection of *pRGL3::RGL3-GFP* (**i**) and *pRGA::GFP-RGA* (**j**) in the SAM. Arrowheads mark the regions with lower GFP-RGA expression. Scale bars = 20 μm.

**Extended Data Figure 12.**
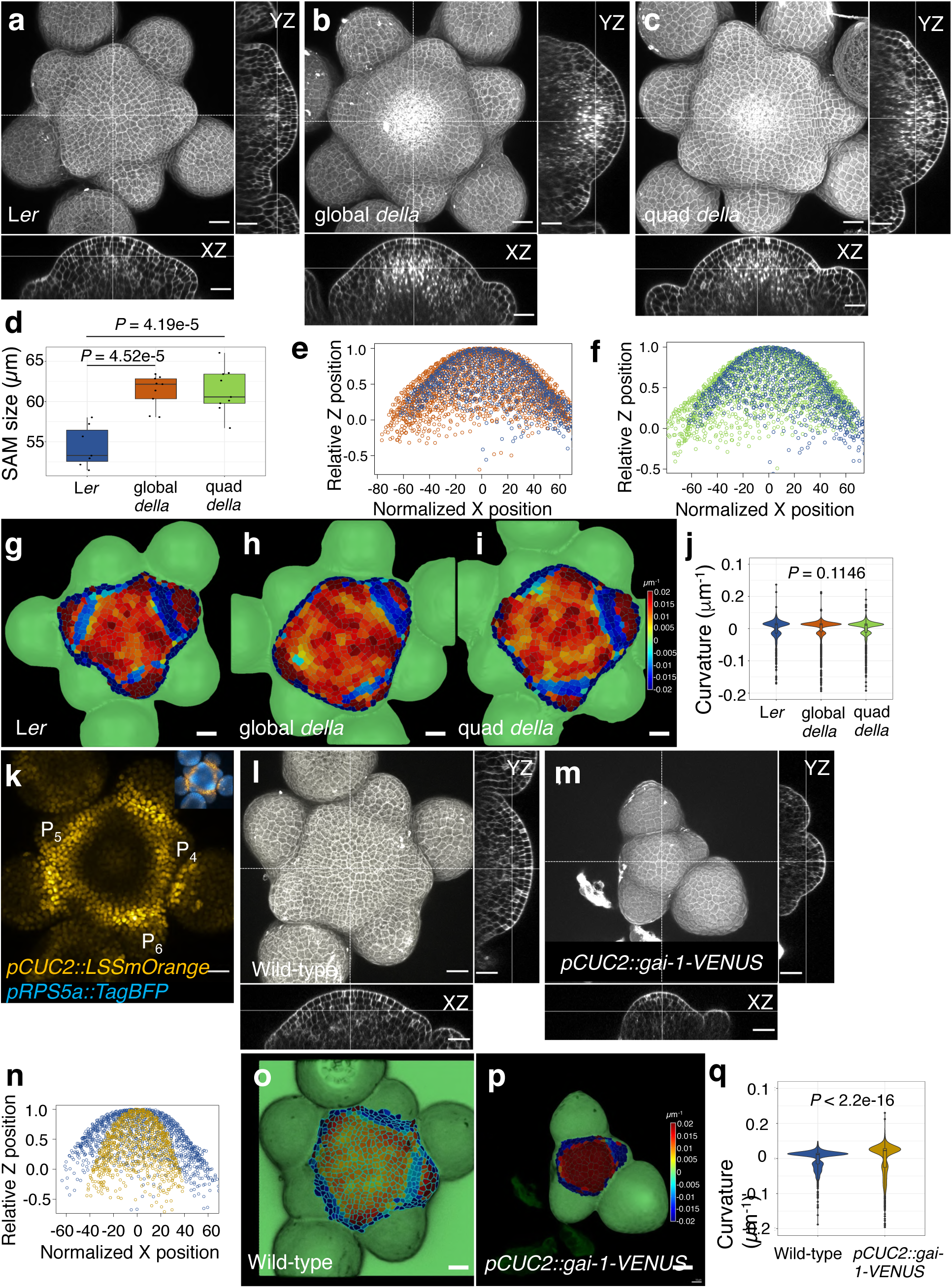
Analysis of SAM shape of global and quadruple *della* mutants and *pCUC2::gai-1-VENUS* transgenic plants. a-c, Maximum projection (**a-c**) and orthogonal views (XZ dimension, lower panels; YZ dimension, left panels) of PI-stained SAMs of L*er* (**a**), global *della* (**b**) and quadruple *della* mutant (**c**) using confocal microscopy. **d**, Comparison of SAM size between L*er*, global and quadruple *della* mutants. *P* values are from one-way ANOVA with Tukey’s multiple comparison. **e**,**f**, XZ point plots of L1 nuclei of L*er* (blue), global *della* (orange) and quadruple *della* (green) showing the tissue-level curvature of the SAM. **g-j**, Heatmaps of cellular curvature of L1 cells in the SAM (including P_4_) of L*er* (**g**), global *della* (**h**) and quadruple *della* (**i**). Statistics are shown in (**j**). *P* values are from Kruskal-Wallis tests. **k**, Maximum projection view of the SAM of *pCUC2::LSSmOrange-NLS* line. The insert shows the same image superimposed with the expression of *pRPS5a::TagBFP-NLS*. **l-m**, Maximum projection and orthogonal views (XZ dimension, lower panels; YZ dimension, left panels) of PI-stained SAM of *pCUC2::gai-1-VENUS* (**m**) transgenic plant and its wild-type control (**l**). **n**, XZ point plots of L1 nuclei of wild-type (blue) and *pCUC2::gai-1-VENUS* transgenic plants (yellow) showing the tissue-level curvature of the SAM. **o-q**, Heatmaps of cellular curvature of L1 cells in the SAM of wild-type (**o**) and *pCUC2::gai-1-VENUS* (**p**) transgenic plants. Statistics are shown in (**q**). *P* value is from Wilcoxon rank-sum test. Scale bars = 20 μm.

**Extended Data Figure 13.**
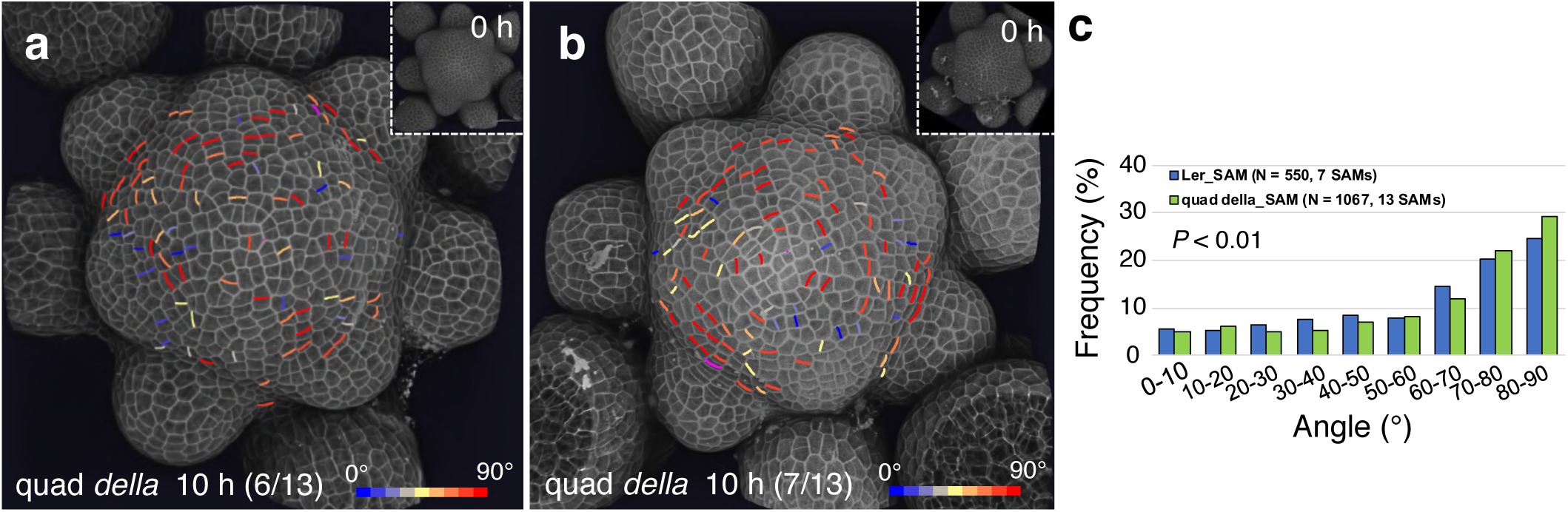
Cell division plane orientation distribution in the SAM of quadruple *della* mutants. **a-b**, 3D visualization of the L1 of PI-stained SAM of quadruple *della* mutant using confocal microscopy. New cell walls formed in the SAM (but not in primordia) in 10 h are shown and coloured according to their angle values. (**a**) Representative results of 6 out of 13 SAMs, and (**b**) of the remaining 7 SAMs where a higher frequency of transverse cell divisions could be detected. Inserts are images at 0 h. (**c**) Comparison of frequency distributions of division plane orientation between cells located in the entire SAM of L*er* and quadruple *della* mutants. *P* value is from Kolmogorov-Smirnov test.

**Extended Data Figure 14.**
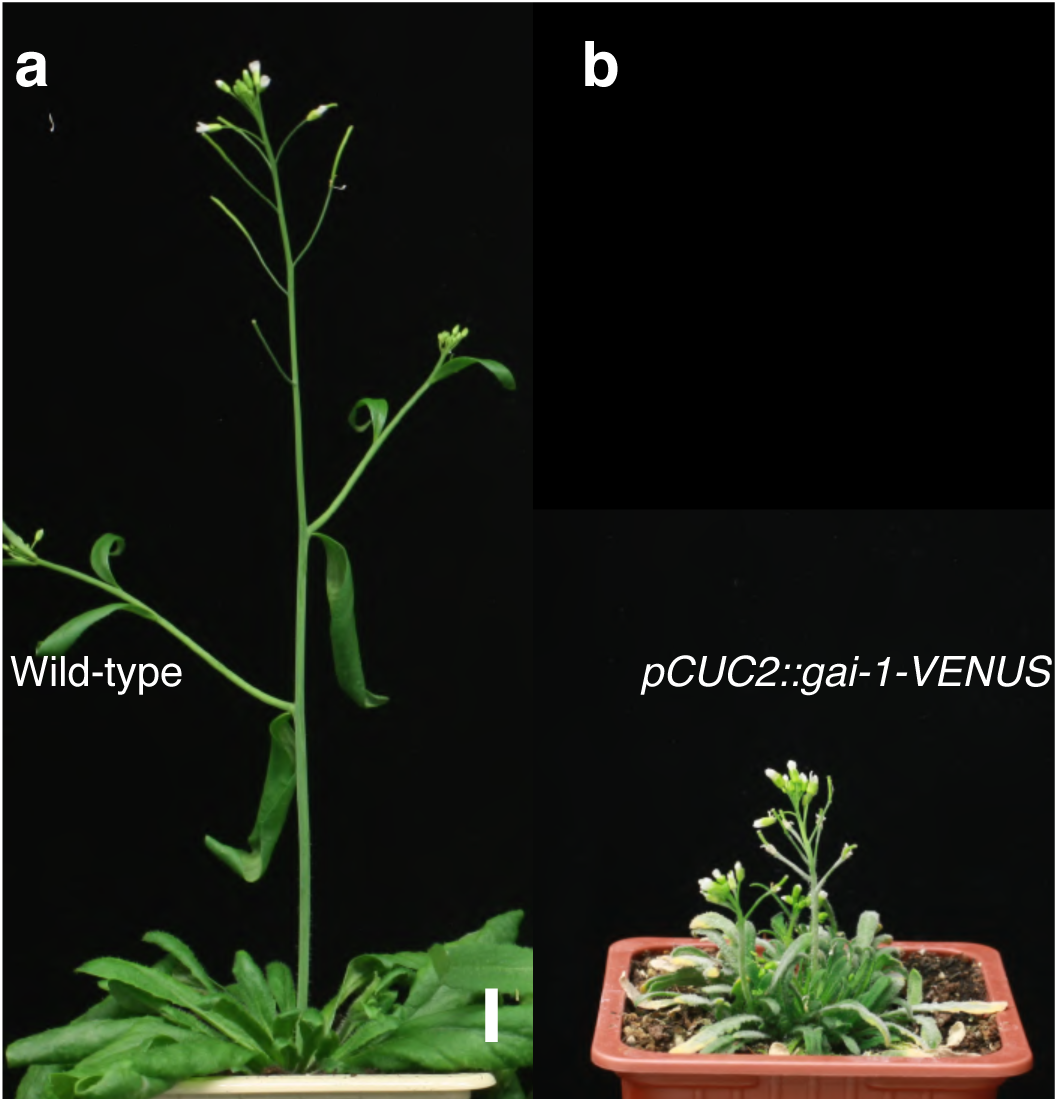
Altered plant height in *pCUC2::gai-1-VENUS* transgenic plants. a-b, Representative 37-d-old wild-type (**a**) and 56-d-old *pCUC2::gai-1-VENUS* transgenic plants (**b**) grown under long-day conditions. Scale bar = 1 cm.

**Extended Data Figure 15.**
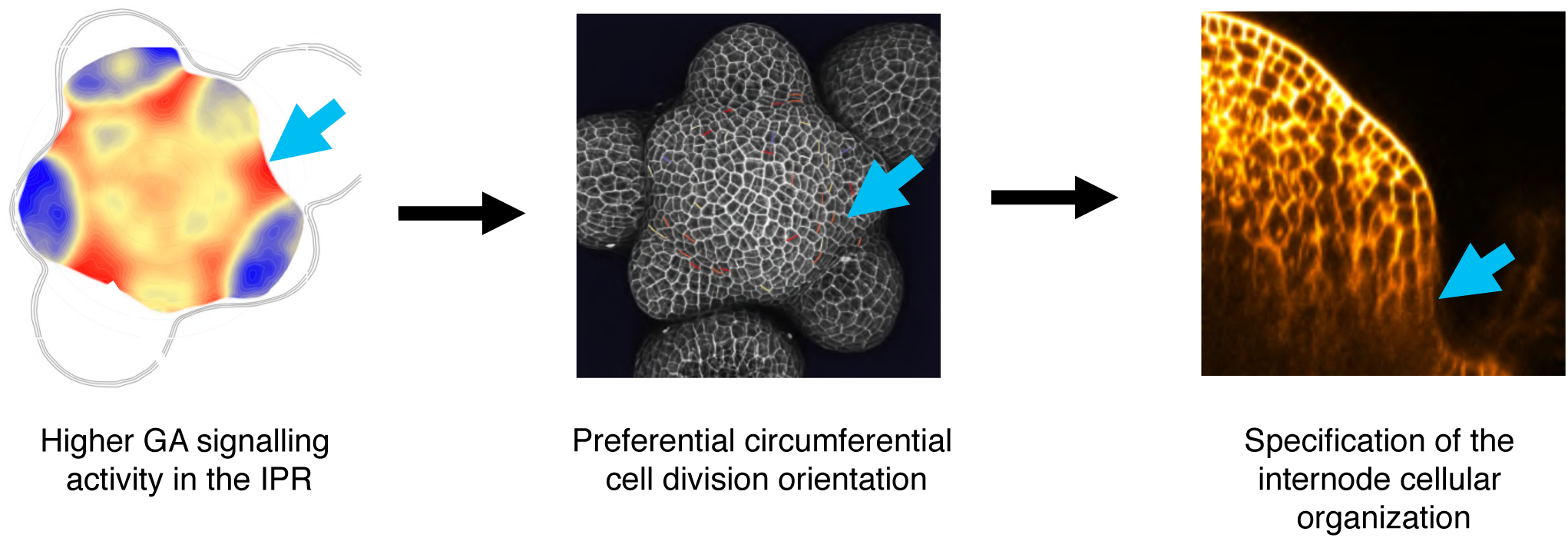
Hypothesis of the role of GA signalling in internode specification. Higher levels of GA signalling activity in the IPR specify the cellular organization of internodes in the SAM via the regulation of the orientation of cell division.

**Supplementary Table 1. RGA^m2^ loses its ability to bind with RGA-interacting partners.** Extended pairwise Y2H interaction assays between RGA, RGA^mPFYR^ and RGA-interacting transcription factors. The colour code and numbers (0 to 5) indicate the strength of the interactions based on the size of the yeast colonies on selective media without leucine, tryptophan and adenine (-LWH) without or with 1 mM 3-aminotriazole (3AT). Empty pGBKT7 and pGADT7 vectors were included as negative controls.

## Acknowledgments

We thank the colleagues of the Laboratoire Reproduction et Développement des Plantes for fruitful discussions on this work and for sharing material and advice. We also thank Miguel Perez-Amador, Jan Lohmann, Yvon Jaillais, Tai-ping Sun, Miguel Blazquez, David Alabadi and the NASC for seeds and plasmids. We acknowledge the contribution of SFR Biosciences (UMS3444/CNRS, US8/Inserm, ENS de Lyon, UCBL) facility PLATIM for assistance with microscopy and image analysis. This work was supported by the ANR-16-CE13-0014 (GrowthDynamics) grant to T.V. and P.A.; a European Research Council grant (GAtransport) to R.W.; a Guangdong Laboratory for Lingnan Modern Agriculture Grant NG2021001 to B.S..

## Author contribution

P.A. and T.V. designed the study and supervised the work; B.S., A.F.-B., P.A. and T.V. designed the experiments. B.S., A.W. and A.M. performed the FRET analysis; A.N.-G. and S.P. conducted the large-scale two-hybrid analysis; S.L. and R.W. designed the synthesis strategy and synthesized the GA-Fl fluorescent probes. G.C. developed the pipeline for quantitative image analysis with the help of J.L.; B.S., A.F.-B., J.M. and G.C. performed the image analysis. B.S., A.F.-B., C.G.-A., L.J., G.B., E.V., L.S.-A. and J.-M.D. performed all the other experiments; all authors were involved in data analysis; B.S., A.F.-B., P.A. and T.V. wrote the manuscript with inputs from all authors.

## Corresponding authors

Correspondence and request for materials should be addressed to T.V. or P.A.

