## Supplementary Method 1 for "A quantitative gibberellin signalling biosensor reveals a role for gibberellins in internode specification at the shoot apical meristem"

Supplementary Methods

### 1 Quantitative analysis of microscopy images

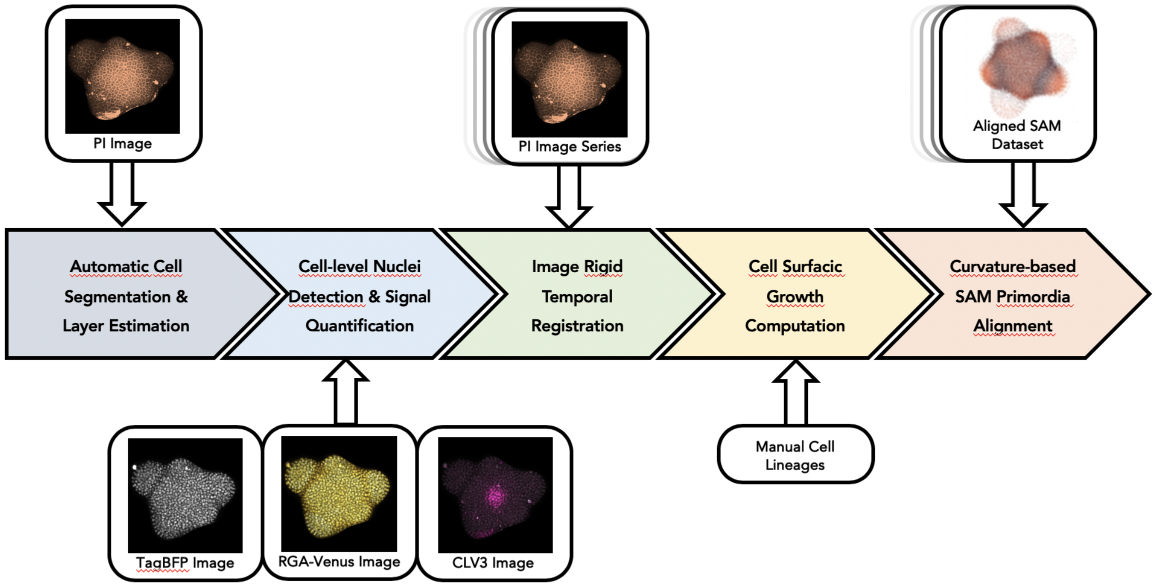


Supplementary Figure 1: **Automatic quantification pipeline for the time-lapse microscopy images.** To obtain quantitative data from the images produced under the microscope, various sequential processing steps need to be performed, from the extraction of the relevant objects (cells regions based on a cell wall marker channel, nuclei positions with their different channel intensity values) to the geometrical characterization and the spatio-temporal registration of the tissues, to finally get a complete, aligned and consistent dataset gathering all the imaged meristems.

The images obtained from confocal microscopy are processed using a complex computational pipeline derived from the one introduced in [1]. It involves several steps of image analysis, computational geometry and data manipulation that allow going from raw 3D images to aligned quantified data points (Supplementary Figure 1). In the following, we describe the methods used for these different steps, highlighting when new algorithms were developed with respect to the original methods from [1].

#### 1.1 Automatic Cell Segmentation & Layer Estimation

##### 1.1.1 Image Notations

We consider that a 3D image consists of an array $I$ of size $K_x \times K_y \times K_z$ filled with values taken in an integer intensity interval $\mathcal{I} \subset \mathbb{N}$. The elements of this 3D array are called voxels. In the case of a 16-bit-encoded unsigned integer image, the intensity interval $\mathcal{I} = [\![0, 2^{16}[\![$. We denote:

$$I = \left\{ I_{ijk} \in \mathcal{I} \;\middle|\; (i,j,k) \in [\![0, K_x[\![ \times [\![0, K_y[\![ \times [\![0, K_z[\![ \right\}. \quad (1)$$

The voxels in the array grid can be projected into a physical space $\Omega \subset \mathbb{R}^3$ through a mapping function $\mathbf{x} : [\![0, K_x[\![ \times [\![0, K_y[\![ \times [\![0, K_z[\![ \to \Omega$ that associates the array indices with a discrete, evenly spaced, 3D lattice $\Omega_I \subset \Omega$. The voxel size $v = (v_x, v_y, v_z) \in \mathbb{R}^3$ defines a spacing of the lattice that is potentially different on each dimension:

$$\Omega_I = \left\{\mathbf{x}(i,j,k) = (i \cdot v_x, j \cdot v_y, k \cdot v_z) \;\middle|\; (i,j,k) \in [\![0, K_x[\![ \times [\![0, K_y[\![ \times [\![0, K_z[\![\right\}. \tag{2}$$

This mapping allows to define the image as a function $I : \Omega_I \rightarrow \mathcal{I}$ that associates a 3D point $\mathbf{x} = (x, y, z)$ in the image definition space $\Omega_I$ with a fluorescence intensity value $I(\mathbf{x}) \in \mathcal{I}$, such that $\forall\, (i,j,k) \in [\![0, K_x[\![ \times [\![0, K_y[\![ \times [\![0, K_z[\![$,

$$I\left(\mathbf{x}(i,j,k)\right) = I_{ijk}. \tag{3}$$

In the case of a multichannel image, we denote $I_S$ the image channel corresponding to the signal $S$. Cells are segmented for each meristem acquisition independently using the *Propidium Iodide* (PI) channel, which we denote $I_{\mathrm{PI}}$.

###### 1.1.2 Automatic Cell Segmentation

As described in [1], the image segmentation method we use is an auto-seeded 3D watershed algorithm derived from the MARS pipeline [2] applied to the $I_{\mathrm{PI}}$ image (Supplementary Figure 2 a). The parameters are the following:

- The standard deviation of the Gaussian filter used for seed detection is $0.75\mu m$
- The $h$ value used for the H-min transform is either 200 (for 16-bit images) or 2 (for 8-bit images)
- The standard deviation of the Gaussian filter used for watershed is $0.5\mu m$

In the end we obtain a segmented image $I_{\mathrm{seg}}$ that assigns an integer label to every voxel of the image grid $\Omega_I$ on which the image $I_{\mathrm{PI}}$ to segment is defined (Supplementary Figure 2 b). The cells of the tissue are represented by independent connected regions of voxels (so that the same label can not be assigned to voxels that are not part of the same connected component of $I_{\mathrm{seg}}$). The background corresponds to a specific label, that is systematically set to 1 to ensure consistency between images. Each cell labeled $c \in \mathcal{C} = [\![1, K_c]\!]$ is the then represented by a connected region $\Gamma_c$ so that:

$$\Gamma_c = \{\mathbf{x} \in \Omega_I \mid I_{\mathrm{seg}}(\mathbf{x}) = c\} \tag{4}$$

###### 1.1.3 Tissue Surface Extraction

To determine cell layers and to estimate the curvature of L1 cells, the surface of the tissue is computed based on the $I_{\mathrm{seg}}$ image as a 3D triangle mesh $\mathcal{M} = \{\mathcal{V}, \mathcal{T}\}$, where:

- $\mathcal{V}$ is a set of vertices
- Each vertex $v \in \mathcal{V}$ is associated with a 3D position $M_v \in \mathbb{R}^3$
- $\mathcal{T}$ a set of triangles defined by triplets of indices $(v_1, v_2, v_3) \in \mathcal{V}^3$
- $\mathcal{T}$ is such that the resulting simplicial complex forms a 2-manifold [3] (Chapter 5.3: Topological Spaces, Chapter 6.3: Simplicial Complexes).

To obtain this triangle mesh, we use the binary mask $I_{\mathrm{bin}}$ of the segmented image $I_{\mathrm{seg}}$ that associates the values 0 to background voxels and 1 to tissue voxels:

$$\forall \mathbf{x} \in \Omega_I, I_{\mathrm{bin}}(\mathbf{x}) = \begin{cases} 0 & \text{if} \quad I_{\mathrm{seg}}(\mathbf{x}) = 1 \\ 1 & \text{otherwise.} \end{cases} \tag{5}$$

This binary image is meshed by applying a Marching Cubes algorithm [4] on a resampled version of the image, to ensure a cubic shape of the image voxels. This triangular mesh undergoes a phase of triangle decimation [5] and isotropic remeshing [6] to obtain a surface composed of roughly 50000 regular faces.

###### 1.1.4 Topological Relationships between Cells

The notion of adjacency between cells in the segmented image $I_{\mathrm{seg}}$ is directly related to the notion of adjacency between the voxels of the underlying 3D matrix. There are 3 possible conceptions of voxel adjacency in 3D images, which can be summed up by:

- In *6-connectivity*, 2 voxels are adjacent if they share a common face, called a *surfel*

- In *18-connectivity*, 4 voxels are adjacent if they share a common edge, called a *linel*
- In *26-connectivity*, 8 voxels are adjacent if they share a common vertex, called a *pointel*

Note that each of these *topological elements* (of dimension 2, 1 and 0 respectively) can be associated with a 3D position, defined as the barycenter of the voxels that share it. For instance a *surfel* shared by voxels $\mathbf{x}$ and $\mathbf{x}'$ is represented by the 3D point $\frac{1}{2}(\mathbf{x} + \mathbf{x}')$.

We use this notion to define sets of *topological elements* that constitute the interface between cells. Depending on the number of cells considered (2, 3 or 4), those sets will consist repectively of *surfels*, *linels* or *pointels*. More precisely, we define:

- $\Gamma^2_{c,c'}$ as the set of 3D positions of *surfels* shared by voxels labelled $c$ and $c'$
- $\Gamma^1_{c,c',c''}$ as the set of 3D positions of *linels* shared by voxels labelled $c$, $c'$ and $c''$
- $\Gamma^0_{c,c',c'',c'''}$ as the set of 3D positions of *pointels* shared by voxels labelled $c$, $c'$, $c''$ and $c'''$

With this framework, we consider that two cells are adjacent if their interface (composed of surfels) is larger than a minimal amount $n_{\min}$, so that $\forall c, c' \in \mathcal{C}$ :

$$c \text{ is } \textit{adjacent} \text{ to } c' \iff \left|\Gamma^2_{c,c'}\right| \geq n_{\min} \tag{6}$$

This adjacency definition is obviously symmetrical, and we denote $N(c)$ the set of cell labels $c'$ such that $c'$ is *adjacent* to $c$. Therefore $c' \in N(c) \iff c \in N(c')$. Practically, we used the value $n_{\min} = 8$ to determine cell adjacency.

##### 1.1.5 Cell Layer Estimation

For the purpose of the analysis, we want to discriminate between the first layer of cells (L1) and the rest of the tissue. The most straightforward way to define L1 cells in segmented images is to consider the set $N(1)$ of cells that are adjacent to the background region. However, due to artifacts of the segmentation method, this criterion alone might cause cells from deeper layers to be wrongly identified as L1.

To overcome this issue, we use an additional criterion which is the proximity of the cell centers to the surface of the tissue, represented by the triangular surface mesh $\mathcal{M} = \{\mathcal{V}, \mathcal{T}\}$. The center $C_c$ of the cell labelled $c$ can be computed as:

$$C_c = \frac{1}{|\Gamma_c|} \sum_{\mathbf{x} \in \Gamma_c} \mathbf{x}. \tag{7}$$

For each cell labelled $c$, we identify the mesh vertex $v(c)$ that achieves the minimal distance to the cell center $C_c$: $\forall c \in \mathcal{C} \setminus \{1\}$,

$$v(c) = \underset{v \in \mathcal{V}}{\operatorname{argmin}} \left(\|C_c - M_v\|\right). \tag{8}$$

We define $\mathcal{L}_1$ as the subset of $\mathcal{C}$ formed by indices $c$ of cells that are *closer* to the surface $\mathcal{M}$ than a distance threshold $d_{\max}$, while being adjacent to the background region in the segmented image. In other terms, $\forall c \in \mathcal{C} \setminus \{1\}$:

$$c \in \mathcal{L}_1 \iff \begin{cases} c \in N(1) \\ \|C_c - M_{v(c)}\| \leq d_{\max}. \end{cases} \tag{9}$$

In images of the shoot apex meristem, we use a maximal distance value of $d_{\max} = 7\mu m$, that roughly corresponds to the typical cell diameter in this tissue.

##### 1.1.6 L1 Cell Curvature Estimation

By computing normal vectors on every vertex $v$ of the surface mesh $\mathcal{M}$, it is possible to estimate the local curvature parameters of the surface [7]. We use this method compute the values of principal curvatures at the vertex $v$, denoted $\kappa^-(v)$ for the minimum curvature value and $\kappa^+(v)$ for the maximum curvature value. From these, we can derive the mean curvature value $\kappa(v) = \frac{1}{2}(\kappa^+(v) + \kappa^-(v))$.

As a first approximation, we considered that the surface curvature of a L1 cell $c$ could be measured by the curvature of the surface mesh $\mathcal{M}$ at its closest vertex $v(c)$. We estimate all L1 cell curvature

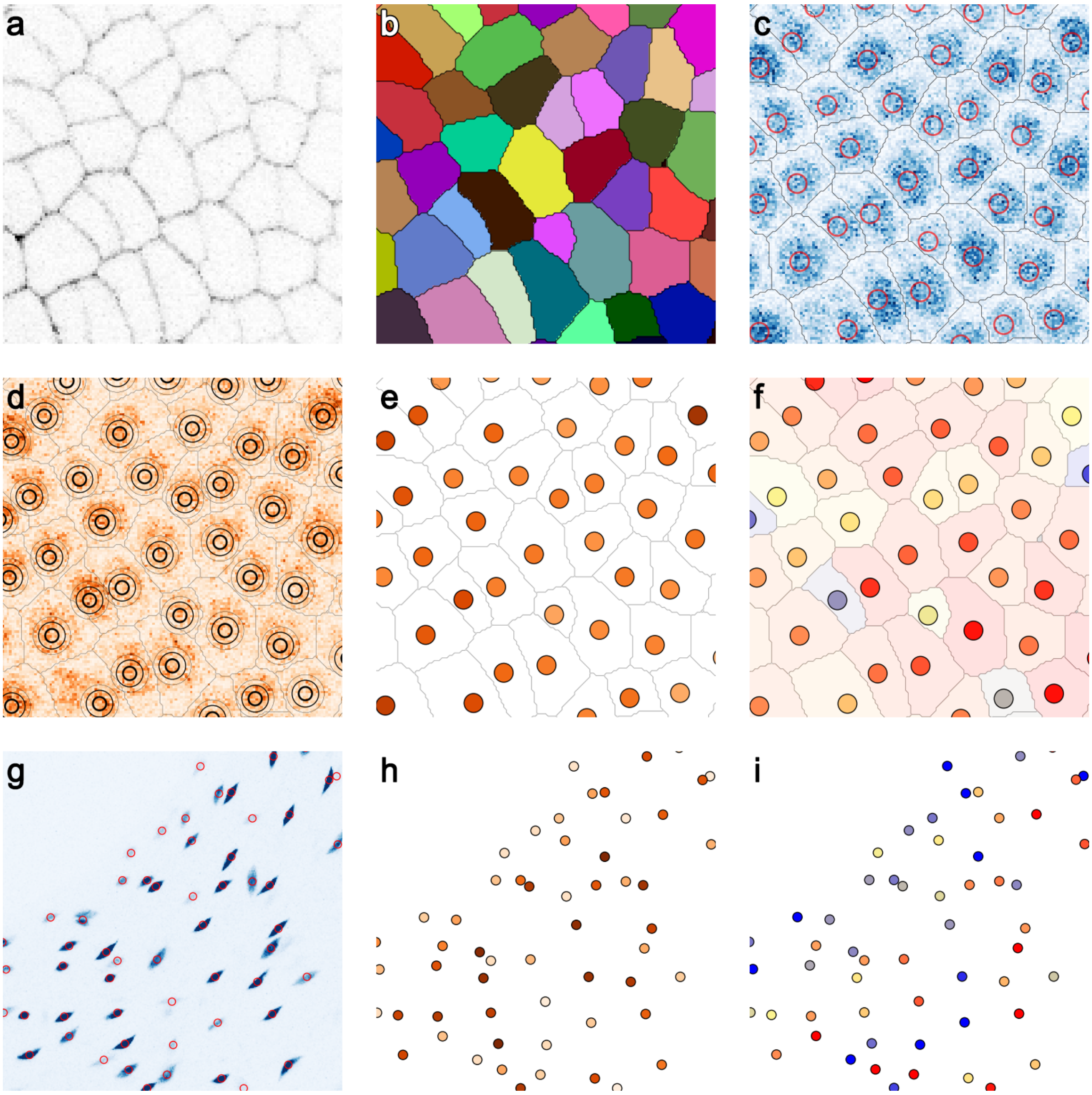


Supplementary Figure 2: **Automatic cell nuclei detection and ratiometric signal quantification.** **a-b**, The PI channel (**a**, slice view) provides a reference for the position of the cell walls in the 3D confocal stack, which is used to perform a watershed segmentation (**b**, surface view) that assigns each voxel of the image to a cell region, identified by a unique label. **c**, The Tag channel (**c**, slice view) is used to detect precisely the position of cell nuclei through a Gaussian scale-space transform, using the cell regions from the segmented image to limit the detection to one nucleus per cell. **d-e**, Signals such as the one provided by the RGA channel (**d**, slice view) are quantified as a Gaussian-weighted average of the intensity values around each detected nucleus, associating each cell with a signal value (**e**). To obtain the level of GA signalling, the Tag signal is quantified in the same way to be able to compute a ratio of the two signals. The quantified value for the GA signalling is obtained through the formula $GA = 3 - RGA/Tag$. **g-i**, In the case of hypocotyl images, the nuclei detection is performed on the Tag channel (**g**, max-intensity projection) with specific parameter values and without the cell region constraint, signals are then quantified similarly from their respective channel (**h**) and the GA ratiometric value computed using the same formula (**i**).

parameters this way, denoted $\kappa_c^-$, $\kappa_c^+$ and $\kappa_c$ for the minimum principal curvature, the maximal principal curvature and the mean curvature respectively, so that $\forall c \in \mathcal{L}_1$:

$$\kappa_c^- = \kappa^-(v(c)), \quad \kappa_c^+ = \kappa^+(v(c)), \quad \kappa_c = \kappa(v(c)) \tag{10}$$

#### 1.2 Cell-level Nuclei Detection & Signal Quantification

To quantify accurately the information provided by the GA biosensor, it is necessary to look at fluorescence intensity not at the global cell scale but more finely, at the level of the nucleus where the fluorescent proteins are targeted. To do so, we look within each cell region $\Gamma_c \mid c \in \mathcal{C} \setminus \{1\}$ for the precise position of the nucleus center point.

##### 1.2.1 Cell-level Nuclei Detection

To locate nuclei, we use the same 4D Gaussian scale-space transform [8] applied to the *pRPS5a:TagBFP* (Tag) channel, introduced in [1]. We obtain it by convolving the image $I_{\text{Tag}}$ with a sequence of 3D isotropic Gaussian kernels of increasing standard deviations $\sigma \in \mathfrak{s}$. The scale interval $\mathfrak{s}$ is defined as a geometric sequence varying from $\sigma_{\min}$ to $\sigma_{\max}$.

This results in a 4D response image, denoted by extension $I_{\text{Tag}} : \Omega_I \times \mathfrak{s} \subset \mathbb{R}^4 \to \mathcal{I}$, which measures how the signal intensity locally forms a spherical blob of scale $\sigma \in \mathfrak{s}$.

But instead of detecting local maxima in this 4D image, we use the information from the segmented image to detect only one nucleus per cell (Supplementary Figure 2 c). To do so, we retain for each cell labelled $c$, among all the voxels belonging to its region $\Gamma_c$, the position $P_c$ of the voxel that achieves the highest response independently of scale, so that $\forall c \in \mathcal{C} \setminus \{1\}$:

$$P_c = \underset{\mathbf{x} \in \Gamma_c}{\operatorname{argmax}} \Big( \max_{\sigma \in \mathfrak{s}} \left( I_{\text{Tag}}(\mathbf{x}, \sigma) \right) \Big). \tag{11}$$

We denote $\mathcal{P}$ the set of those nuclei points, identified by the integer label $c$ of the cell region to which they belong, so that we can write:

$$\mathcal{P} = \{ P_c = (x_c, y_c, z_c) \mid c \in \mathcal{C} \setminus \{1\} \} . \tag{12}$$

The parameter values we used are the ones identified in the evaluation study performed in [1], namely $|\mathfrak{s}| = 3$, $\sigma_{\min} = 0.8\mu m$ and $\sigma_{\max} = 1.4\mu m$.

##### 1.2.2 Nuclei Signal Quantification

We use the method of [1] to quantify the signal values at the level of the cell nuclei in each image channel separately. For the signal $S$, the signal value for the the cell labelled $c$ is computed as the value of the image $I_S$ filtered by a Gaussian kernel of radius $\sigma_{\mathcal{N}}$ at the voxel position $P_c$ (Supplementary Figure 2 d-e), so that $\forall c \in \mathcal{C} \setminus \{1\}$,

$$S_c = \left( I_S * G(\sigma_{\mathcal{N}}) \right) (P_c). \tag{13}$$

For example, the local level of expression of the $CLV3$ gene, imaged using *pCLV3:mCHERRY* in the channel $I_{\text{CLV3}}$ would we quantified as $\text{CLV3}_c = \left( I_{\text{CLV3}} * G(\sigma_{\mathcal{N}}) \right) (P_c)$.

The standard deviation we used for the quantification of nuclei signals in SAM images is $\sigma_{\mathcal{N}} = 0.75\mu m$.

##### 1.2.3 Nuclei Detection and Signal Quantification in Hypocotyl Images

In the specific case of hypocotyl images where no cell wall staining was performed, we rely on the nuclei detection method described in [1] to detect the nuclei in the $I_{\text{Tag}}$ (Supplementary Figure 2 g) and quantify both Tag and RGA signals (Supplementary Figure 2 h). Given that the cell sizes are different from those of shoot apical meristems, where the best parameter values were identified, we adjusted the method parameters to perform best on those images. The parameter values we retained are $|\mathfrak{s}| = 4$, $\sigma_{\min} = 2.5\mu m$, $\sigma_{\max} = 5\mu m$ and $\sigma_{\mathcal{N}} = 1\mu m$.

##### 1.2.4 Quantification of GA Signalling

In the case of the ratiometric GA sensor qRGA, we combine the information of two fluorescence channels $I_{\text{Tag}}$ and $I_{\text{RGA}}$ to compute the ratio of estimated signals for each nuclei point, so that $\forall c \in \mathcal{C} \setminus \{1\}$,

$$\text{qRGA}_c = \frac{\text{RGA}_c}{\text{Tag}_c}. \tag{14}$$

Finally, to represent the GA activity in the cell $c$, we estimate the quantity $\text{GA}_c$ referred to as *GA Signalling*. To cover the dynamic range of changes in the qRGA ratio, we define $\forall c \in \mathcal{C} \setminus \{1\}$,

$$\text{GA}_c = 3 - \text{qRGA}_c = 3 - \frac{\text{RGA}_c}{\text{Tag}_c}. \tag{15}$$

The formula is the same for both SAM images (Supplementary Figure 2 f) and hypocotyl images (Supplementary Figure 2 i). This quantity is the one displayed if Fig. 2fh, Fig. 3d, Fig. 4bd, and in Extended Data Fig. 3 and Extended Data Fig. 6.

#### 1.3 Image Rigid Temporal Registration

The previous steps were performed individually on each frame of the time-lapse acquisitions. In our study, we focused on sequences of observations of the same individual over its development, consisting of $K_t$ multichannel images $\{I(t_i) \mid i \in [\![0, K_t]\!]\}$, indexed by their temporal position $t_i \in \mathbb{N}$ in hours relatively to the first time of acquisition $t_0 = 0h$. In the remaining, we will consistently index data computed from the $i$-th acquisition $I(t_i)$ by the temporal index. For instance $\mathcal{P}(t_i) = \{P_c \mid c \in \mathcal{C}(t_i) \setminus \{1\}\}$ denotes the set of nuclei points detected in $I(t_i)_{\text{Tag}}$ using the cell regions defined by $I(t_i)_{\text{Seg}}$.

##### 1.3.1 Rigid Registration

Similarly to what was done in [1], we estimate 3D rigid transformations between consecutive time frames of a sequence using a block matching algorithm [9], in order to place the quantitative cell information of a given sequence in the same spatial reference frame. This estimation is performed on the *Propidium Iodide* channel $\{I(t_i)_{\text{PI}} \mid i \in [\![0, K_t]\!]\}$ of the consecutive images. This produces $K_t - 1$ isometry matrices in homogeneous coordinates $R_{t_i \leftarrow t_{i+1}}$ that can be inverted and/or multiplied to transform any frame of the sequence into the spatial reference frame of any other.

##### 1.3.2 Registered Nuclei Points

We use the registration output to transform all the detected nuclei points into the coordinate system of the first frame of the sequence. By applying the resulting rigid transforms to the the nuclei points detected at time $t_i$, we obtain a new point cloud $\mathcal{P}(t_i)^0$, indexed by the same set of integer cell labels $\mathcal{C}(t_i)$, such that $\forall c \in \mathcal{C}(t_i) \setminus \{1\}$,

$$P_c^0 = \left( \prod_{j=i-1}^{0} R_{t_j \leftarrow t_{j+1}} \right) P_c. \tag{16}$$

#### 1.4 Cell Surfacic Growth Computation

Cellular growth is generally expressed in terms of *strain*, which represents a relative variation of length [10]. On a 3D surface, the strain is locally two-dimensional and can be described by two *strain values* $\gamma^+ > \gamma^-$ associated to two orthogonal *principal directions of growth*. Each value characterizes the change of length of a 2D element along the corresponding direction, as a percentage of its initial length.

Assuming that the epidermis of a given L1 cell can be approximated by a planar surface, we consider that *cell surfacic growth* can be described by two strain values and two principal directions of growth, which lie within the cell surface plane. The surfacic strain between two consecutive time points $t_i$ and $t_{i+1}$ is computed by comparing the surface of a cell $c$ at time $t_i$ with the combined surface of all its descendant cells at time $t_{i+1}$.

##### 1.4.1 Manual L1 Cell Lineages

Cell lineages were manually generated by expertizing the segmented images $I(t_i)_{\text{seg}}$ and $I(t_{i+1})_{\text{seg}}$ obtained for two consecutive time points $t_i$ and $t_{i+1}$ (Supplementary Figure 3 a-b). The rigid transformation was applied to the images to make the visual comparison of consecutive time points easier. The cell lineage itself $L(t_i, t_{i+1})$ consists of a set of couples of L1 cell labels $(c, c')$ where:

- $c' \in \mathcal{L}_1(t_{i+1})$ is the label of a cell in the segmented image $I(t_{i+1})_{\text{seg}}$.
- $c \in \mathcal{L}_1(t_i)$ is the label of the cell in the segmented image $I(t_i)_{\text{seg}}$ that will give rise to the cell labelled $c'$ at $t_{i+1}$.

We say that the cell $c$ is the *mother cell* of the cell $c'$, and conversely that the cell $c'$ is a *daughter cell* of the cell $c$.

##### 1.4.2 Cell Landmark Extraction

We choose to use the topological elements representing junctions between 2 or more cells in the segmented image as landmarks to compute the deformation of the tissue, as it is done in the literature [11]. However, to avoid relying only on the position of cell vertices, which can be very sensitive to segmentation artifacts, we use additional landmarks representing cell edges and cell interfaces, as proposed in [12] (Chapter 3.2.2: Quantifying cellular features).

We compute the landmark positions using the 3D positions of image topological elements (surfels, linels and pointels) at the interface between cells, and assign them a unique identifier consisting in a cell label tuple (Supplementary Figure 3 d). More precisely to extract landmarks on a segmented image $I_{\text{seg}}$, we extract for each cell $c \in \mathcal{L}_1$:

- A landmark identified by $(c)$ computed as the geometric median of $\Gamma^2_{1,c}$
- $\forall c' \in N(c) \cap \mathcal{L}_1$, a landmark identified by $(c, c')$ computed as the geometric median of $\Gamma^1_{1,c,c'}$
- $\forall c', c'' \in N(c) \cap \mathcal{L}_1$, such that $c'' \in N(c')$, a landmark identified by $(c, c', c'')$ computed as the geometric median of $\Gamma^0_{1,c,c',c''}$

Combining the segmented image $I(t_{i+1})_{\text{seg}}$ with the lineage information makes it possible to obtain a relabelled image where all the daughter cells of a cell $c$ from $I(t_i)_{\text{seg}}$ carry the label $c$ (Supplementary Figure 3 b-c). We perform the landmark extraction process on both the *mother* segmented image $I(t_i)_{\text{seg}}$ (Supplementary Figure 3 d) and the relabelled *daughter* image (Supplementary Figure 3 e) to obtain two sets of landmark points that can then be paired using their cell label identifiers.

##### 1.4.3 Surfacic Strain Estimation

For each cell $c \in \mathcal{L}_1(t_i)$ that has at least one daughter in $\mathcal{L}_1(t_{i+1})$, we are able to identify pairs of conserved cell landmark points between the mother and daughter segmented images. Each point is first projected on the 2D plane approximating the interface between the cell $c$ and the background in its respective image. It is then centered by subtracting the projected position of the landmark identified by $(c)$, representing the median point of the considered interface.

Based on the paired centered 2D positions (provided there are at least 4 of them), we compute a 2D transformation (rotation and scaling) by a linear regression without intercept, minimizing squared

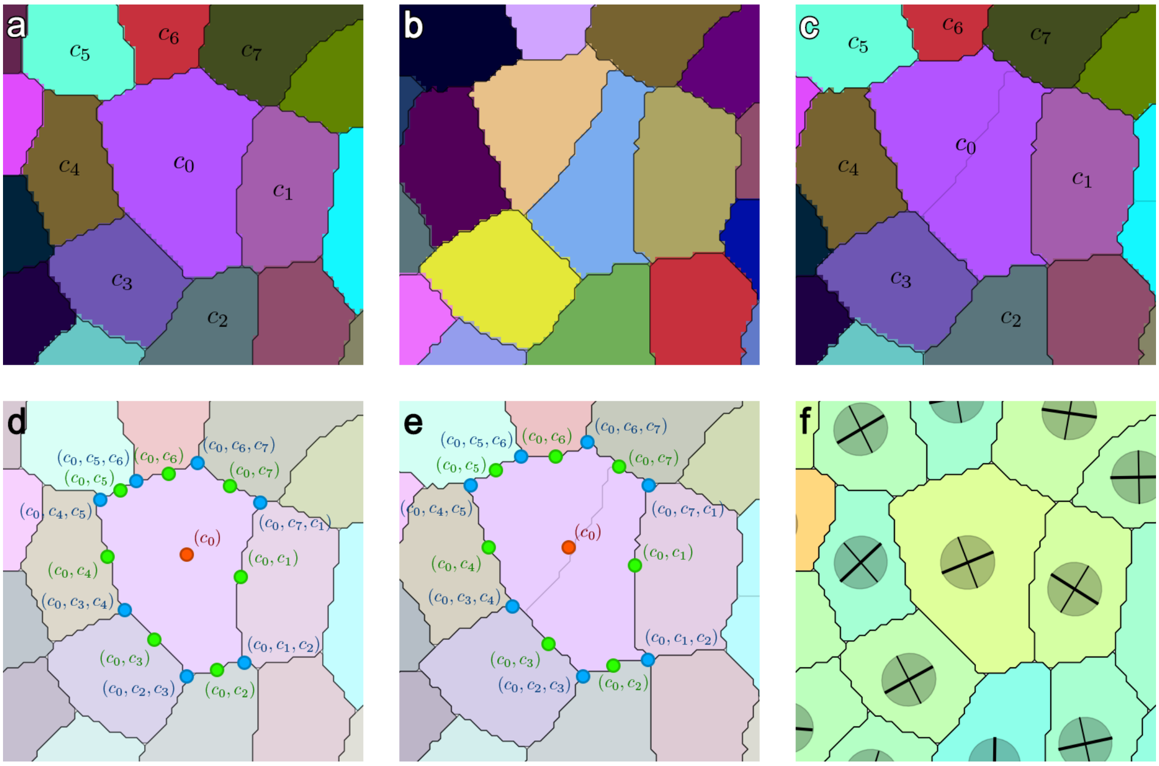


Supplementary Figure 3: **Surfacic strain estimation using cell landmarks. a-b**, View of the surface of segmented images of the same tissue at two consecutive time points. Each cell carries a label identifier that is used to manually define lineages of L1 cells. **c**, The lineage information allows to relabel the image of the second time point using the cell labels of the first one. **d-e**, Landmarks representing the junctions between two or more labels in the image are extracted on both the first image (**d**) and the relabelled second image (**e**). The tuples of cell labels associated with each landmark point allow to pair them between the two consecutive time points. **f**, In each cell, the linear least-squares estimation of the deformation results in the determination of a surfacic strain tensor, defining two strain values and two orthogonal growth directions.

distance between pairs of landmarks [12, 11]. The singular value decomposition of the obtained 2D matrix allows to obtain the two strain values $\gamma_c^+$ and $\gamma_c^-$ and their associated directions in the 2D cell plane. We reconstruct a 3D strain matrix in the reference frame of $I(t_{i+1})_{\text{seg}}$ by transforming the 2D matrix back into the projection base, assuming a strain value of 0 in the normal direction to the cell plane. This produces a "flat" tensor that represents the surfacic strain in 3D (Supplementary Figure 3 f).

The surfacic strain estimation gives us several important growth features. We derive the *surfacic growth intensity* $\gamma_c$ as the product of the strain values, so that $\forall c \in \mathcal{L}_1(t_i)$,

$$\gamma_c = \gamma_c^+ \cdot \gamma_c^- . \tag{17}$$

We also define the *surfacic growth anisotropy* $a_c$ that measures how much the cell grows in a preferential direction. A growth anisotropy equal to 0 reflects the fact that growth is homogeneous in every direction, hence we define anisotropy as the norm of the deviatoric part of the strain tensor, normalized by the norm of its isotropic part. In the case of the 2D surfacic strain tensor, it is equivalent to define this measure as $\forall c \in \mathcal{L}_1(t_i)$,

$$a_c = \frac{\gamma_c^+ - \gamma_c^-}{\gamma_c^+ + \gamma_c^-} . \tag{18}$$

#### 1.5 Curvature-based SAM Primordia Alignment

In order to pool cell data coming from several individuals SAMs, we perform a population alignment, similar to what was done in [1]. The general idea is to find a geometrical transformation that places organs with a similar state of development at the same location in the 3D space. This allows then to aggregate the data initially expressed in each individual's own image reference frame into a common reference frame where pointwise comparison is meaningful.

##### 1.5.1 SAM Reference Frame

The target reference frame into which we aim to transform the data is the one introduced in [1]. It consists in a cylindrical coordinate system $(r, \theta, z)$ in which:

- the origin corresponds to the apex center in the central zone (CZ) of the meristematic dome
- the $z$-axis corresponds to the main rotational symmetry axis of the meristematic dome
- the *polar* axis ($\theta = 0$) corresponds to the direction of the last initiated primordium ($P_0$)
- the rotation orientation corresponds to the orientation (either clockwise or counter-clockwise) of the phyllotactic spiral.

Such a coordinate system can actually be mapped on the data by identifying the spatial positions of its key landmarks (CZ center $\mathbf{c} = (x_\mathbf{c}, y_\mathbf{c}, z_\mathbf{c}) \in \mathbb{R}^3$, unitary vertical axis $\mathbf{a} \in \mathbb{R}^3$, unitary $P_0$ radial vector $\mathbf{r} \in \mathbb{R}^3 \mid \mathbf{a} \perp \mathbf{r}$, and binary orientation $o \in \{-1, 1\}$). The computation of the alignment rigid geometric transformation from this set of landmarks is then direct.

##### 1.5.2 2D Maps of L1 Signal

To detect the position of the landmarks in the cell-level data of a given individual SAM, we rely on a tool introduced in [1] that allows to infer a value of a signal $S$ in any point of a 2D projection space, using the signal values of the subset $\mathcal{L}_1$ of first-layer cells. This computation relies on a 1D parametric sigmoid density function:

$$\begin{aligned} \eta \ : \ & \mathbb{R} \to [0, 1] \\ & r \mapsto \frac{1}{2} - \frac{1}{2} \tanh\big(k \cdot (r - R)\big) \end{aligned} \tag{19}$$

for which we use the optimal parameter values evidenced in [1]: $R = 7.5\mu m$ and $k = 0.55\mu m^{-1}$. This allows to estimate a continuous signal map $\widehat{S}$ at any point $\mathbf{x} \in \mathbb{R}^2$ based on the projected point cloud of L1 cell points $\big\{P_c \in \mathbb{R}^2 \mid c \in \mathcal{L}_1\big\}$ and the associated signal values $\{S_c \mid c \in \mathcal{L}_1\}$ as: $\forall \mathbf{x} \in \mathbb{R}^2$,

$$\widehat{S}(\mathbf{x}) = \frac{1}{\sum\limits_{c \in \mathcal{L}_1} \eta\left(\|\mathbf{x} - P_c\|\right)} \sum_{c \in \mathcal{L}_1} \eta\left(\|\mathbf{x} - P_c\|\right) S_c. \tag{20}$$

We consider that the signal map $\widehat{S}(\mathbf{x})$ is defined only for points $\mathbf{x}$ where the total density $H(\mathbf{x}) = \sum_{c \in \mathcal{L}_1} \eta\left(\|\mathbf{x} - P_c\|\right)$ is greater than $\frac{1}{2}$.

##### 1.5.3 SAM Center Detection

The *pCLV3:mCHERRY* image channel (Supplementary Figure 4 a) provides a marker of the central zone (CZ) of the meristem. We use the same method as in [1] to estimate the 3D position of the CZ center $\mathbf{c} = (x_\mathbf{c}, y_\mathbf{c}, z_\mathbf{c})$, using the quantified signal values $\{\mathrm{CLV3}_n \mid n \in \mathcal{L}_1\}$.

The procedure relies on the computation of a 2D map $\widehat{\mathrm{CLV3}}$ based on the registered nuclei points $\mathcal{P}^0$ from all the time points of the sequence (Supplementary Figure 4 b). This map allows to estimate a 2D position $(x_\mathbf{c}, y_\mathbf{c})$ of the CZ center, at which we compute the value of the altitude map $\widehat{z}$ to obtain the third coordinate of the center point $z_\mathbf{c}$.

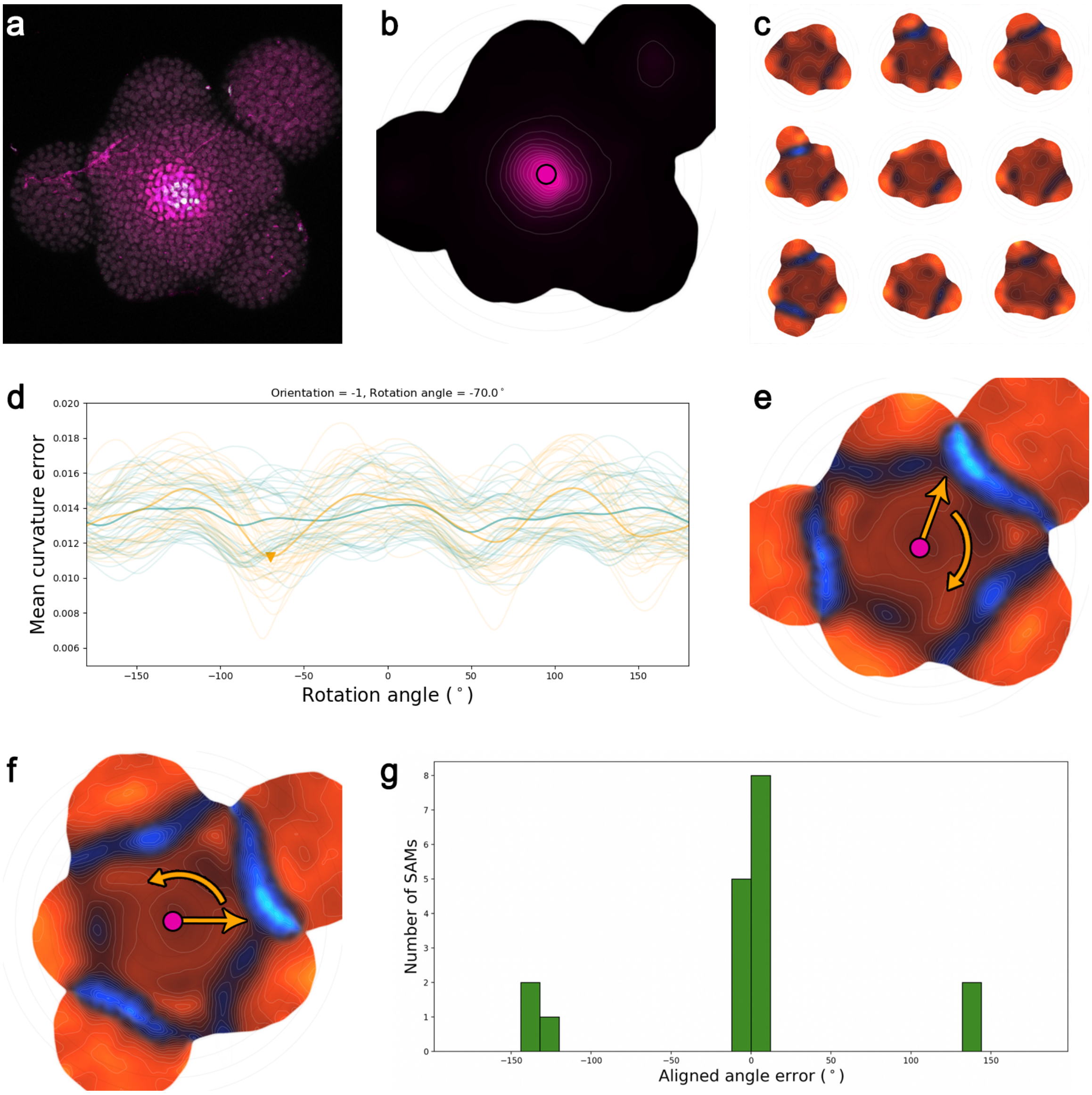


Supplementary Figure 4: **SAM alignment onto a reference set of curvature maps. a-b**, The CLV3 image channel (**a**, max-intensity projection), quantified at the level of each cell nucleus, is used to locate precisely the center of the SAM. It is done using a continuous 2D map (**b**), which is computed using the registered positions of all L1 nuclei of the sequence. This allows to center the nuclei point cloud, and subsequently to correct its tilting by estimating the main vertical axis of the SAM. **c-e**, Next, to identify the direction of the lastly initiated primordium, we rely on a database of aligned SAM point clouds from [1] on which curvature has been computed. This provides a set of reference 2D curvature maps (**c**, subset of the database) that are used to determine an optimal angle and orientation (**d**, error curves obtained for each reference map, $o = 1$ in green and $o = -1$ in orange) by minimizing the error with the 2D curvature map of the considered SAM (**e**). **f**, This process allows to align the SAM in a common reference frame where inter-individual comparison is possible. **g**, The evaluation of this alignment procedure on the reference database shows that orientation is correctly predicted and that the correct alignment angle is retrieved in 72% of the cases. All the remaining cases show an error corresponding to a shift of ś1 divergence angle between two consecutive organs.

##### 1.5.4 Vertical Axis Optimization

Using the method from [1], we estimate the rotational symmetry axis $\mathbf{a}$ by looking for the direction that minimizes the dispersion of $z$ coordinates of the CZ nuclei as a function of their radial distance.

##### 1.5.5 Curvature-based Alignment

The two previous landmarks $\mathbf{c}$ and $\mathbf{a}$ allow to define a reference frame equipped with a cylindrical coordinate system $(r, \theta, z)$, in which the positioning of the direction $\mathbf{r}$ of the $P_0$ organ primordium comes down to the determination of an angular coordinate $\theta_0$.

In [1], this was achieved by detecting the maximal point of the 2D map of Auxin signal in the peripheral zone (PZ) of the meristem. In our case however, the Auxin information is not available. Still, it is possible to indirectly use this information by using the SAM data aligned with the method from [1] as a geometrical reference for the alignment of new meristems.

We rely on the mean curvature information $\kappa$ to represent accurately the geometrical information of the SAM surface in a 2D representation like the signal maps. Practically, we selected the $K_s = 21$ sequences of SAM acquisitions from [1] from which we use the aligned positions of L1 nuclei to compute $K_s$ 2D aligned maps of mean curvature, noted $\left\{\widehat{\kappa_j^*} \mid j \in [\![0, K_s[\![\right\}$ (Supplementary Figure 4 c).

We use this set of reference maps to find optimal values for both the angular coordinate $\theta_0$ of the $P_0$ primordium and the clockwise or counter-clockwise orientation $o$ of the organ spiral by minimizing the error between the mean curvature map $\widehat{\kappa}$ obtained using nuclei points from all the time points of the considered sequence, rotated by the angle $\theta_0$ and reflected if $o = -1$, and all the reference mean curvature maps $\widehat{\kappa_j^*}$.

The error between $\widehat{\kappa}$ and $\widehat{\kappa_j^*}$ is computed as the average of the pointwise absolute error over the intersection of the definition domains of both maps, restricted to the central and peripheral zones (PZ) of the SAM. We define the limits of the PZ by a radial distance threshold $r_{\mathrm{PZ}} = 70\mu m$. This allows to delimit a 2D domain that actually depends of the considered rotation angle and orientation value:

$$\mathrm{PZ}_j(\theta_0, o) = \left\{(r, \theta) \in \mathbb{R}^2 \mid H(r, o \cdot (\theta - \theta_0)) > \frac{1}{2}, H_j^*(r, \theta) > \frac{1}{2}, r < r_{\mathrm{PZ}}\right\}. \quad (21)$$

The optimal values $\theta_0^*$ and $o^*$ (Supplementary Figure 4 d) can then be written as:

$$(\theta_0^*, o^*) = \underset{\substack{\theta_0 \in [-\pi, \pi[ \\ o \in \{-1, 1\}}}{\arg\min} \left( \frac{1}{K_s} \sum_{j=0}^{K_s - 1} \frac{1}{\iint_{\mathrm{PZ}_j(\theta_0, o)} r \mathrm{d}r \mathrm{d}\theta} \iint_{\mathrm{PZ}_j(\theta_0, o)} \left| \widehat{\kappa_j^*}(r, \theta) - \widehat{\kappa}\big(r, o \cdot (\theta - \theta_0)\big) \right| r \mathrm{d}r \mathrm{d}\theta \right). \quad (22)$$

The determination of $\theta_0$ allows to position the radial vector $\mathbf{r}$, and the value of $o$ achieves the definition of the cylindrical coordinate system (Supplementary Figure 4 e)

##### 1.5.6 Evaluation of curvature-based alignment

To evaluate this new alignment procedure, we applied it the 21 aligned SAM sequences we use as reference. We used a leave-one-out strategy, aligning the considered SAM by minimizing the average curvature error with the 20 remaining series. In the ideal case, the optimal orientation $o$ should be equal to 1 and the optimal $P_0$ angular coordinate value $\theta_0$ should be equal to 0°.

Our evaluation shows that the optimal orientation is always 1, and in the vast majority of the cases (72%) the optimal angle is such that $|\theta_0| \leq 6°$ (Supplementary Figure 4 g). In the rest of cases, the optimal angle corresponds to one "golden" divergence angle of the organ phyllotactic spiral $\alpha^* \sim 137.5°$, as either $|\theta_0 + \alpha^*| \leq 6°$ (17%) or $|\theta_0 - \alpha^*| \leq 6°$ (11%).

In other terms the alignment procedure either finds the right angle, or performs a shift of one primordium (aligning for instance $P_1$ on $P_0$. In any case, it manages to successfully superimpose organ primordia, but with a possible uncertainty on the equivalence of their developmental state.

This temporal uncertainty is also explained by the fact that the reference SAMs themselves are not perfectly homogeneous in terms of developmental state. The fact that most alignement angles are correct, and that both negative and positive primordium shifts exist in the dataset confirms that the reference SAMs are "centered" on what can be seen as a sound characterization of the $P_0$ stage, and validates the approach of using them as a geometrical reference.

###### 1.5.7 Aligned L1 nuclei points

In the end, the determination of the SAM landmarks **c**, **a**, **r** and $o$ allows to transform the sequence registered points $\mathcal{P}^0$ into the common 3D reference frame in which we will be able to compare different individuals locally, as described in [1].

We note $\mathcal{P}^* = \{P_c^* \mid c \in \mathcal{L}_1\}$ the positions of first-layer nuclei points in this common reference frame. The 2D maps computed using these aligned positions (Supplementary Figure 4 f) make it possible to perform pointwise inter-individual comparisons, as the data points that are superimposed at a given location correspond to tissue areas in a similar state of development. Therefore, it opens the way to population statistics.

These nuclei positions are the ones displayed in Fig. 4c, and were used to compute the average GA signalling map displayed in Fig. 4b and the average surfacic growth intensity and surfacic growth anisotropy maps displayed in Fig. 5a-b.

[11] Pierre Barbier de Reuille, Anne-Lise Routier-Kierzkowska, Daniel Kierzkowski, George W Bassel, Thierry Schüpbach, Gerardo Tauriello, Namrata Bajpai, Sören Strauss, Alain Weber, Annamaria Kiss, Agata Burian, Hugo Hofhuis, Aleksandra Sapala, Marcin Lipowczan, Maria B Heimlicher, Sarah Robinson, Emmanuelle M Bayer, Konrad Basler, Petros Koumoutsakos, Adrienne HK Roeder, Tinri Aegerter-Wilmsen, Naomi Nakayama, Miltos Tsiantis, Angela Hay, Dorota Kwiatkowska, Ioannis Xenarios, Cris Kuhlemeier, and Richard S Smith. Morphographx: A platform for quantifying morphogenesis in 4d. *eLife*, 4:e05864, may 2015.

[12] Jonathan Legrand. *Toward a multi-scale understanding of flower development - from auxin networks to dynamic cellular patterns.* Theses, Ecole normale supérieure de lyon - ENS LYON, November 2014.
