## Supplementary Method 2 for "A quantitative gibberellin signalling biosensor reveals a role for gibberellins in internode specification at the shoot apical meristem"

### 2 Synthesis and characterization of GA-Fluorescein (GA-Fl)

#### 2.1 General chemistry methods and instrumentation

$GA_3$, $GA_4$ and $GA_7$ were purchased from Duchefa Biochemie. All other chemicals were purchased from Merck or Combi-Blocks and were used as received unless otherwise stated. Anhydrous solvents and reagents were obtained as SureSeal bottles from Merck. Thin-layerchromatography and flash chromatography were performed using Merck KGaA pre-coated silica gel 60 F-254 plates and Silicycle silica gel 40-63 (230-400 mesh), respectively. UV absorbance spectra were recorded on Agilent Cary 60 UV-Vis Spectrophotometer. Fluorescence spectra were recorded on Fluorolog 2 (Spex) fluorimeter. Low resolution ESI mass spectrometry was performed on LC/MS Acquity QDa detector coupled with Waters HPLC. High resolution ESI mass spectrometry was performed on a Waters SYNAPT system. $^{1}H$ and $^{13}C$ NMR spectra were collected in DMSO-d6 (Cambridge Isotope Laboratories, Cambridge, MA) at 25oC using a Bruker Advance III spectrometer at 400 MHz and 100 MHz respectively at the Department of Chemistry NMR Facility at Tel-Aviv University. All chemical shifts are reported in the standard δ notation of parts per million using either TMS or residual solvent peak as an internal reference. Abbreviations: MeCN: acetonitrile, DMF: dimethylformamide, HATU: Hexafluorophosphate Azabenzotriazole Tetramethyl Uronium, TFA: Trifluoroacetic acid, DIPEA: *N*,*N*-Diisopropylethylamine.

#### 2.2 HPLC-MS Analysis conditions

HPLC-MS analysis was performed on Waters HPLC with XBridge C18 column (100 X 3 mm, 5 μm), starting with 2 min of solvent A (water), followed by a water-acetonitrile gradient from 0% to 100% solvent B (acetonitrile) in 15 minutes then 1 minute at 100% solvent B and ending with 2 min 100% solvent A at flow rate of 1 mL/min (solvent A = water, solvent B = acetonitrile, both contain 0.1% TFA as an additive). Mass spectrometry was performed on LC/MS Acquity QDa detector coupled with Waters HPLC.

#### 2.3 Preparative HPLC purification conditions

Preparative HPLC was performed on Waters 2545 HPLC with XBridge C18 column (100 X 19 mm, 5 μm) starting with 2 min of solvent A (water), followed by a water-acetonitrile gradient from 0% to 80% solvent B (acetonitrile) in 20 minutes, continuing with 3 min gradient of 80% to 100% solvent B then 3 minutes at 100% solvent B and ending with 2 min 100% solvent A at flow rate of 15 mL/min (solvent A = water, solvent B = acetonitrile, both contain 0.1% TFA as an additive).

#### 2.4 Synthesis of GA-Fluorescein (GA-Fl) derivatives

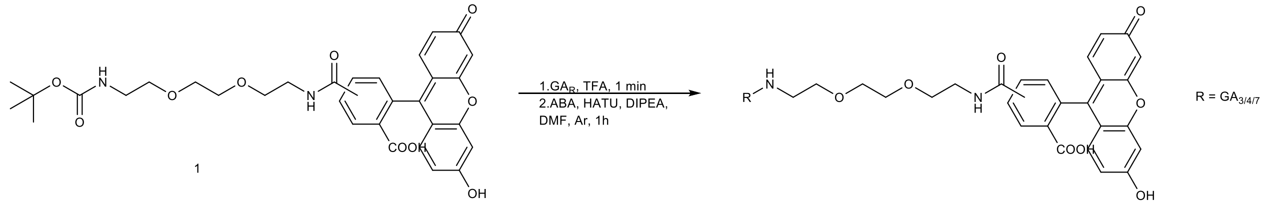


GA-Fluorescein **(GA-Fl)** were synthesized using a previously described protocol [1]. Briefly, compound **1** (10 mg, 1 eq) was dissolved in 0.5 mL TFA and stirred for 1 minute. TFA was immediately removed under reduced pressure. The residue was dissolved in dry DMF under argon atmosphere and DIPEA (3.3 µL, 1.1 eq) was added. In a separate 1.5 mL eppendorf, the appropriate GA (3, 4 or 7, 1.1 eq) was dissolved in dry DMF, then DIPEA (7.6 µL, 1.1 eq) and HATU (7.2 mg, 1.1 eq) were added. The mixture was vortexed for 2 minute and was then added to the TFA-treated compound **1**. The reaction was stirred at room temperature for 1 hour and the solvent was removed under reduced pressure. The residue was dissolved in 2 mL acetonitrile and the desired product was purified using preparative HPLC (see preparative HPLC purification conditions above).

$GA_3$-Fl: Preparative HPLC retention time: 13.43 min. Obtained 10 mg yellow solid (12 µmol, yield 68%). $^1$H NMR (400 MHz, DMSO) δ 10.16 (s, 1H), 8.89 (d, *J* = 5.5 Hz, 1H), 8.74 (s, 1H), 8.44 (d, *J* = 1.6 Hz, 1H), 8.22 (dd, *J* = 8.1, 1.6 Hz, 1H), 8.17 – 8.01 (m, 2H), 7.35 (d, *J* = 8.0 Hz, 1H), 6.67 (d, *J* = 2.3 Hz, 1H), 6.59 – 6.49 (m, 3H), 6.30 (dd, *J* = 9.3, 3.2 Hz, 1H), 5.76 (td, *J* = 9.4, 3.6 Hz, 1H), 5.56 (d, *J* = 6.6 Hz, 1H), 5.48 (dd, *J* = 6.7, 2.0 Hz, 1H), 5.10 (s, 1H), 5.03 (s, 1H), 4.82 (s, 1H), 4.76 (d, *J* = 4.7 Hz, 1H), 3.83 (ddd, *J* = 13.2, 7.0, 3.9 Hz, 2H), 3.57 – 3.39 (m, 9H), 3.11 – 3.02 (m, 2H), 2.55 (s, 1H), 2.14 (d, *J* = 8.0 Hz, 1H), 2.05 (s, 1H), 1.90 – 1.51 (m, 9H), 1.21 (s, 1H), 1.07 – 1.00 (m, 3H). $^{13}$C NMR (101 MHz, DMSO) δ 171.29, 160.09, 152.29, 133.76, 131.98, 129.58, 124.69, 113.14, 106.63, 105.94, 102.74, 91.36, 77.24, 77.05, 69.99, 69.53, 53.57, 50.91, 50.19, 49.86, 44.77, 17.00, 14.95, 14.67.; LC/MS: Retention time 9.15 min, 835.33 $[M+H]^+$. HR-MS(ESI) calcd. for formula $C_{46}H_{47}N_2O_{13}$ $[M+H]^+$: 835.3076; found: 835.3078.

$GA_4$-Fl: Preparative HPLC retention time: 16.23 min. Obtained 10 mg yellow solid (12 µmol, yield 70%). $^1$H NMR (400 MHz, DMSO) δ 9.98 (s, 1H), 8.71 (t, *J* = 5.5 Hz, 1H), 8.29 (d, *J* = 1.6 Hz, 1H), 8.07 (dd, *J* = 8.0, 1.6 Hz, 1H), 7.93 – 7.79 (m, 1H), 7.20 (d, *J* = 8.1 Hz, 1H), 6.52 (t, *J* = 2.6 Hz, 1H), 6.39 (dtq, *J* = 8.7, 4.5, 2.0 Hz, 2H), 4.70 (d, *J* = 8.3 Hz, 1H), 4.60 (d, *J* = 7.2 Hz, 1H), 3.37 – 3.26 (m, 7H), 3.03 – 2.87 (m, 7H), 2.36 (d, *J* = 1.9 Hz, 1H), 2.31 (d, *J* = 1.9 Hz, 1H), 1.84 (dd, *J* = 12.5, 5.4 Hz, 3H), 1.72 – 1.62 (m, 2H), 1.59 – 1.48 (m, 3H), 1.41 – 1.25 (m, 6H), 1.12 – 1.04 (m, 7H), 0.81 (t, *J* = 6.2 Hz, 2H), 0.68 (t, *J* = 6.5 Hz, 1H).$^{13}$C NMR (101 MHz, DMSO) δ 160.06, 157.78, 152.29, 129.59, 113.13, 109.56, 106.91, 102.73, 94.30, 69.98, 69.52, 69.24, 69.15, 54.76, 53.37, 52.56, 51.18, 50.99, 44.84, 37.35, 33.79, 31.54, 29.46, 27.29, 24.91, 22.53, 16.12, 14.87.; LC/MS: Retention time 10.23 min, 821.27 $[M+H]^+$. HR-MS(ESI) calcd. for formula $C_{46}H_{47}N_2O_{12}$ $[M-H]^+$: 819.3129; found: 819.3123.

$GA_7$-Fl: Preparative HPLC retention time: 16.44 min. Obtained 9 mg yellow solid (11 µmol, yield 63%). $^1$H NMR (400 MHz, DMSO) δ 10.12 (s, 1H), 8.72 (t, *J* = 5.5 Hz, 1H), 8.15 (dd, *J* = 8.1, 1.4 Hz, 1H), 8.06 (d, *J* = 8.1 Hz, 1H), 8.01 (d, *J* = 6.0 Hz, 1H), 7.66 (d, *J* = 1.1 Hz, 1H), 6.67 (t, *J* = 2.6 Hz, 1H), 6.60 – 6.50 (m, 3H), 6.31 (d, *J* = 9.3 Hz, 1H), 5.75 (dd, *J* = 9.3, 3.6 Hz, 1H), 4.86 (s, 1H), 4.75 (s, 1H), 3.82 (s, 1H), 3.56 – 3.38 (m, 9H), 3.21 – 2.99 (Im, 10H), 2.56 – 2.52 (m, 1H), 2.45 (s, 1H), 1.98 (s, 2H), 1.90 – 1.73 (m, 3H), 1.51 (tt, *J* = 17.4, 9.1 Hz, 4H), 1.31 – 1.19 (m, 2H), 1.03 (d, *J* = 6.5 Hz, 2H), 0.95 (d, *J* = 6.5 Hz, 1H).$^{13}$C NMR (101 MHz, DMSO) δ 179.60, 178.98, 171.19, 171.17, 168.49, 165.06, 160.09, 158.49, 158.18, 157.74, 157.70, 153.17, 152.30, 141.02, 136.63, 135.11, 133.67, 132.17, 129.85, 129.66, 128.66, 125.31, 122.74, 113.20, 109.60, 107.09, 102.72, 91.63, 83.73, 69.98, 69.88, 69.52, 69.45, 69.23, 69.09, 68.99, 53.56, 52.11, 51.95, 51.52, 44.99, 38.94, 36.93, 31.49, 15.87, 14.71.; LC/MS: Retention time 10.30 min, 819.30 $[M+H]^+$. HR-MS(ESI) calcd. for formula $C_{46}H_{45}N_2O_{12}$ $[M-H]^+$: 817.2973; found: 817.2961.

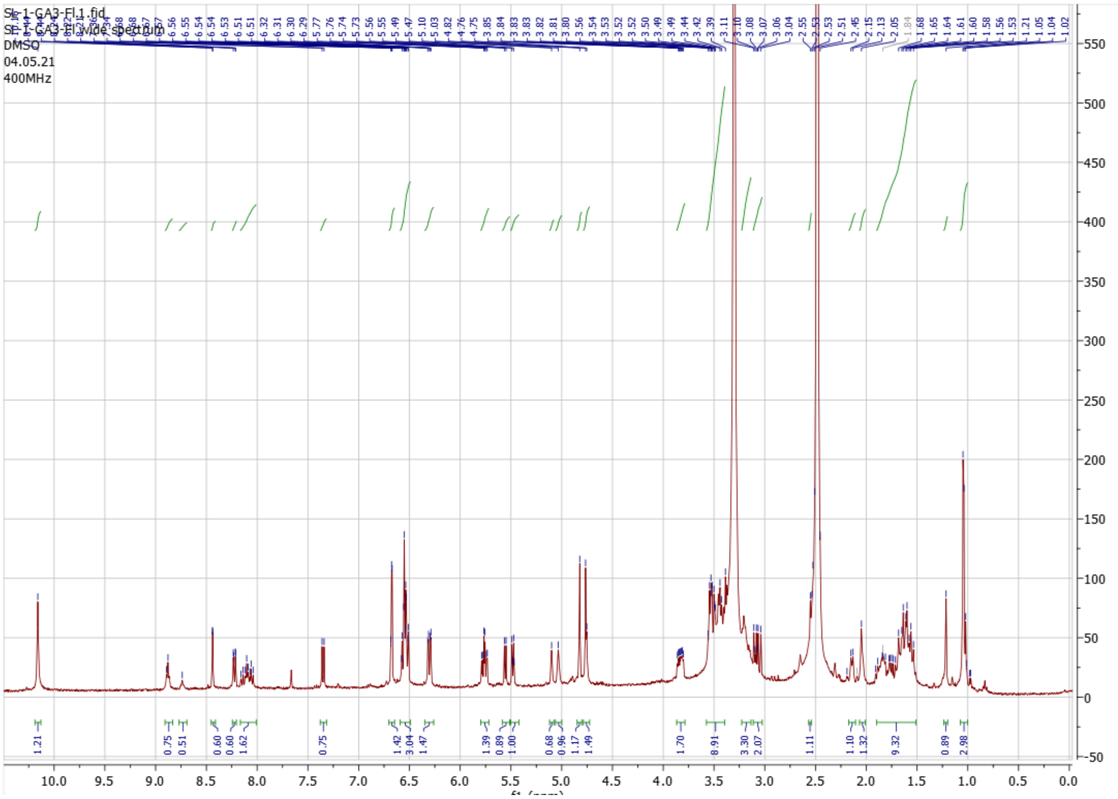


Supplementary Figure 5. $^{1}$H-NMR spectrum of $GA_3$-Fl.

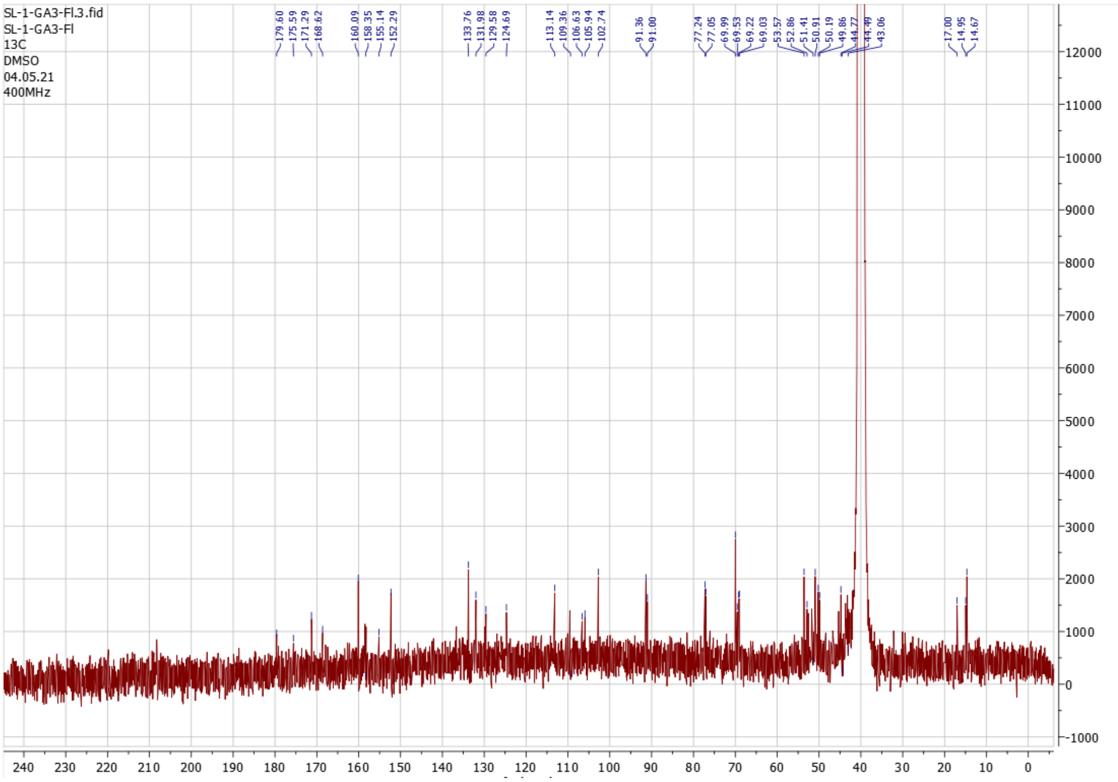


Supplementary Figure 6. $^{13}$C-NMR spectrum of $GA_3$-Fl

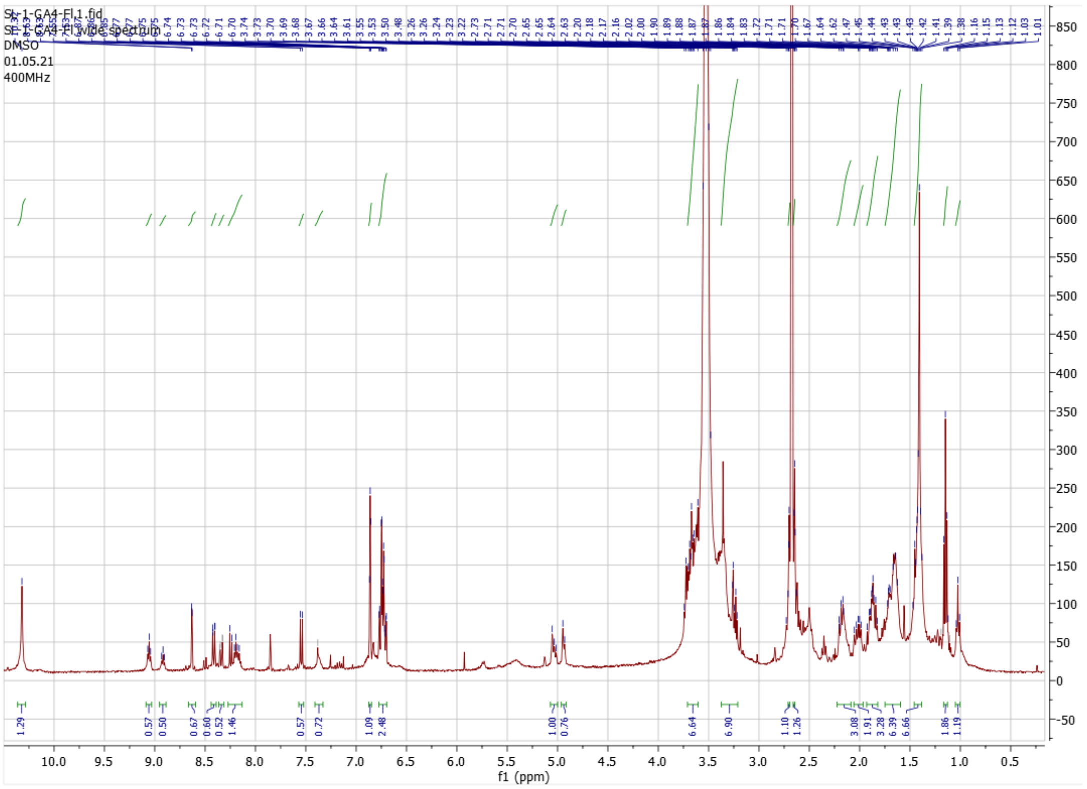


Supplementary Figure 7. $^{1}$H-NMR spectrum of $GA_4$-Fl

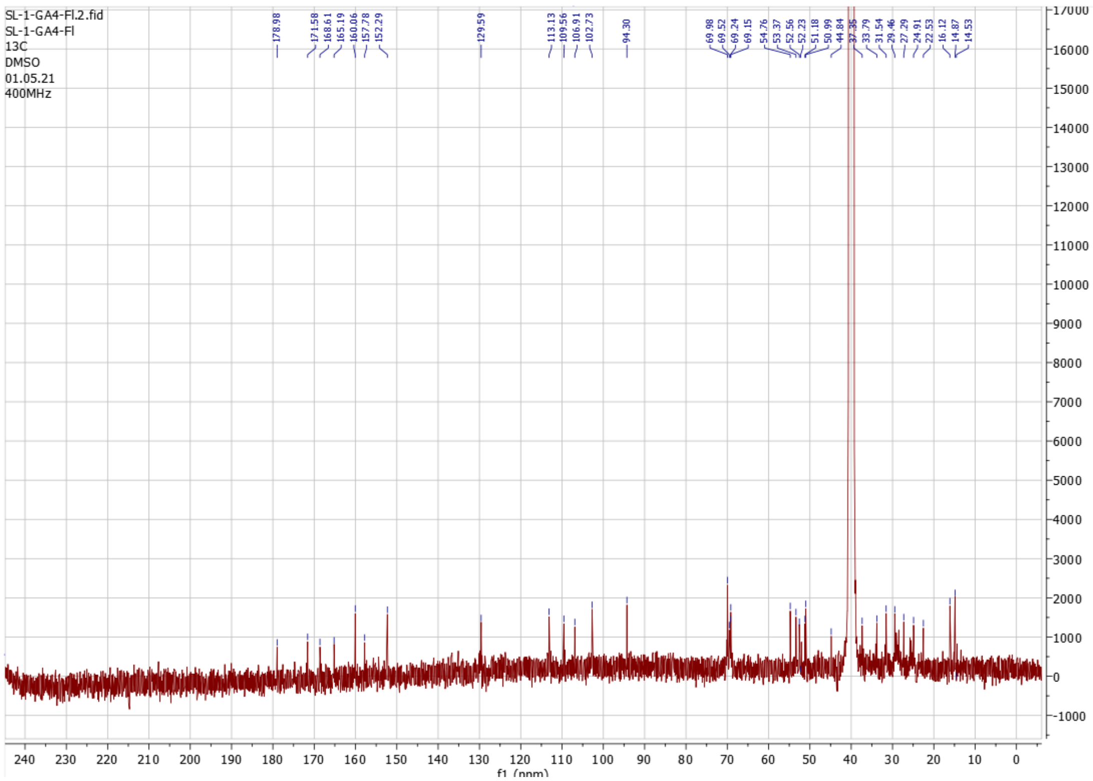


Supplementary Figure 8. $^{13}$C-NMR spectrum of $GA_4$-Fl

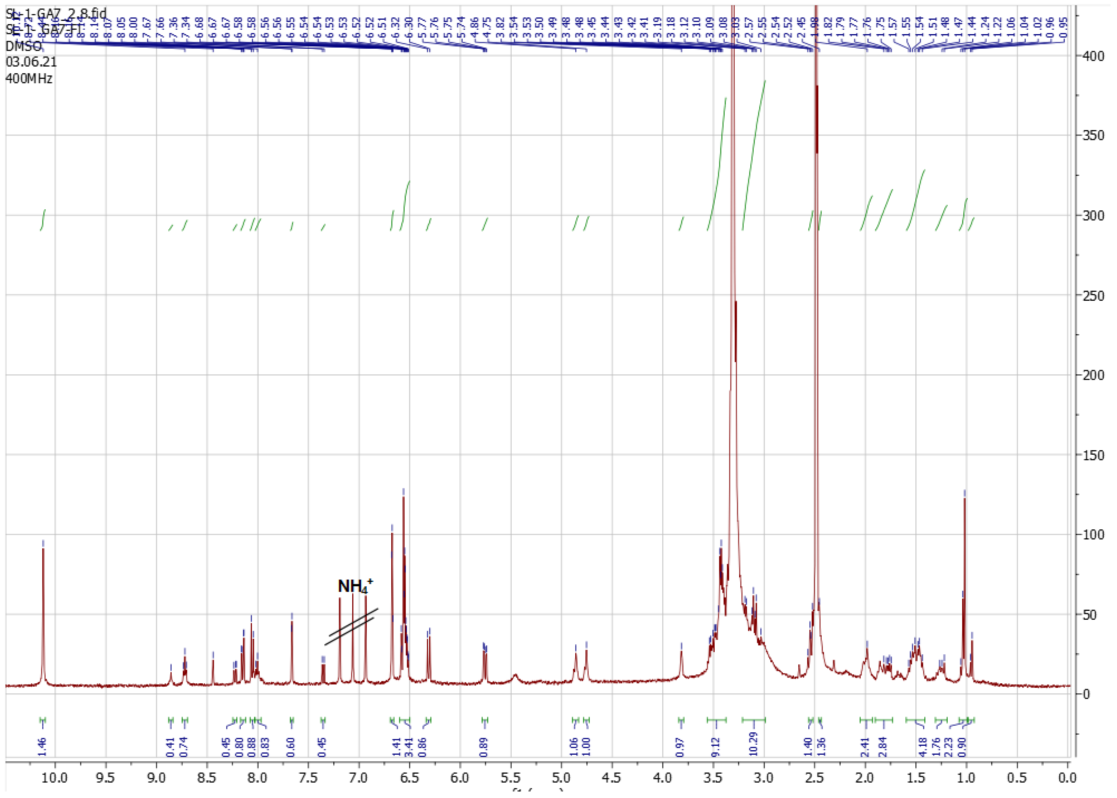

Supplementary Figure 9. $^{1}$H-NMR spectrum of $GA_7$-Fl

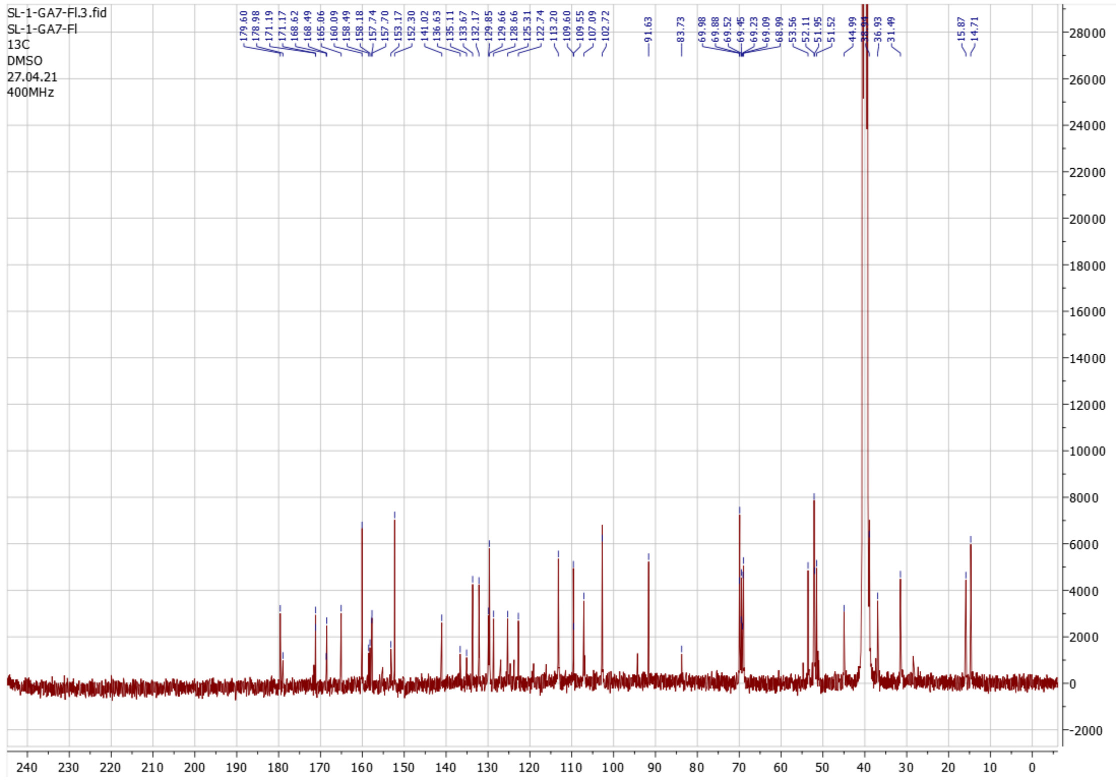

Supplementary Figure 10. $^{13}$C-NMR spectrum of $GA_7$-Fl

1. Shani, E. *et al.* Gibberellins accumulate in the elongating endodermal cells of Arabidopsis root. *PNAS* **110**, 4834–4839 (2013).
