## Supplementary Table 2 for "A quantitative gibberellin signalling biosensor reveals a role for gibberellins in internode specification at the shoot apical meristem"

**Supplementary Table 2. Oligos used in this study.**

| No | Name | Sequence (5’-3’) | Usage |
| --- | --- | --- | --- |
| 1 | RGA.attB1.F | GGGGACAAGTTTGTACAAAAAAGCAGGCTTCATGAAGAGAGATCATCACCAATTCCA | Forward primer synthesizing RGA |
| 2 | RGA.attB2.R | GGGGACCACTTTGTACAAGAAAGCTGGGTTGTACGCCGCCGTCGAGAGTTTCCAAG | Reverse primer synthesizing RGA |
| 3 | RGAm1.2R | AGCCATAAGCGCGTGGACTAAAGCAGCAGCGTTCTCTTGCGAGTCAACC | Reverse primer synthesizing RGA^m1^ |
| 4 | RGAm1.3F | GGTTGACTCGCAAGAGAACGCTGCTGCTTTAGTCCACGCGCTTATGGCT | Forward primer synthesizing RGA ^m1^ |
| 5 | RGAm2.2R | AATCAAACAGAGTCGAAGCAGCAGCTAACGATTCAGTAAAC | Reverse primer synthesizing RGA^m2^ |
| 6 | RGAm2.3F | GTTTACTGAATCGTTAGCTGCTGCTTCGACTCTGTTTGATT | Forward primer synthesizing RGA ^m2^ |
| 7 | RGAm3.attB2.R | GGGGACCACTTTGTACAAGAAAGCTGGGTTGGTGGTAATGAGTGGACGAGTGTGCC | Reverse primer synthesizing RGA ^m3^ |
| 8 | RGAm4.R | GGGGACCACTTTGTACAAGAAAGCTGGGTTGTACGCCGCCGTCGAGAGTTTCCAAGCGTCGGTGG | Reverse primer synthesizing RGA ^m4^ |
| 9 | GID1a.attB1.F | GGGGACAAGTTTGTACAAAAAAGCAGGCTTCATGGCTGCGAGCGATGAAGTTAATCTT | Forward primer synthesizing GID1a |
| 10 | GID1a.attB2.R | GGGGACCACTTTGTACAAGAAAGCTGGGTTACATTCCGCGTTTACAAACGCCGAAA | Reverse primer synthesizing GID1a |
| 11 | M5RGA.attB1.F | GGGGACAAGTTTGTACAAAAAAGCAGGCTTCACGGCGGCGGGTGAGTCAACTCGTTC | Forward primer synthesizing M5RGA |
| 12 | VENUS.attB2.R | GGGGACCACTTTGTACAAGAAAGCTGGGTTGATAGATCTCTTGTACAGCTCGTC | Reverse primer synthesizing VENUS |
| 13 | GID1b.attB1.F | GGGGACAAGTTTGTACAAAAAAGCAGGCTTCATGGCTGGTGGTAACGAAGTCAACCTTAACGAA | Forward primer synthesizing GID1b |
| 14 | GID1b.attB2.R | GGGGACCACTTTGTACAAGAAAGCTGGGTTCTAAGGAGTAAGAAGCACAGGACTTGACTTGCTTT | Reverse primer synthesizing GID1b |
| 15 | GID1c.attB1.F | GGGGACAAGTTTGTACAAAAAAGCAGGCTTCATGGCTGGAAGTGAAGAAGTTAATCTTATTGAG | Forward primer synthesizing GID1c |
| 16 | GID1c.attB2.R | GGGGACCACTTTGTACAAGAAAGCTGGGTTTCATTGGCATTCTGCGTTTACAAATGCAGCTAT | Reverse primer synthesizing GID1c |
| 17 | pUBQ10.F | AAAGTCTGTATATATGACACAGAA | Forward primer synthesizing UBQ10 promoter |
| 18 | pHDG4.F | AAACGCTTTGTCGGTGATCTAAGAA | Forward primer synthesizing HDG4 promoter |
| 19 | pPDF1.F | ATAGCGGAATAGCTGGCAACTTCAA | Forward primer synthesizing PDF1 promoter |
| 20 | GID1a-CDS-F | TTAATATCCAACTTCAAAGTAGCC | Forward primer synthesizing probe of GID1a CDS (969 bp) |
| 21 | GID1a-CDS-R T7 | TAATACGACTCACTATAGGG ACATTCCGCGTTTACAAACG | Reverse primer synthesizing probe of GID1a CDS |
| 22 | GID1a-3UTR-F | CACTGGGTTAGAGAAAGAAG | Forward primer synthesizing probe of GID1a 3’-UTR (557 bp) |
| 23 | GID1a-cDNA-R1914 | TAATACGACTCACTATAGGG GGCTTTTTGAAACACATTATATC | Reverse primer synthesizing probe of GID1a 3’-UTR |
| 24 | GID1b-CDS-F40 | AGAATTGTCCCACTCAACACATGGG | Forward primer synthesizing probe of GID1b CDS (1035 bp) |
| 25 | GID1B-antisenP1 | TAATACGACTCACTATAGGG AGGAGTAAGAAGCACAGG | Reverse primer synthesizing probe of GID1b CDS |
| 26 | GID1B-senP2 | CAACGACAATGTCTGGTCATG | Forward primer synthesizing probe of GID1b 3’-UTR (488 bp) |
| 27 | GID1B-antisenP2 | TAATACGACTCACTATAGGG AATGCATGAACTAAAACAAGAAGG | Reverse primer synthesizing probe of GID1b 3’-UTR |
| 28 | GID1c-CDS-F | CTAATATCCAACTTTAAGCTAGC | Forward primer synthesizing probe of GID1c CDS (966 bp) |
| 29 | GID1c-CDS-R T7 | TAATACGACTCACTATAGGG TTGGCATTCTGCGTTTAC | Reverse primer synthesizing probe of GID1c CDS |
| 30 | GID1c-3UTR-F | GAACACTCTTATCTCTCACTG | Forward primer synthesizing probe of GID1c 3’-UTR (545 bp) |
| 31 | GID1c-cDNA-R1819 | TAATACGACTCACTATAGGG TAAGACTTATACATCAAATCTTTGTC | Reverse primer synthesizing probe of GID1c 3’-UTR |
| 32 | GAI_cDNA-F | TTGAGCTGTAGATGTTGCTGTTAG | Forward primer synthesizing probe of GAI cDNA (1061 bp) |
| 33 | GAI_cDNA-R T7 | TAATACGACTCACTATAGGG AAGTGAGCGAACTTGAGA | Reverse primer synthesizing probe of GAI cDNA |
| 34 | GAI_3UTR-F | TAGATGGTGGCTCAATGAATTG | Forward primer synthesizing probe of GAI 3’-UTR (382 bp) |
| 35 | GAI_3UTR-R T7 | TAATACGACTCACTATAGGG CTGGTCCACCTATAATACCATGC | Reverse primer synthesizing probe of GAI 3’-UTR |
| 36 | RGL1_CDS-F | ATGAAGAGAGAGCACAACCAC | Forward primer synthesizing probe of RGL1 cDNA (1530 bp) |
| 37 | RGL1_CDS-R T7 | TAATACGACTCACTATAGGG CACACGATTGATTCGCC | Reverse primer synthesizing probe of RGL1 cDNA |
| 38 | RGL1_3UTR-F | ATGGGAAAAGTGAAAATGTGC | Forward primer synthesizing probe of RGL1 3’-UTR (310 bp) |
| 39 | RGL1_3UTR-R T7 | TAATACGACTCACTATAGGG TCGTGTCATGAAACATTTATCC | Reverse primer synthesizing probe of RGL1 3’-UTR |
| 40 | RGL2_cDNA-F | GGAGTAAATTTCTTGGAGTTAGG | Forward primer synthesizing probe of RGL2 cDNA (1050 bp) |
| 41 | RGL2_cDNA-R T7 | TAATACGACTCACTATAGGG TGAGCAAAATACGTAGCGAC | Reverse primer synthesizing probe of RGL2 cDNA |
| 42 | RGL2_3UTR-F | CGGTAGAGATGACTCGCC | Forward primer synthesizing probe of RGL2 3’-UTR (258 bp) |
| 43 | RGL2_3UTR-R T7 | TAATACGACTCACTATAGGG CATACAGAATCGTTATCCTTCC | Reverse primer synthesizing probe of RGL2 3’-UTR |
| 44 | RGL3_cDNA-F | GACTATGACAGTCCATTGGACC | Forward primer synthesizing probe of RGL3 cDNA (1101 bp) |
| 45 | RGL3_cDNA-R T7 | TAATACGACTCACTATAGGG ACGTCGTAACAGCTTCTAGAATC | Reverse primer synthesizing probe of RGL3 cDNA |
| 46 | RGL3_3UTR-F | TAGATACGTCGTCATAAAGAGG | Forward primer synthesizing probe of RGL3 3’-UTR (330 bp) |
| 47 | RGL3_3UTR-R T7 | TAATACGACTCACTATAGGG CAAATTCATTCATGCGTCATTTTG | Reverse primer synthesizing probe of RGL3 3’-UTR |
| 48 | GID1b-attB4 | GGGGACAACTTTGTATAGAAAAGTTGGGATTTGTTATGATCTTGTCGGG | Forward primer synthesizing GID1b promoter |
| 49 | GID1b-p3.9k-attB1r | GGGGACTGCTTTTTTGTACAAACTTGA AGTCTCCAAAACCCAGCAAAAAAG | Reverse primer synthesizing GID1b promoter |
| 50 | GID1b-attB2r-2 | GGGGACAGCTTTCTTGTACAAAGTGGATATGGCTGGTGGTAACGAAG | Forward primer synthesizing GID1b cDNA plus terminator |
| 51 | GID1b-attB3 | GGGGACAACTTTGTATAATAAAGTTGG GATTAACATAACCGCTCGTC | Reverse primer synthesizing GID1b cDNA plus terminator |
